# Antibiotics of the future are prone to resistance in Gram-negative pathogens

**DOI:** 10.1101/2023.07.23.550022

**Authors:** Lejla Daruka, Márton Simon Czikkely, Petra Szili, Zoltán Farkas, Dávid Balogh, Gábor Grézal, Elvin Maharramov, Thu-Hien Vu, Levente Sipos, Szilvia Juhász, Anett Dunai, Andreea Daraba, Mónika Számel, Tóbiás Sári, Tamás Stirling, Bálint Márk Vásárhelyi, Eszter Ari, Chryso Christodoulou, Máté Manczinger, Márton Zsolt Enyedi, Gábor Jaksa, Károly Kovács, Stineke van Houte, Elizabeth Pursey, Lajos Pintér, Lajos Haracska, Bálint Kintses, Balázs Papp, Csaba Pál

## Abstract

Despite the ongoing development of new antibiotics, the future evolution of bacterial resistance may render them ineffective. We demonstrate that antibiotic candidates currently under development are as prone to resistance evolution in Gram-negative pathogens as clinically employed antibiotics. Resistance generally stems from both genomic mutations and the transfer of antibiotic resistance genes from microbiomes associated with humans, both factors carrying equal significance. The molecular mechanisms of resistance overlap with those found in commonly used antibiotics. Therefore, these mechanisms are already present in natural populations of pathogens, indicating that resistance can rapidly emerge through selection of pre-existing bacterial variants. However, certain combinations of antibiotics and bacterial strains are less prone to developing resistance, emphasizing the potential of narrow-spectrum antibacterial therapies that could remain effective. Our comprehensive framework allows for predicting future health risks associated with bacterial resistance to new antibiotics.

## Main Text

Multi-drug resistant bacterial infections have been recognized as a major public health concern, and are responsible for a significant proportion of deaths worldwide^1^. Alarmingly, the O’Neill report^2^ estimated that the mortality of drug-resistant bacterial infections might exceed that of malignant tumors by 2050, unless the current trends are reversed. Paradoxically, however, many pharmaceutical companies have discontinued their antibiotic research programs^3^. This is partly due to the rapid spread of multi-drug resistant bacteria, which makes the commercial success of new antimicrobial drugs unpredictable^4,5^.

The danger of wasting considerable time, effort, and funding on antibiotic candidates prone to resistance deter companies from developing new antibiotics. Of many examples, GlaxoSmithKline (GSK) spent 15 million USD to acquire the GSK2251052 molecule and invested considerable further money in its development. However, the project was later canceled due to the rapid emergence of resistance against GSK2251052^6^. Antibiotics launched to the market can also lose utility and revenues in only a few years due to fast emerging resistance. A recent example is dalbavancin, one of the few treatments available to patients infected with methicillin-resistant *S. aureus* (MRSA)^7,8^. Approximately 10 years of research and funding was jeopardized, as resistance emerged after merely two years of commercialization.

Therefore, it is imperative to predict the possible evolutionary routes toward resistance, especially at an early stage of antibacterial drug discovery. However, identification of antibacterial compounds with limited resistance is a complex problem for three main reasons: i) the variety of molecular mechanisms contributing to antimicrobial resistance, ii) the numerous pathogenic bacteria to be considered, and iii) the number of potential antibacterial compounds to be tested. Specifically, bacteria can acquire resistance through diverse genetic mechanisms, including point mutations, amplification of genomic segments, and horizontal transfer of resistance genes^9^. Based on the 2021 WHO pipeline report^10^ and recent reviews on the subject^11–13^, we selected new antimicrobial compounds which have been introduced into clinical practice recently (after 2017) or are currently in development (i.e., ‘recent’ antibiotics, see Table 1). The selected compounds are generally small molecules, directly target Gram-negative bacteria, and are planned to be employed as monotherapies, mostly as intravenous or oral applications, but prior knowledge on de novo emergence of resistance is limited (Supplementary Table 4, for selection criteria, see Supplementary Table 1, and Supplementary Note 1.). Our main goal was to compare the resistance profiles of these ‘recent’ antibiotics to clinically employed antibiotics (i.e., ‘control’). The ‘control’ antibiotics belong to distinct major classes of antibiotics, and all of them have been in clinical usage for many years (Supplementary Table 1). To systematically characterize the bacterial capacity for resistance and the underlying molecular mechanisms, we combined laboratory evolution, functional metagenomic screens, and targeted mutagenesis. To explore potential clinical relevance of our findings, we examined whether the identified resistance mutations and antibiotic resistance genes can be found in natural bacterial isolates and in human associated microbiomes (Fig. 1).

**Fig. 1.**
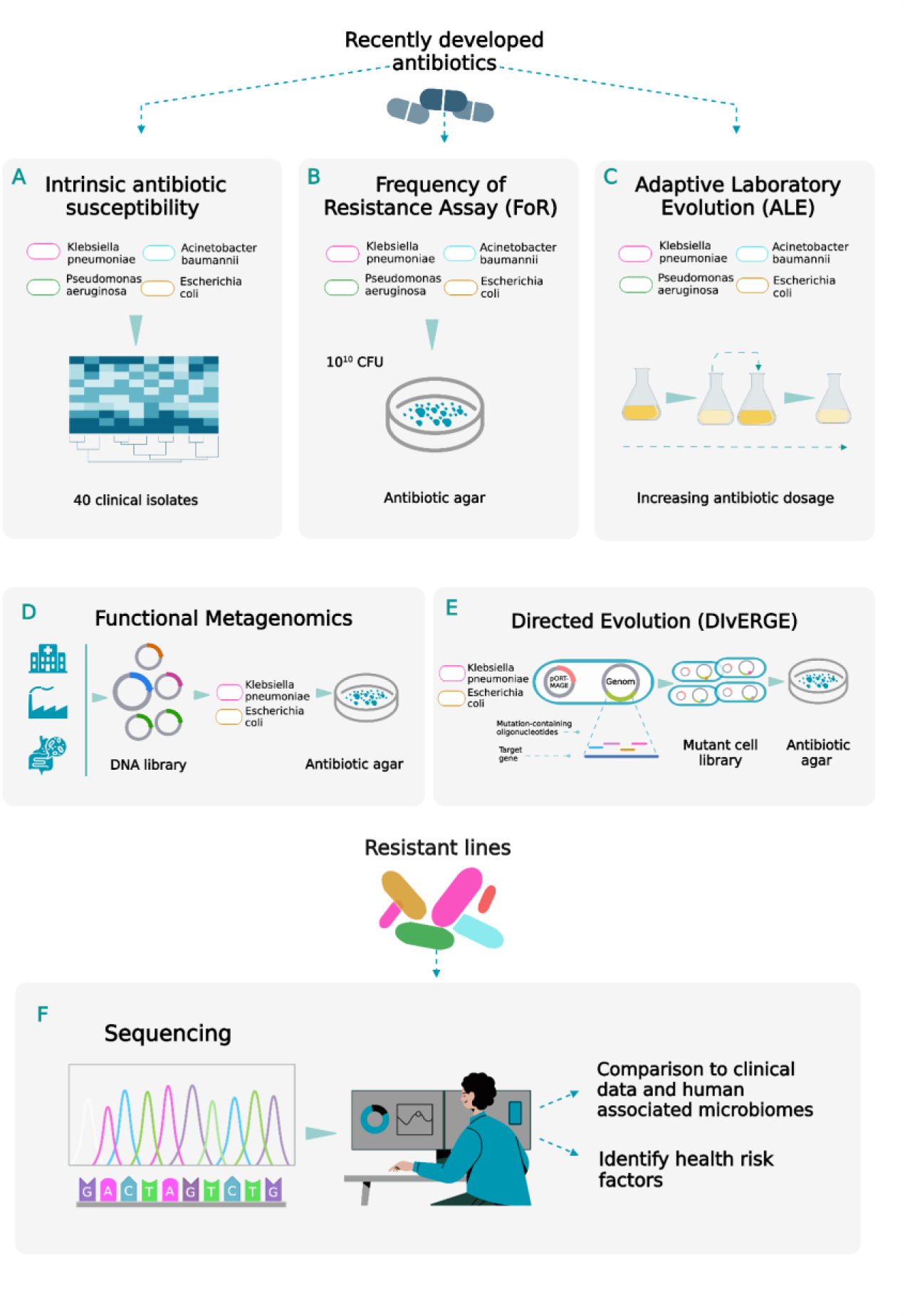
Deep analysis of resistance evolution. Evolution of resistance to 13 recently developed antibiotics was studied. To this aim we selected new antimicrobial compounds which are currently under development or have reached the market recently. To systematically study the possible routes of resistance evolution and the underlying molecular mechanisms, we combined multiple methods shown in gray boxes. We used multiple clinically relevant Gram-negative bacterial species as indicated in each box. **A)** Intrinsic antibiotic susceptibility testing was performed using clinical isolates of *E. coli, K. pneumoniae, A. baumannii* and *P. aeruginosa*. **B-C)** Standard frequency of resistance assays (FoR) with ∼10^10^ cells and adaptive laboratory evolution (ALE) experiments were performed using a sensitive and MDR strain of *E. coli, K. pneumoniae, A. baumannii* and *P. aeruginosa.* **D)** Functional metagenomic screens for antibiotic resistance genes from three metagenomic DNA libraries (soil and gut microbiomes and clinical genomes) were performed in *E. coli* and *K. pneumoniae.* **E)** The directed evolution method called DIvERGE was used in *E. coli* and *K. pneumoniae* to test cross-resistance between recent and control topoisomerase inhibitors. **F)** We sequenced the complete genomes of 457 resistant lines and 690 independent DNA fragments to decipher the underlying molecular mechanisms of resistance and explore the prevalence of these mutations in the genomes of naturally occurring clinical isolates. Additionally, we evaluated whether the detected putative antibiotic resistance genes could pose threats to public health.

**Table 1.** Antibiotics employed in this study. The table shows the antibiotics used in this study, including ‘control’ (blue) and ‘recent’ (orange) antibiotics. Some of the recent antibiotics are derivatives of established antibiotic classes and are supposed to address problems associated with class-specific resistance mechanisms. When it comes to compounds with novel molecular targets (or novel mode of action against a known target), the antibiotic pipeline is thin ^10^. In this group, we considered lead compounds in clinical trials or at least established efficacy against Gram-negative ESKAPE pathogens in mouse infections models. The list includes multi-targeting compounds (indicated by asterisks, *), which are considered to be less prone to resistance ^97,98^. Similarly, compounds that attack essential components of the outer cell membrane have been previously suggested to be immune to bacterial resistance ^11,99^, as potential resistance mutations to these drugs would seriously compromise normal cellular functioning. APR has been utilized in veterinary medicine for more than ten years, its current focus lies in clinical trials for the treatment of systemic Gram-negative infections in human patients. As the evolutionary dynamics of resistance against antibiotic combinations can be very different from that of monotherapies, we study recent advances in adjuvant therapies (e.g. β-lactamase inhibitors) in a separate study. Similarly, Gram-positive specific antibiotic candidates, such as teixobactin, will be studied elsewhere. For more information on antibiotic choice, see Supplementary Note 1.

| Antibiotic | Abbreviation | Antibiotic class | Generation | Date of approval/clinical phase |
| --- | --- | --- | --- | --- |
| Omadacycline | OMA | tetracyclines | recent | 2018 |
| Eravacycline | ERA |  | recent | 2018 |
| Doxycycline | DOX |  | control | 1967 |
| Ceftobiprole | CTO | cephalosporins | recent | 2019 |
| Cefiderocol | CID |  | recent | 2019 |
| Cefepime | CEP |  | control | 1994 |
| Delafloxacin* | DEL | topoisomerase inhibitors | recent | 2017 |
| Gepotidacin* | GEP |  | recent | Phase 3 |
| Zoliflodacin | ZOL |  | recent | Phase 3 |
| Moxifloxacin* | MOX |  | control | 1999 |
| Apramycin | APR | aminoglycosides | recent | Phase 1 |
| Gentamicin | GEN |  | control | 1964 |
| Sulopenem | SUO | carbapenems | recent | Phase 3 |
| Meropenem | MER |  | control | 1996 |
| Tridecaptin M152-P3* | TRD | membrane targetings | recent | pre-clinical |
| POL-7306* | POL |  | recent | pre-clinical |
| SCH79797* | SCH |  | recent | pre-clinical |
| SPR-206 | SPR |  | recent | Phase 1 |
| Polymyxin B | PMB |  | control | 1964 |

## Results

### Overlap in naturally occurring resistance between control and recent antibiotics

We first selected 40 representative strains from four Gram-negative bacterial pathogens, including *Escherichia coli*, *Klebsiella pneumoniae*, *Acinetobacter baumannii* and *Pseudomonas aeruginosa* (Supplementary Table 2), and measured their susceptibilities to 22 ‘control’ and 13 ‘recent’ antibiotics (Supplementary Table 3). Several strains with clinical origins were confirmed to be extensively drug resistant (XDR), as the MICs of nearly all clinically recommended antibiotics were above the established clinical breakpoints (Extended Data Fig. 1, Extended Data Fig. 2, Extended Data Fig. 3). On this panel of bacterial strains, cefiderocol, SPR-206, eravacycline and delafloxacin have on average significantly higher efficacy (i.e., lower average MIC) compared to their control counterparts with similar modes of action (Fig. 2A). Indeed, hierarchical clustering based on the heatmap of antibiotic susceptibility profiles have revealed that ‘control’ and ‘recent’ antibiotics with related modes of action cluster together (Fig. 2B). Moreover, MDR and XDR bacterial strains generally displayed reduced susceptibilities to both control and recent antibiotics, compared to antibiotic sensitive (SEN) strains belonging to the same species (Fig. 2C). Together, these results indicate an overlap in resistance mechanisms between the two antibiotic groups. Notably, certain recent membrane targeting antibiotics, such as POL-7306 and SPR-206 were as effective against MDR/XDR strains, as against sensitive strains (Extended Data Fig. 4), highlighting their promising antibacterial activity.

**Fig. 2A.**
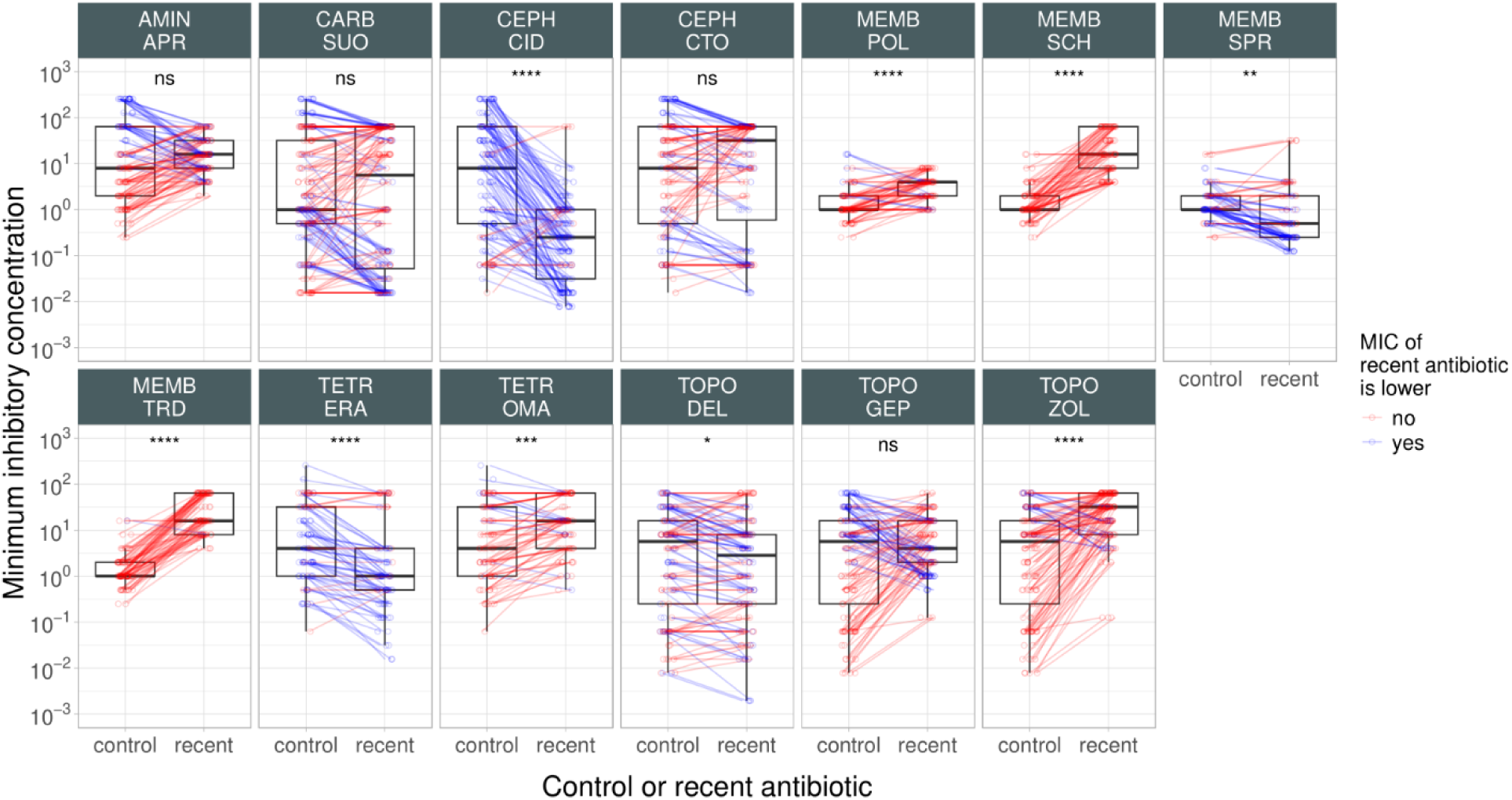
Comparison of minimum inhibitory concentration for control and recent antibiotics. The figure shows the MIC values (on a log10 scale) of control and recent antibiotics across all tested bacterial strains. Each panel represents a specific recent antibiotic, as indicated at the top with the class it belongs to. Individual points depict strain-antibiotic pairs, with lines connecting paired data points representing MIC values of the specific recent and one within-class control antibiotic for the same strain. Blue and red colors indicate cases where the MIC of a recent antibiotic is lower or not than that of the control antibiotic for the same strain. Boxplots display the median, first, and third quartiles, with whiskers indicating the 5th and 95th percentiles. Paired Wilcoxon rank sum analysis was performed to assess significant difference between control and recent antibiotics within each class. ****/***/**/* indicate P < 0.0001/0.001/0.01/0.05, ns indicates that the P value is not significant. For antibiotic abbreviations, see Table 1. Antibiotic classes: TOPO (topoisomerase inhibitors), TETR (tetracyclines), AMIN (aminoglycosides), CARB (carbapenems), CEPH (cephalosporins), and MEMB (membrane-targeting antibiotics.

**Fig. 2B.**
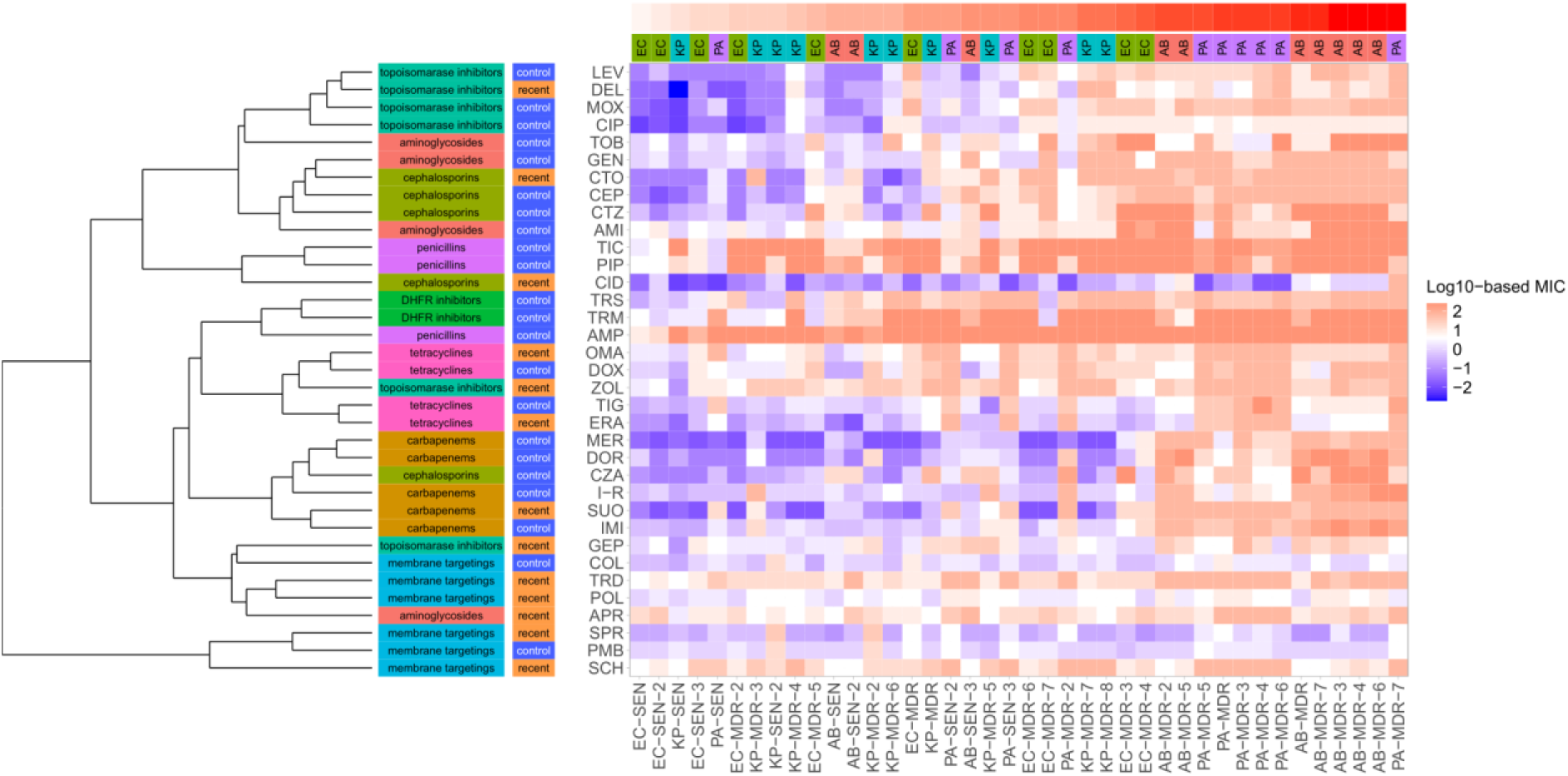
The heatmap shows the antibiotic susceptibility profiles of bacterial strains belonging to four bacterial species, including *Escherichia coli* (EC), *Klebsiella pneumoniae* (KP), *Acinetobacter baumannii* (AB) and *Pseudomonas aeruginosa* (PA). The antibiotic panel consists of 22 ‘control’ (blue) and 13 ‘recent’ (orange) antibiotics. Antibiotic generations and classes are indicated on the left panels. Antibiotic clustering was based on calculating Spearman’s correlation of MIC values and using the complete hierarchical clustering method. The bacterial strains are ordered by the fraction of ‘control’ antibiotics (top panel) they are resistant to (defined by the corresponding species-specific clinical breakpoint values). For abbreviations, see Table 1 and Supplementary Table 2.

**Fig. 2C.**
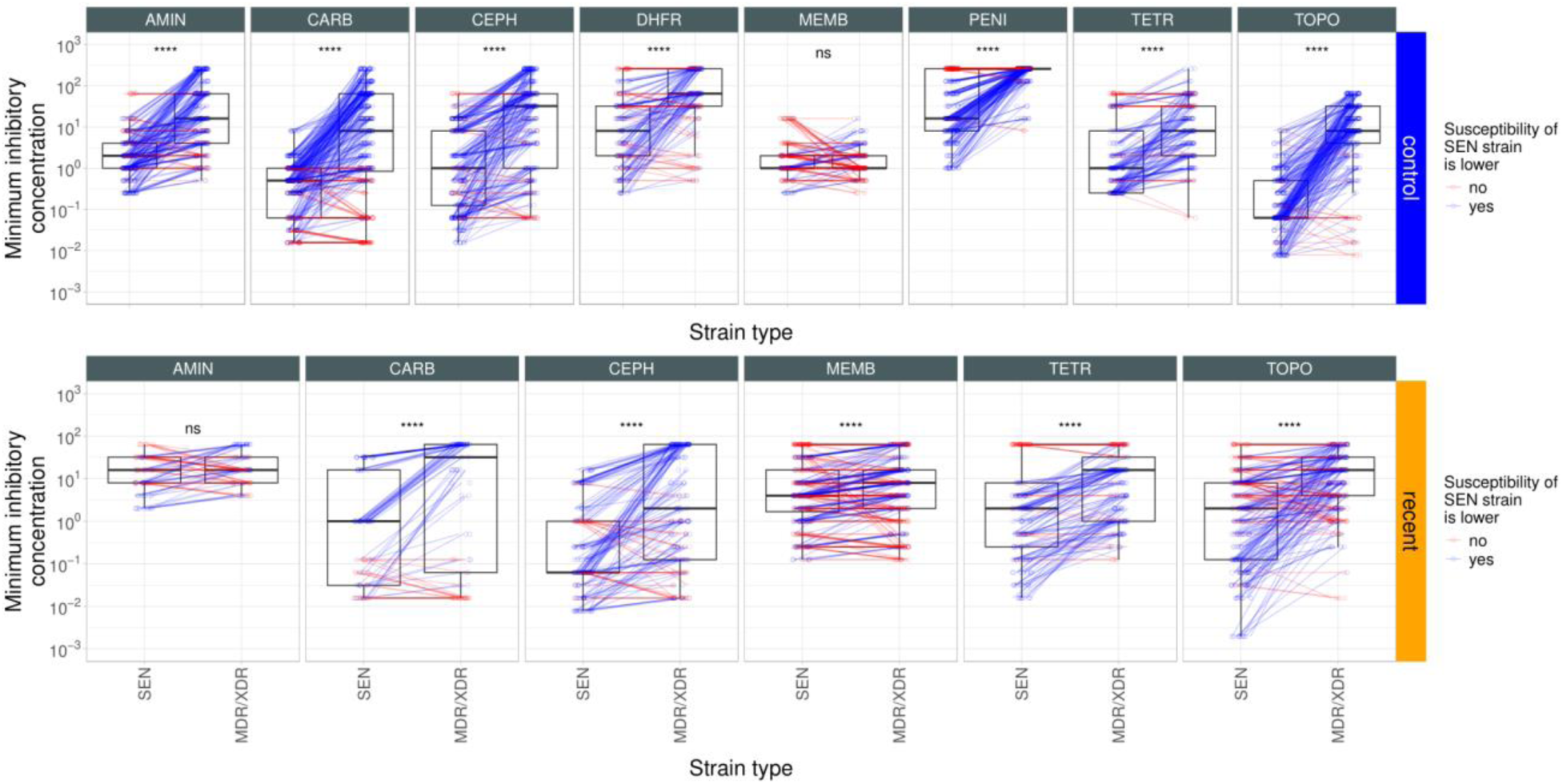
Comparison of minimum inhibitory concentration (MIC) of antibiotics for sensitive and multidrug-resistant/extensively drug-resistant strains. The figure shows the MIC values (on a log10 scale) of control (top row) and recent (bottom row) antibiotics across all tested bacterial strains. Each vertical panel represents a specific antibiotic class, as indicated at the top. Individual points depict strain-antibiotic pairs, with lines connecting paired data points representing MIC values of one antibiotic for one sensitive (SEN) and one multidrug-resistant/extensively drug-resistant (MDR/XDR) strain for the same species. Blue and red colors indicate cases where the MIC of a single antibiotic for a SEN strain is lower or not than that of the MDR/XDR strain for the same species. Boxplots display the median, first, and third quartiles, with whiskers indicating the 5th and 95th percentiles. Paired Wilcoxon rank sum analysis was performed to assess significant difference in sensitivity between antibiotic sensitive (SEN) and MDR/XDR bacterial strains **** indicates P < 0.0001, whereas ns indicates that the P value is not significant. For antibiotic abbreviations, see Table 1. Antibiotic classes: TOPO (topoisomerase inhibitors), TETR (tetracyclines), AMIN (aminoglycosides), CARB (carbapenems), CEPH (cephalosporins), and MEMB (membrane-targeting antibiotics.

### Rapid species-specific evolution of resistance in the laboratory

Next, we asked whether resistance can readily evolve in bacterial pathogens, rendering these compounds less effective in the long-term. Prior laboratory evolution studies generally focused on only a single species, and frequently on non-pathogens. Here we selected one MDR and one sensitive strain of *E. coli*, *K. pneumoniae*, *A. baumannii* and *P. aeruginosa*, respectively (Supplementary Table 2). 32% of all antibiotic-strain combinations were excluded from the analysis due to relatively low initial antibiotic susceptibility (i.e., MIC > 4 µg/ml) (Supplementary Table 3).

To explore first-step resistance, we used a standard protocol for spontaneous frequency-of-resistance analysis (FoR assay)^14–17^ at multiple concentrations of each antibiotic. Approximately 10^10^ bacterial cells were exposed to one of each antibiotic on agar plates for two days at concentrations where the given strain is susceptible (Supplementary Table 3). Mutants with decreased antibiotic susceptibility were detected in 49.8% of the populations (i.e. at least four-fold MIC-fold change). Although clinical breakpoint is generally unknown, recommended intravenous usage is available for most of the studied ‘recent’ antibiotics. Therefore, data on the highest available peak plasma concentration of the drug (measured at intravenous administration) were used as a proxy to estimate potential clinical relevance of the MIC changes in the mutants (Supplementary Table 1). In spite of the short time-scale of the screens, the MICs were either equal to or above the peak plasma concentration in as high as 18.7% of the mutant lines (Fig. 3A and Supplementary Table 5). In addition, for 30% of the FoR adapted lines, the MIC surpassed the clinical breakpoint, where such data was available (Extended Data Fig. 5). On average, ’recent’ and ’control’ antibiotics were equally prone to bacterial resistance, as neither the frequency of appearance per generation of mutants (Extended Data Fig. 6, Wilcoxon rank-sum test, P = 0.9), nor the fold-change in resistance was statistically different (Extended Data Fig. 7A, paired T-test, P = 0.68).

**Fig. 3A.**
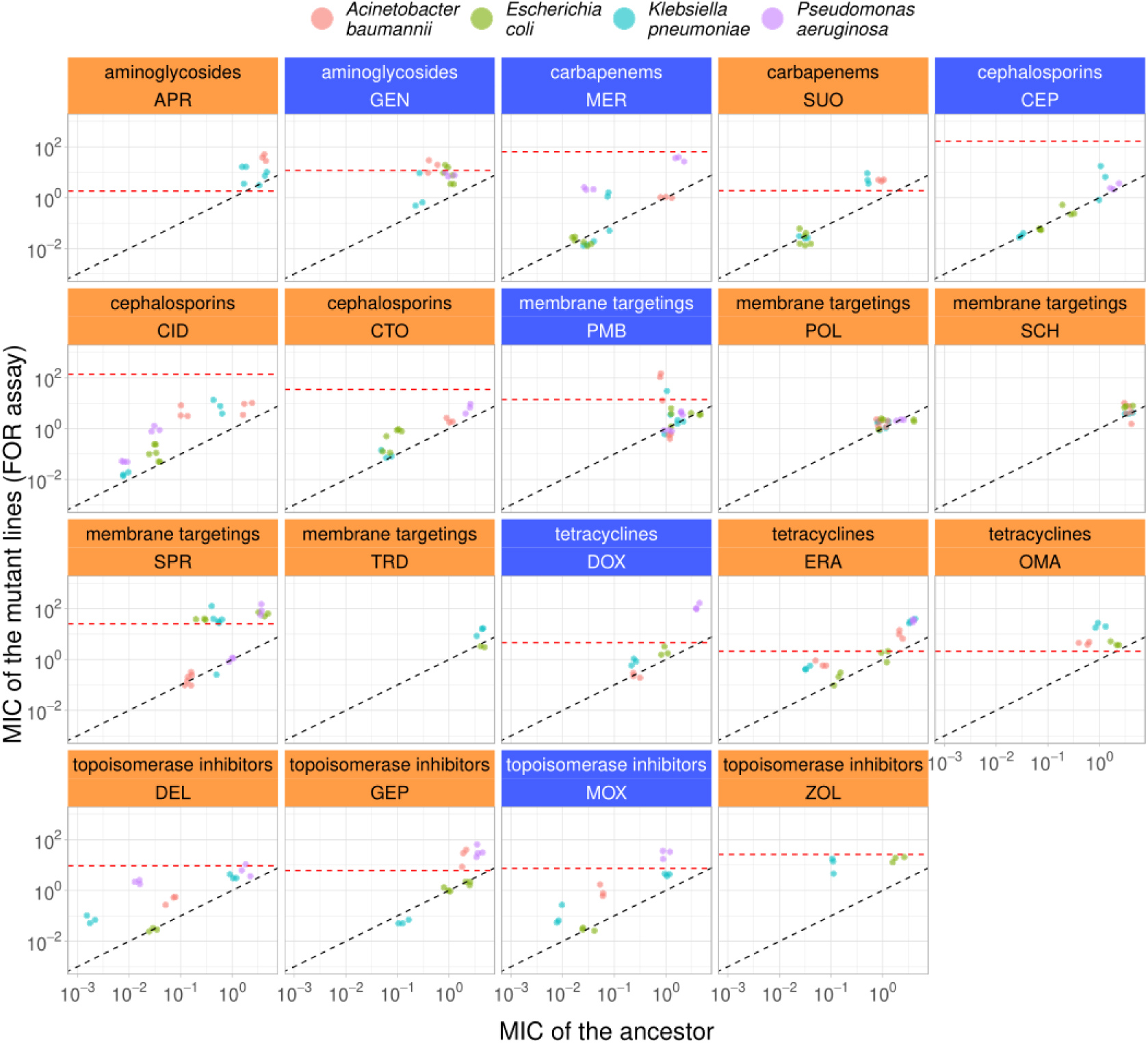
Changes in minimum inhibitory concentrations (MIC) in mutants isolated in frequency-of-resistance (FoR) assays. Each point represents the MICs of a mutant line and the corresponding ancestor (log10 scale). Control and recent antibiotics are indicated by blue and orange panels, respectively. The color of the data points represents the bacterial species. The black dashed line indicated y = x (i.e. no changes in MIC in the mutants), whereas the red dashed line shows the antibiotic specific peak plasma concentration (Supplementary Table 1). For abbreviations, see Table 1.

The above data demonstrates that clinically relevant resistance to ‘recent’ antibiotics frequently arises in a matter of days. However, the underlying FoR assays cannot detect very rare mutations and combinations thereof^18^, and hence underestimate bacterial potential for resistance. Using the same set of eight starting strain (Supplementary Table 2), we next initiated adaptive laboratory evolution (ALE) with the aim to maximize the level of antibiotic resistance in the populations achieved during a longer, but fixed time period (for up to ∼120 generations, see methods). 10 parallel evolving populations were exposed to increasing concentrations of one of each ‘recent’ or ‘control’ antibiotics. The level of resistance was estimated by comparing the MICs of the evolved lines and the corresponding ancestor strains (Fig. 3B).

**Fig. 3B.**
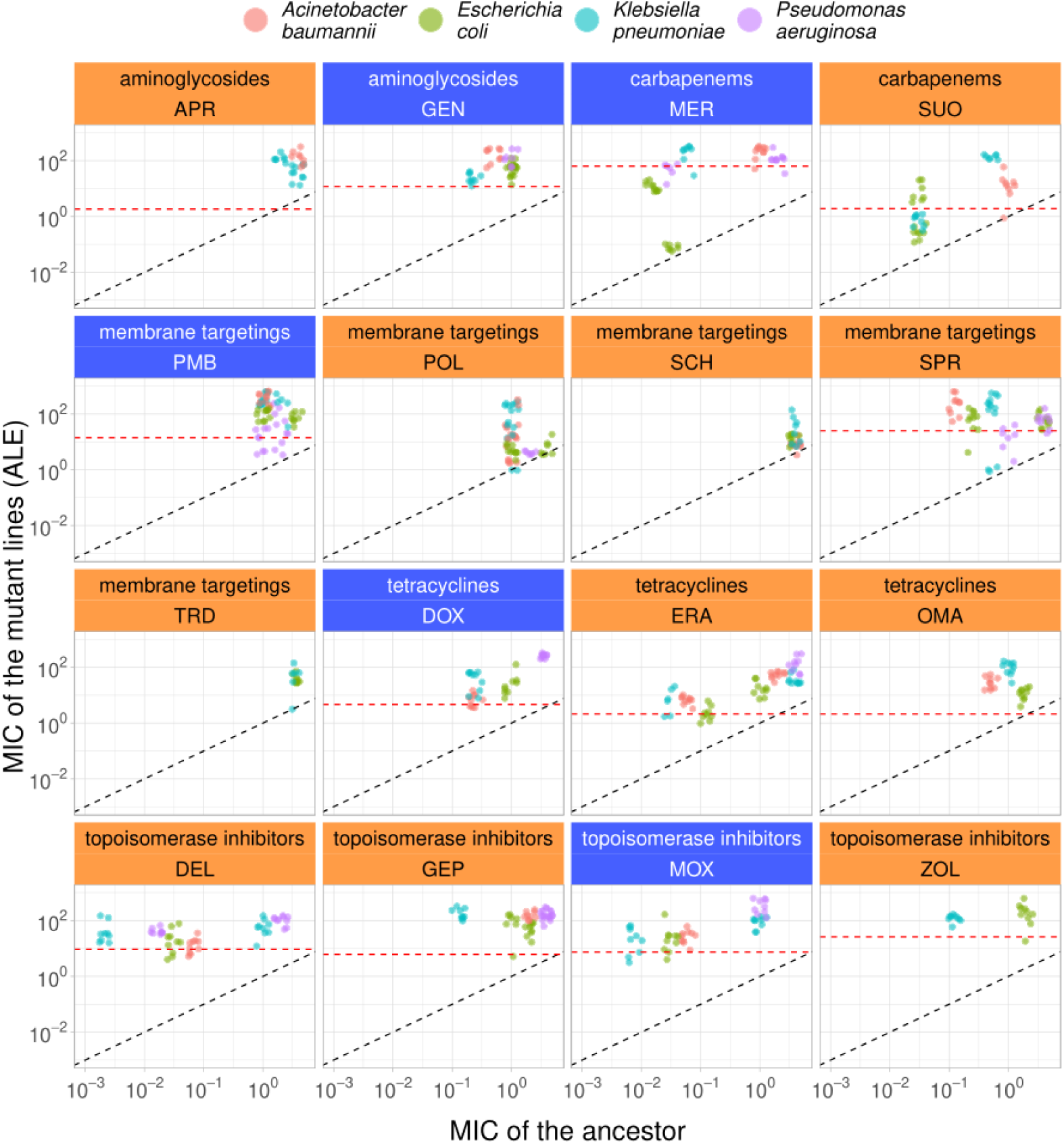
Changes in minimum inhibitory concentrations (MIC) after adaptive laboratory evolution (ALE). Each point represents the MICs of a laboratory evolved line and the corresponding ancestor (log10 scale). ‘Control’ and ‘recent’ antibiotics are indicated by blue and orange panels, respectively. The color of the data points represents the bacterial species. The black dashed line indicated y = x (i.e. no changes in MIC during the course of laboratory evolution), whereas the red dashed line shows the antibiotic specific peak plasma concentration. For abbreviations, see Table 1. Due to low stability in the liquid laboratory medium employed, cephaslosoporin antibiotics were not subjected to ALE.

In general, the evolution of resistance was a rapid process (Supplementary Table 5); the median antibiotic resistance level in the evolved lines was ∼64 times higher compared to the ancestor. The MIC levels were either equal to or above the peak plasma concentration in 87% of all studied populations, indicating their potential clinical relevance. Moreover, for 88.3% of the ALE adapted lines, the MIC surpassed the clinical breakpoint, where such data was available (Extended Data Fig. 5). On average, ’recent’ and ’control’ antibiotics were equally prone to bacterial resistance (Extended Data Fig. 7B, paired T-test, P = 0.37). Alarmingly, high levels of resistance readily emerged against ‘recent’ antibiotics (Fig. 3B) with potent antibacterial activities against a wide-range of MDR and XDR clinical isolates (Extended Data Fig. 4).

### Species-specific potential for resistance evolution to new antibiotics

There was large heterogeneity in the relative level of resistance (i.e., MIC fold-change) across antibiotic-strain combinations, spanning a 65 000-fold range between the observed minimum and maximum values (Fig. 3C). In this section, we explore the key features underlying this variation in the capacity to evolve resistance. The first important issue is whether the bacterium’s initial antibiotic susceptibility reliably forecasts the drug’s long-term efficacy. We found a significant positive correlation between the initial MIC and the increase in resistance level across antibiotics in lines derived from adaptive laboratory evolution in 2 out of 4 species (Extended Data Fig. 8A) and in 5 out of the 8 bacterial strains studied (Extended Data Fig. 8B). We also asked whether initial MIC correlates with the increase in resistance level across strains when each antibiotic is analyzed separately. We found a significant positive trend in 5 out of the 16 antibiotics (Extended Data Fig 9). Overall, these results indicate that the initial MIC is predictive of the antibiotics’ long-term efficacy in a strain- and antibiotic-specific manner.

**Fig. 3C.**
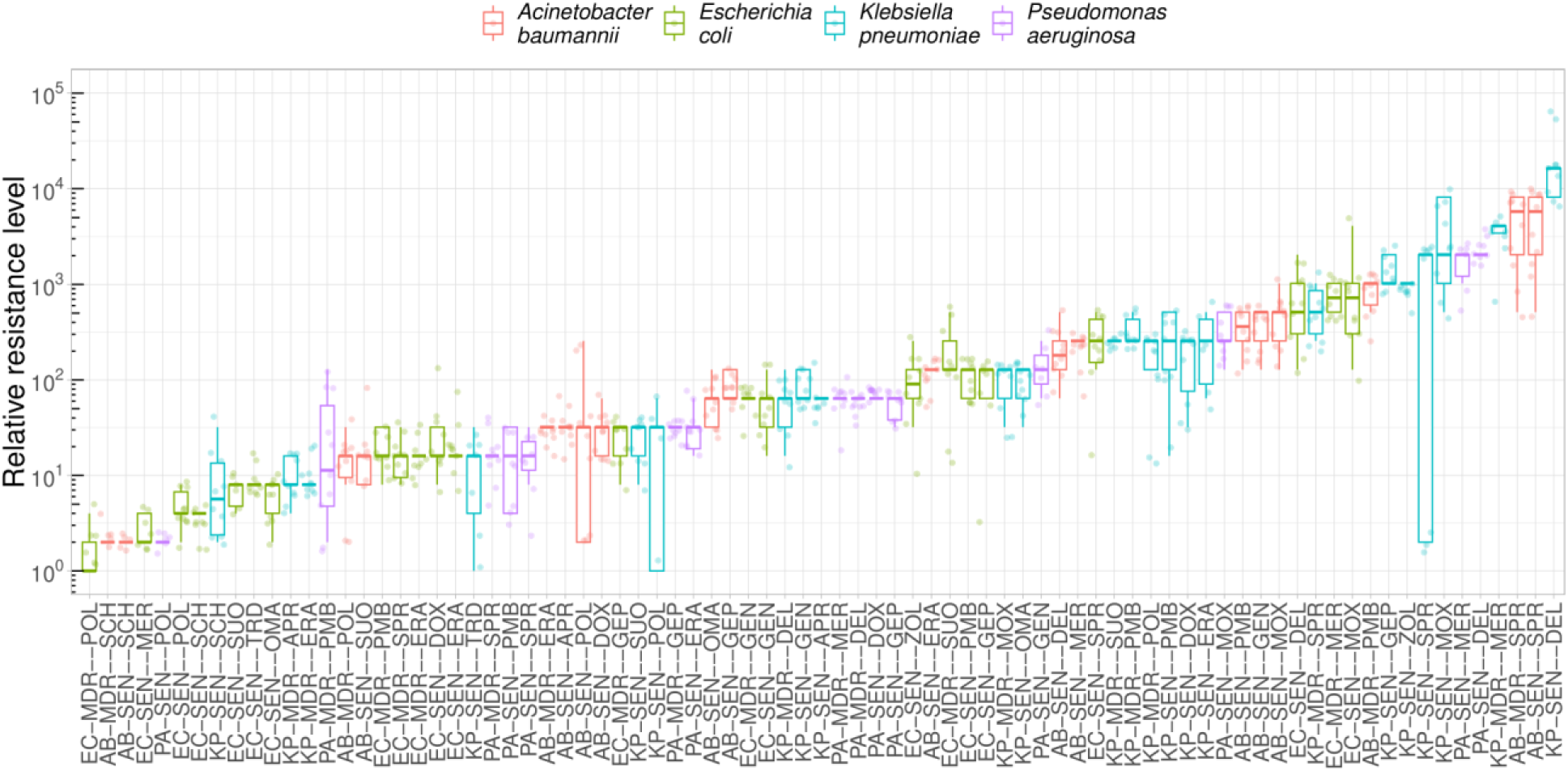
Relative MIC of laboratory evolved lines across all antibiotic-ancestor strain combinations. Relative MIC is the MIC of the evolved line divided by the MIC of the corresponding ancestor. Each point is a laboratory evolved line from adaptive laboratory evolution (ALE), the colors indicate the bacterial species. The boxplots show the median, first and third quartiles, with whiskers showing the 5th and 95th percentiles. There is a highly significant heterogeneity in relative MIC across antibiotic-strain combinations (Kruskal-Wallis chi-squared = 630.43, df = 80, P < 2.2e-16). For antibiotic and bacterial strain abbreviations, see Table 1 and Supplementary Table 2.

Prior works^19,20^ indicate that certain antibiotics are more susceptible to resistance evolution for particular bacterial strains and species compared to others. Accordingly, we used multiple linear regression to investigate the global influence of both antibiotic and the strain’s genetic background on the increase of resistance level, while also considering the resistance level of the ancestor (see Fig. 3D). When examining these factors separately, the antibiotic and strain genetic background explained 24.4% and 8.9% of the variation in the increase of resistance levels, respectively, with the initial antibiotic susceptibility level (MIC) contributing 0.9% to the variance. In an additive model, combining the antibiotic, the initial MIC and genetic background explained approximately 33% of the variation. Importantly, a model that allows an interaction term between genetic background and antibiotic combination explains an additional ∼26% of variation in the increase of resistance, in comparison to the simple additive model (i.e. 58.6% vs 32.6%, see Fig. 3D).

**Fig. 3D.**
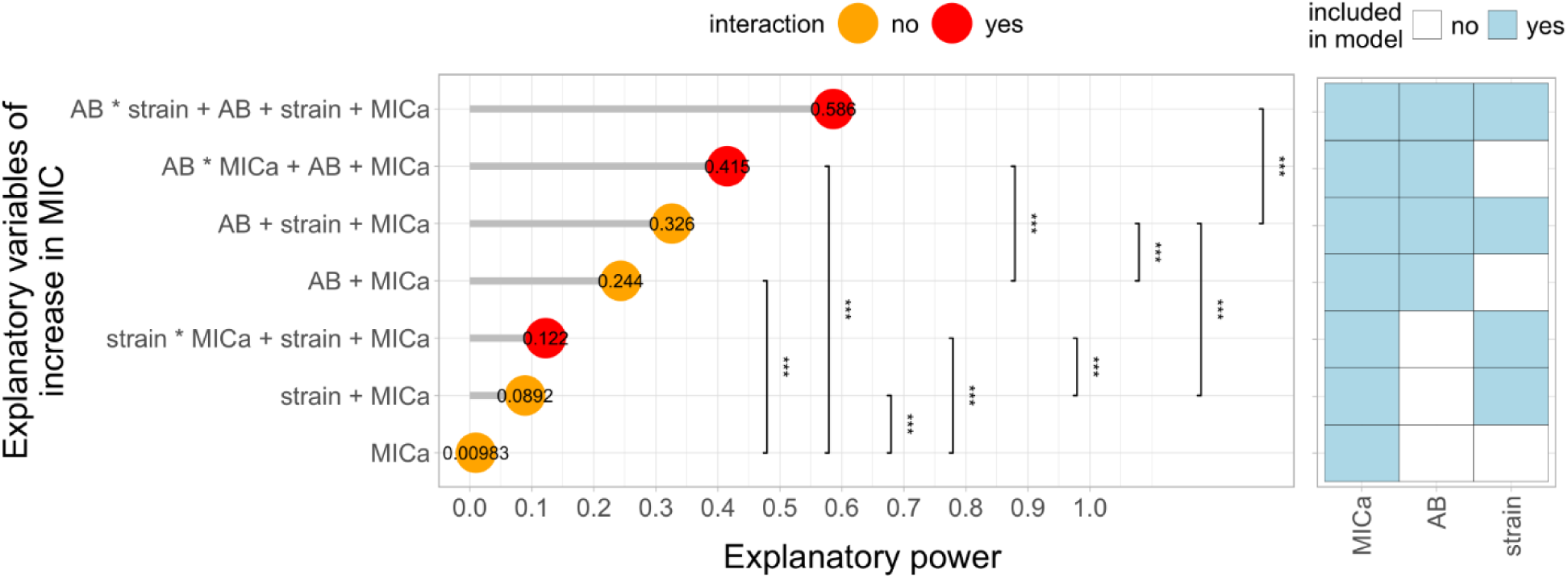
Multiple linear regression (MLR) analysis on features shaping the resistance level reached during evolution. The analysis focused on three main features i) the MIC level of the ancestor strain (MICa), ii) the antibiotic employed (AB) and the iii) genetic background (strain). The adjusted coefficient of determination (adjusted R-square) was used as a statistical metric to measure the explanatory power of the models, ie. how much of the variation in the absolute increase in MIC (log2) can be explained by the variation in these features and combinations thereof, while adjusting for the number of parameters used in the fitted model. Additivity (indicated with + sign in axis labels) and interaction (* sign in axis labels) between explanatory variables are marked with orange and red colors, respectively. The predictors included in the models are also depicted in the right panel. We found a significant increase in adjusted R-square in all cases when a more complex model was compared to a simpler one (ANOVA, P < 0.0001).

Together, these results indicate that the initial genetic makeup of the bacterial population has a large impact on resistance evolution, but predominantly in an antibiotic-specific manner (Extended Data Fig. 10). Two antibiotics, SCH-79797 and SPR-206, highlight this point.

It has been suggested that SCH-79797, a dual-targeting antibiotic, effectively kills a wide range of bacterial pathogens without detectable resistance^21^. By studying this issue in three bacterial pathogens susceptible to SCH-79797, we found high variability in resistance evolution across bacterial species. In particular, an up to 4-fold MIC change was observed in *A. baumannii*, while the same figure was up to 32-fold in *K. pneumoniae* (Extended Data Fig. 11). It is unlikely that the evolution of SCH-79797 resistance would be limited by the mutational supply rate in this species (see Supplementary Note 2.).

Spero Therapeutics (Cambridge, Massachusetts, USA) is currently developing SPR-206 as an innovative treatment option for MDR Gram-negative bacterial infections in hospital settings. SPR- 206 is a direct-acting antibiotic that displayed antibiotic activity against MDR Gram-negative pathogens in preclinical studies^22^. 60 days of laboratory evolution resulted in an up to 8192-fold increment in resistance level to SPR-206 in *A. baumannii*, *K. pneumoniae* and *E. coli* (Extended Data Fig. 11). This raises concerns regarding the clinical application of SPR-206 in these species. In *P. aeruginosa,* an up to 32-fold increment in SPR-206 resistance level was observed (Extended Data Fig. 11). These contrasting results are even more surprising, as the initial SPR-206 susceptibilities show only very limited variation (MIC ranging from 0.125 to 1 µg/ml) across these four bacterial species.

Together, these results indicate that specific bacterial strains and species can evolve higher levels of resistance than others to certain antibiotics.

### Overlap in mutational profiles across antibiotic treatments

To identify mutational events underlying resistance, resistant lines derived from laboratory evolution (n=381) and frequency-of-resistance (n=135) assays were subjected to whole-genome sequencing (Supplementary Table 6). We implemented an established computational pipeline to identify mutations relative to the corresponding ancestral genomes. 10 evolved lines accumulated exceptionally large numbers of mutations, many of which are likely to be functionally irrelevant (Extended Data Fig. 12). These lines exert elevated genomic mutation rates, and indeed 6 out of these 10 lines carry mutations in methyl-directed mismatch repair (*mutS, mutL* or *mutY*). Such mutator bacteria frequently arise in response to antibiotic stress in clinical and laboratory settings^23^. For the remaining 506 lines, we identified 1817 unique mutational events, including 1212 single nucleotide polymorphisms (SNPs) and 605 insertions or deletions (indels) (Extended Data Fig. 13). We found a significant excess of non-synonymous over synonymous mutations, indicating that the accumulation of the SNPs in protein-coding regions was largely driven by selection towards increased resistance (Extended Data Fig. 14). 19.7% of the observed mutations generated in-frame stop codons or frameshifts or disruption of the start codon, which were most likely to yield result loss-of protein function (Extended Data Fig. 15). This result is consistent with prior studies on the role of inactivating mutations in antibiotic resistance^24^.

In total, 604 mutated protein coding genes were detected, of which 193 mutated in at least two independently evolved lines per species (Supplementary Table 7). Remarkably, 69.4% of all parallel mutated genes carried mutation in lines adapted to different antibiotics (Supplementary Table 7). These results indicate that despite substantial differences in antibiotic treatments, there is a significant overlap in the set of mutated genes (Fig. 4A, Fig. 4B, Supplementary Table 7).

**Fig. 4A.**
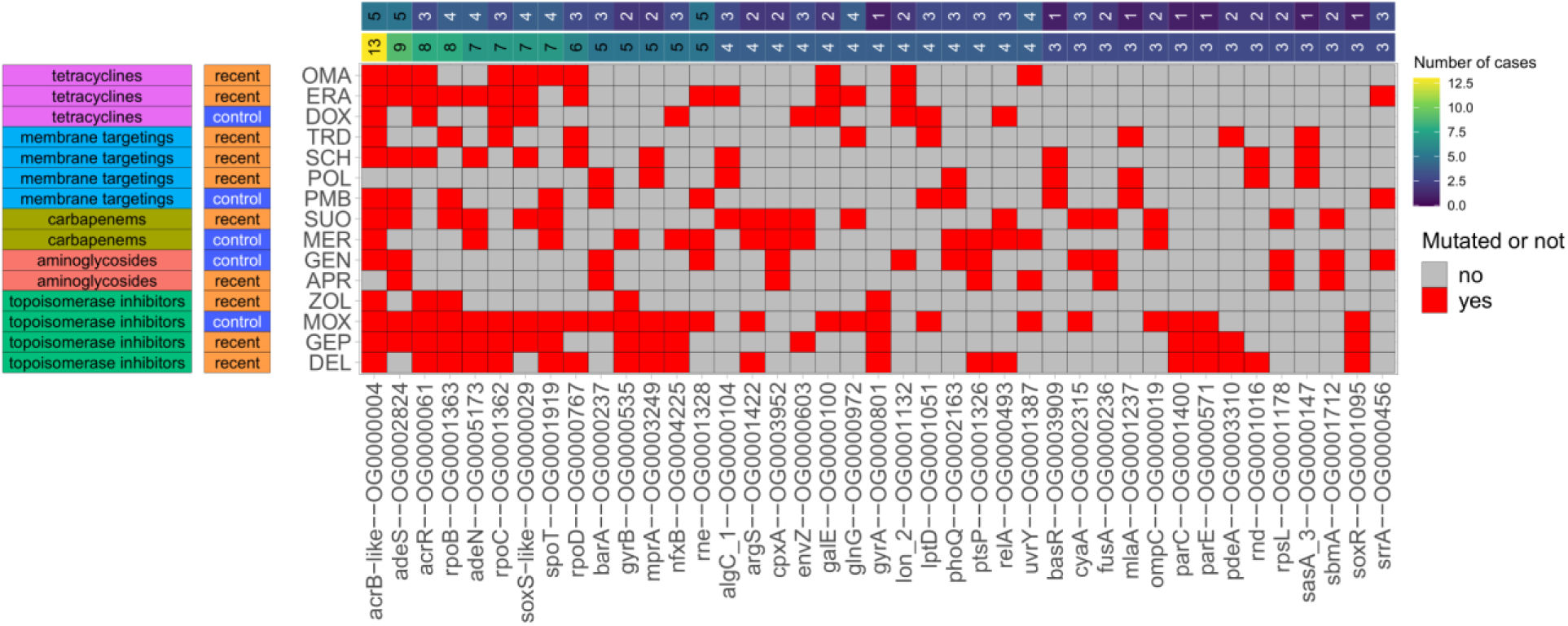
Repeatedly mutated genes during laboratory evolution across different antibiotics. The heatmap shows commonly mutated genes (and the corresponding orthogroups) across the tested antibiotics after adaptive laboratory evolution (ALE). Genes were considered commonly mutated if they accumulated non-synonymous mutations in response to at least three different antibiotic treatments. Panels on the left denote the class and the generation of the given antibiotics. Panels from top to bottom correspond to the number of antibiotic classes and the number of antibiotics a given orthogroup is mutated against. For further details, see Supplementary Table 7.

**Fig. 4B.**
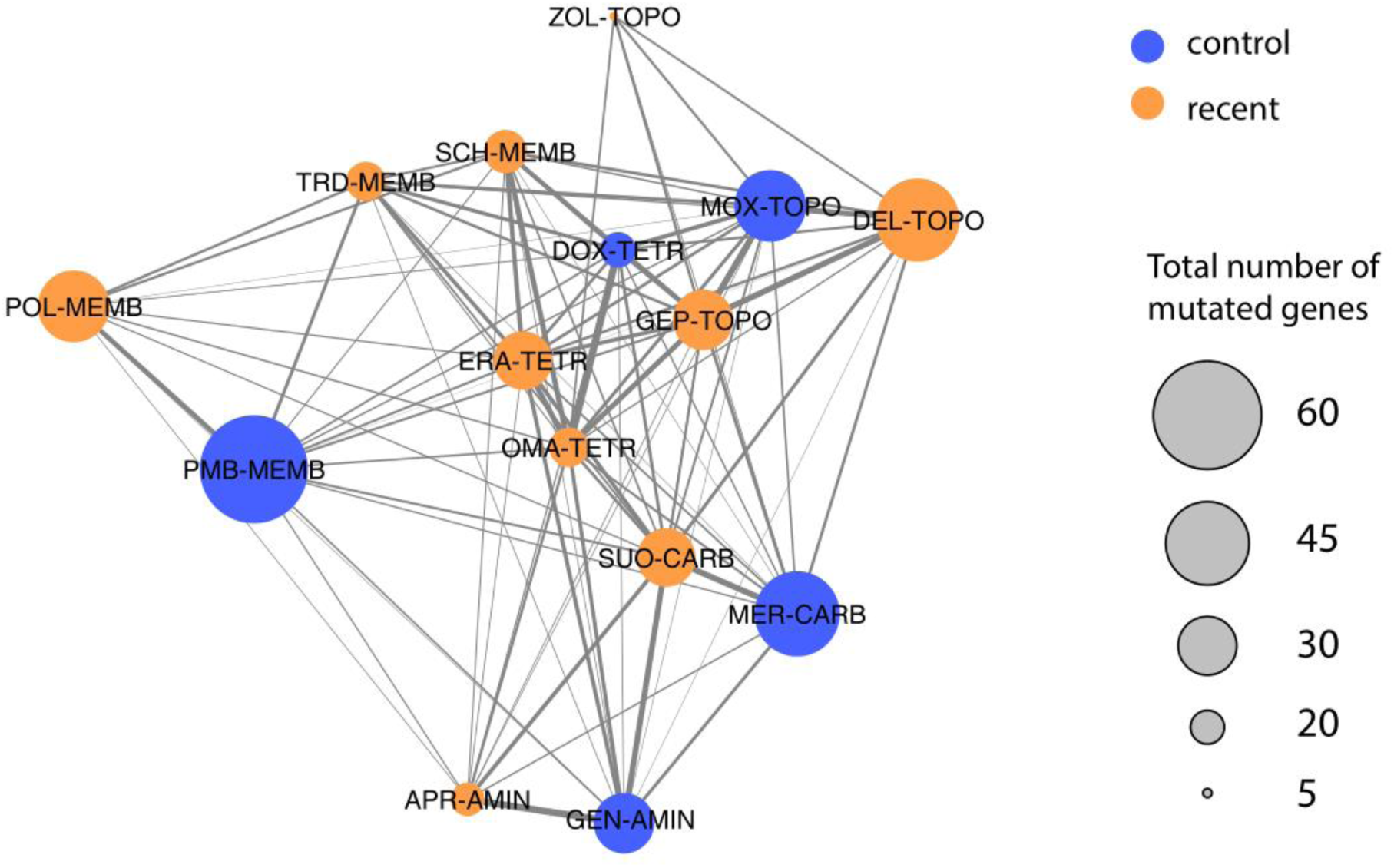
Mutation profile similarity across antibiotics. Each node represents a recent (orange) or control (blue) antibiotic. Links indicate an overlap in the set of mutated genes (or corresponding orthogroups) detected after adaptive laboratory evolution (ALE). Only non-synonymous mutations in protein coding genes were considered. The thickness of the links indicates the extent of overlap (calculated by Jaccard similarity, as earlier^100^) between antibiotic treatments. Abbreviations: TOPO: topoisomerase inhibitors, TETR: tetracyclines, AMIN: aminoglycosides, CARB: carbapenems, CEPH: cephalosporins, MEMB: membrane targeting antibiotics.

An important issue is the extent to which adaptation during the course of laboratory evolution was driven by the physical conditions, including the medium unrelated to antibiotic selection. Several independent lines of evidence indicate that antibiotic selection was the dominant force in our experiments. First, the overlap in the set of mutated genes was especially high when antibiotics with related modes of action were considered, in comparison to antibiotics belonging to different classes (Wilcoxon two-sample test, P < 0.0001). Second, many of the genes detected in multiple antibiotic screens are involved in multidrug resistance (for examples, see Supplementary Table 8). Third, we initiated laboratory evolution experiments intending to identify mutations that provide adaptation to the physical conditions in all four bacterial species (Supplementary Note 3.). The same experimental settings, protocols, and 8 bacterial strains/species were employed, but no antibiotic was added to the medium. Reassuringly, we found that the extent of overlap of molecular mechanisms underlying adaptation in medium-adapted and antibiotic-adapted lines was minimal.

Together, the results indicate that adaptation during the course of laboratory evolution was largely unrelated to the environmental medium employed.

### Resistance mutations to new antibiotics are present in nature

The above results indicate an overlap in the molecular mechanisms of resistance between clinically employed antibiotics and new antibiotic candidates (Fig. 4A, Fig. 4B). Therefore, we hypothesized that the laboratory observed mutations may be already present in environmental and clinical bacterial isolates, indicating their potential clinical relevance.

To investigate this hypothesis, we analyzed the prevalence of the observed mutations from laboratory evolved *E. coli* and *A. baumannii* lines in a catalog of genomes derived from natural isolates of *E. coli* (n=20,786) and *A. baumannii* (n=15,185) (Fig. 4C). For simplicity, the analysis focused on non-synonymous mutations in protein coding sequences, and estimated their frequencies in the genomes of environmental isolates in these two species. In case of *E. coli*, as high as 31.4% of the 245 laboratory observed non-synonymous mutations were identified in at least one of the investigated genomes, while the same figure is 27.3% of 216 mutations for *A. baumannii*. While the majority of the mutations found in *E. coli* were found to be relatively rare (i.e., typically found in less than 1% of the isolates, Supplementary Table 9), they were specifically enriched among pathogenic isolates than in other natural isolates (Fisher’s test, P < 2.2×10-16, odds ratio = 3.16). Several adaptive mutations were equal or even more abundant than canonical antibiotic resistance mutations in clinical isolates (Extended Data Fig. 16).

**Fig. 4C.**
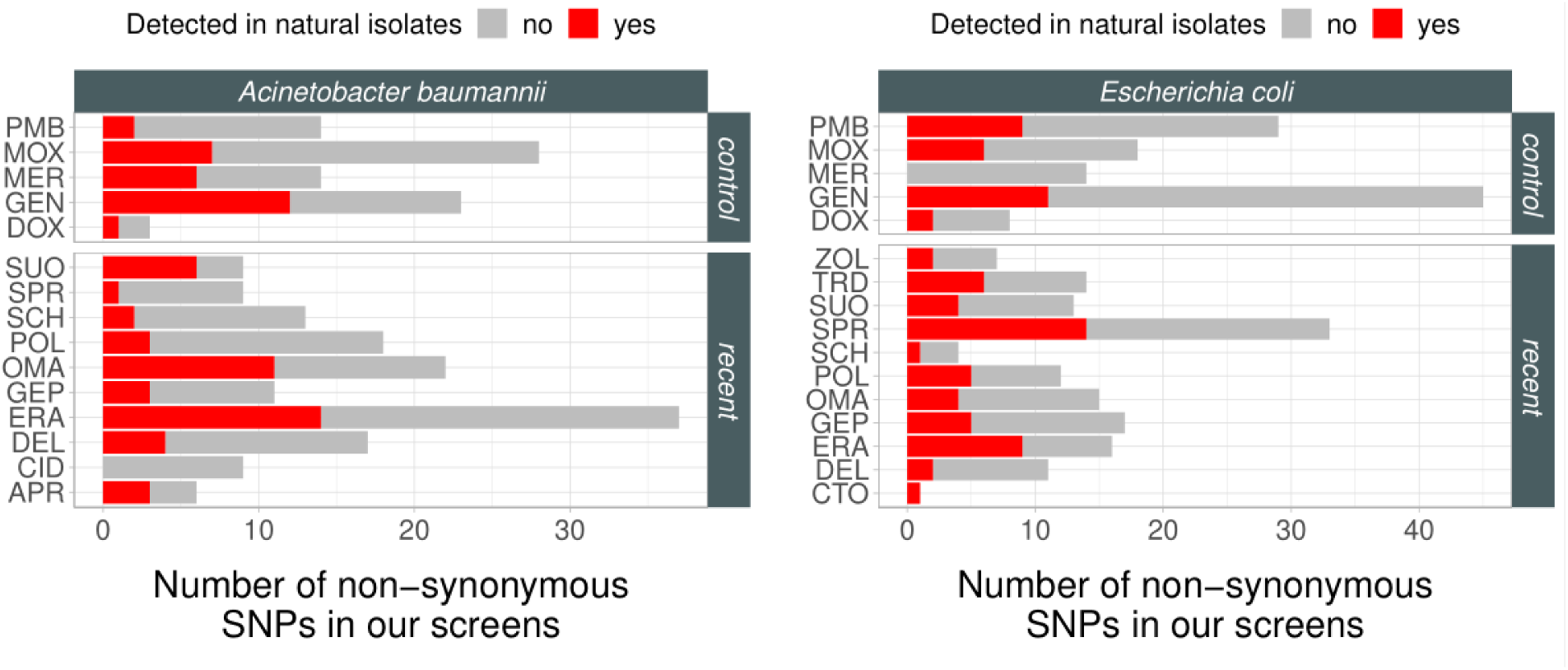
Non-synonymous mutations shared by laboratory evolved lines and natural isolates of *E. coli* and *A. baumannii*. The barplots show the number of non-synonymous mutations found in laboratory evolved *A. baumannii* (left panel) or *E. coli* (right panel) adapted to different antibiotics. Mutations also detected in the genomes of natural isolates of the same species are marked with red, whereas those that remained undetected are marked with gray. No significant difference was found in the fraction of non-synonymous mutations shared by natural strains between control and recent antibiotics (binomial regression model, P = 0.206). For abbreviations, see Table 1.

### Cross-resistance analysis of novel topoisomerase inhibitors

To explore cross-resistance explicitly, we focused on topoisomerase inhibitors, for two reasons. First, this drug class involves a substantial fraction of the antibiotics that are currently in clinical trials. Second, clinically significant levels of resistance against DNA gyrase and topoisomerase IV inhibitors are generally encoded by multiple resistance mutations in the pathogen genome. Resistance genes encoded by plasmids provide a much lower level of resistance against these antibiotics^18^. As the targets of these antibiotics are well-described, deep-scan mutagenesis can find resistance-conferring mutations in these genes in an unprecedentedly comprehensive manner.

We performed deep-scanning mutagenesis in the target genes of moxifloxacin, an established topoisomerase inhibitor. The quinolone resistance-determining regions (QRDR)^25^ of the *gyrA* and *parC* genes were subjected to a single round of mutagenesis using DIvERGE^18^ in *E. coli* and *K. pneumoniae*. The mutant libraries were subsequently exposed to moxifloxacin stress, resulting in an up to 128-fold increment in resistance levels (Extended Data Fig. 17). 20 mutant clones for each species were isolated; the moxifloxacin MIC level reached or was above the clinical breakpoint (0.25 µg/ml) in all cases. Targeted sequencing of QRDR revealed 1 to 3 mutations per clone. In total, 33 different combinations of mutations involving 44 distinct mutations were found (Supplementary Table 10).

Moxifloxacin resistance-conferring mutation combinations reduced susceptibility to topoisomerase inhibitors under clinical development, including delafloxacin and gepotidacin (Extended Data Fig. 17 and Supplementary Table 10). The result with gepotidacin is unexpected, as it is a new topoisomerase inhibitor in development, featuring innovative target sites and modes of action^26^. Prior papers reported that fluoroquinolone-resistant clinical isolates displayed no cross-resistance to this antibiotic, but the data was limited^27^. Remarkably, 52% of the 44 mutations have been previously implicated in resistance to topoisomerase inhibitors or are present in the genomes of natural *E. coli* isolates (Extended Data Fig. 18). Furthermore, natural strains carry some of the specific mutational combinations identified by DIvERGE (Extended Data Fig. 18).

Together these data indicate that prolonged use of topoisomerase inhibitors widely employed in clinical practice selects for mutations provide reduced susceptibility to new antibiotics. Future works should perform detailed mutational-scan analysis with other antibiotic classes to see whether this conclusion holds more generally, including those with larger mutational target size, or to antibiotics with identical target proteins but non-overlapping binding regions.

### Mobile resistance genes are diverse against recent antibiotics

Laboratory evolution can identify resistance mutations only, and thus it ignores the impact of horizontally transferable resistance mechanisms. Therefore, we next explored the abundance of mobile antibiotic resistance genes from environmental and clinical resistomes. We previously created metagenomic libraries from (i) river sediment and soil samples at 7 antibiotic polluted industrial sites in the close vicinity of antibiotic production plants in India (that is, anthropogenic soil microbiome), (ii) the stool samples of 10 European individuals who had not taken any antibiotics for at least one year before sample donation (that is, gut microbiome), and (iii) samples from a pool of 68 multi-drug resistant bacteria isolated in healthcare facilities or obtained from strain collections (that is, clinical microbiome)^28^ (Supplementary Table 11). Each library contained up to 5 million DNA fragments (contigs), corresponding to a total coverage of 25 Gb (i.e., the size of ∼5,000 bacterial genomes). Established functional metagenomic protocols were used to detect small DNA fragments in these libraries (∼1.7 KB long on average) that confer resistance in intrinsically susceptible clinical *E. coli* and *K. pneumoniae* strains^28^.

690 independent DNA fragments emerged against the tested ’recent’ and ’control’ antibiotics, resulting in an up to 256-fold increment in resistance level of the bacterial hosts (Supplementary Table 12, Extended Data Fig. 19). Overall, there is no significant difference in the number of contigs between recent antibiotics and their corresponding within-class controls (paired Wilcox- test, p = 0.791, Fig. 5A). However, no resistance conferring DNA fragment was detected to Tridecaptin M152-P3 in any of the metagenomic libraries and host species (Fig. 5A). Alarmingly, the clinical microbiome was an especially rich source of antibiotic resistance conferring DNA segments (Fig. 5A). In total, 642 non-redundant open reading frames (ORFs) were detected, many of which were present in multiple DNA fragments (Supplementary Table 12). 77% of the 690 DNA fragments displayed close sequence similarity to known resistance genes (i.e., ARG) in relevant databases (Supplementary Table 12). These ARGs are involved in antibiotic inactivation, antibiotic efflux or protect the antibiotic targets (Extended Data Fig. 20), and their phylogenetic origin is diverse (Extended Data Fig. 21).

**Fig. 5A.**
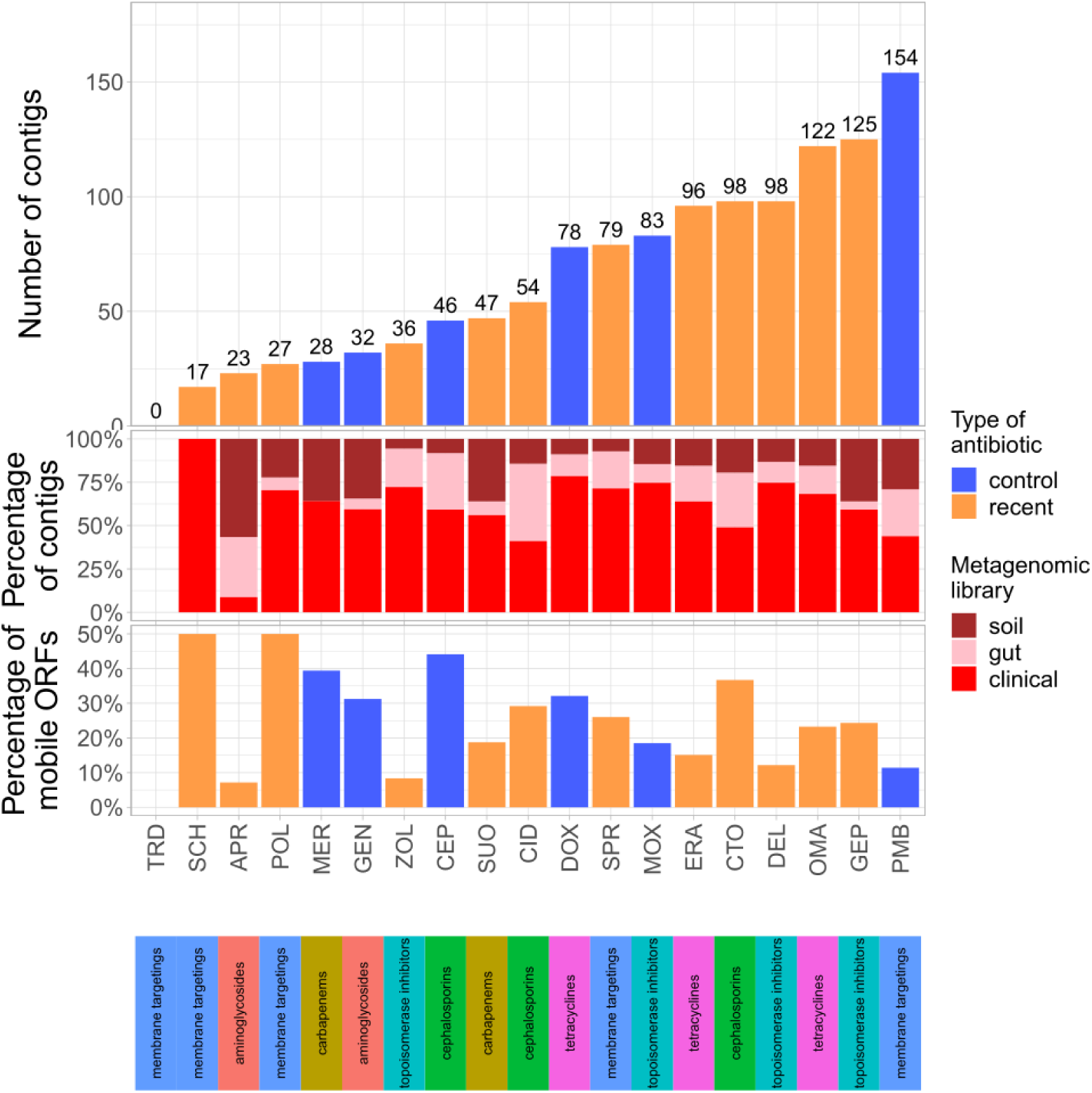
Functional selection of metagenomic libraries with 18 antibiotics resulted in numerous distinct resistance-conferring DNA contigs, with the only exception of Tridecaptin M152-P3 (TRD). The barplot on the top shows the number of unique DNA fragments (contigs) that confer resistance to control (blue) and recent (orange) antibiotics, respectively, whereas the barplots at the bottom show the distribution of the identified resistance conferring contigs across metagenomic libraries and the percentage of mobile ORFs per antibiotic, respectively. ORF mobility was defined by evidence for recent horizontal gene transfer events or presence on mobile plasmids (see Methods). The panel below the barplots denote the class of the antibiotics analyzed.We observed no significant difference in the number of contigs between recent antibiotics and their corresponding within-class controls (paired Wilcox-test, P = 0.4973). The same pattern is true for the percentage of mobile ORFs (paired Wilcox-test, P = 0.576). For abbreviations, see Table 1.

### Prevalence of multi-drug resistance-conferring DNA segments

The functional metagenomics screens also revealed an overlap in the set of resistance-conferring DNA fragments across ‘recent’ and ‘control’ antibiotics (Fig. 5B). This pattern holds for antibiotic pairs with different modes of action, such as topoisomerase inhibitors and tetracyclines. For example, 136 resistance contigs were detected overall against moxifloxacin, a clinically employed topoisomerase inhibitor and omadacycline, a new tetracycline antibiotic, and as high as 50% of those contigs overlap between the two antibiotics. The analysis identified two key genes, *baeR* and *ramA*, repeatedly carried by these contigs. BaeR and RamA enhance the expression of the MdtABC/AcrD^29^ and AcrAB-TolC efflux pump complexes^30^, respectively, with RamA additionally downregulating the porin OmpF expression^31^. This result indicates a broad substrate specificity of these efflux systems, underscoring their ability to extrude a range of antibiotics, including those which are currently in clinical development.

**Fig. 5B.**
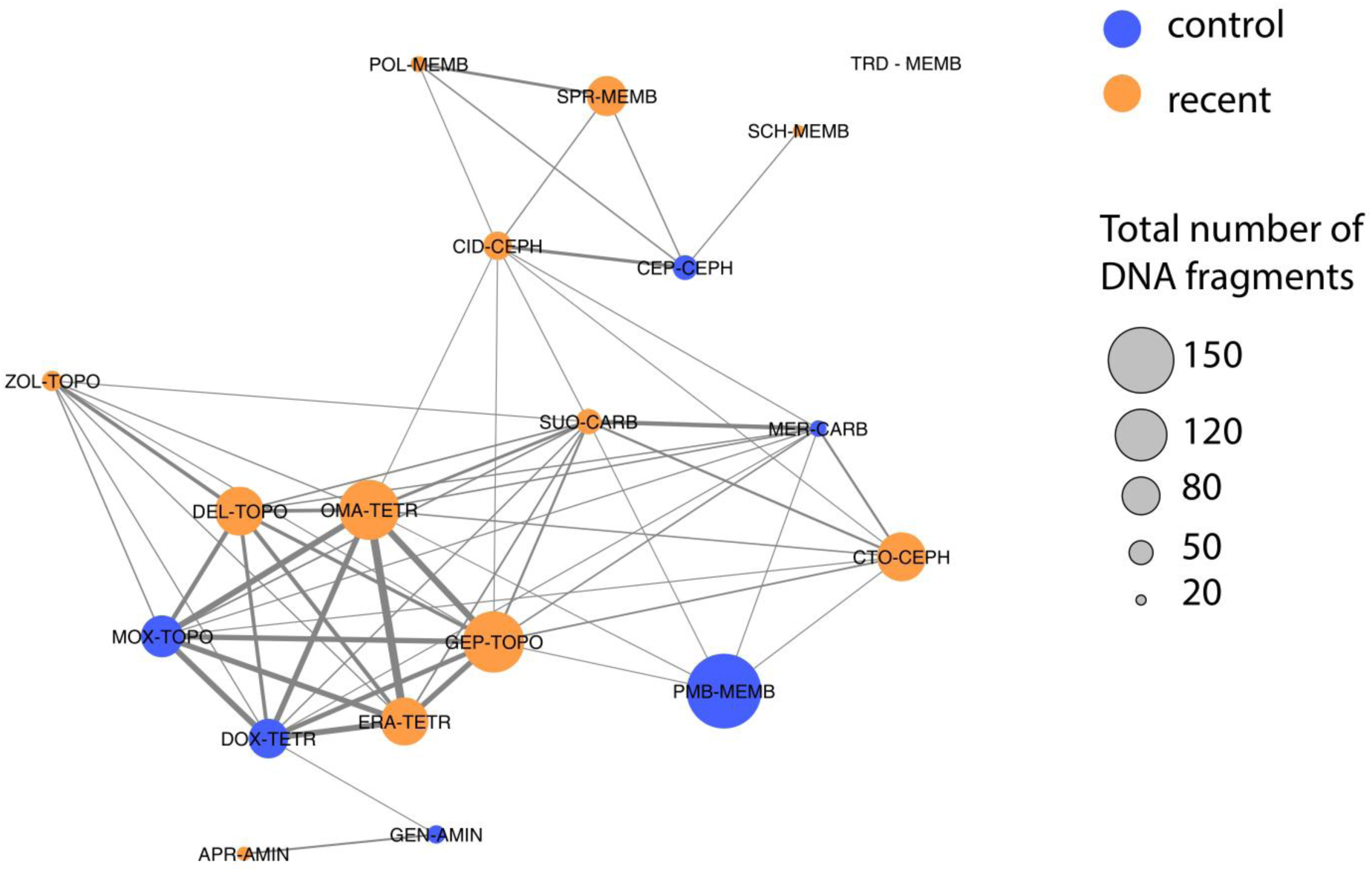
Overlap in the set of resistance conferring DNA-fragments (contigs) across antibiotics. Each node represents a recent (orange) or control (blue) antibiotic, while links indicate overlap in the resistance conferring DNA contigs identified in functional metagenomic screens. The thickness of the link indicates the extent of overlap (calculated by Jaccard similarity) between antibiotic treatments. The size of the nodes corresponds to the total number of detected DNA fragments per antibiotic. The class a given antibiotic belongs to is also indicated (TOPO: topoisomerase inhibitors, TETR: tetracyclines, AMIN: aminoglycosides, CARB: carbapenems, CEPH: cephalosporins, MEMB: membrane targeting antibiotics). For abbreviations, see Table 1.

More generally, the fact that certain DNA fragments confer resistance to more than one antibiotic may reflect that certain antibiotics share a mechanism of action or are chemically similar. To investigate the impact of chemical similarity on co-resistance, we quantified the structural similarity of these pairs using SMILES identifiers and examined their correlation with profile similarity based on detected DNA fragments. We found that antibiotic pairs with similar structures are more likely to exhibit overlapping resistance-conferring DNA fragments in the functional metagenomics screens (Extended Data Fig. 22). This pattern holds when only antibiotic pairs belonging to different antibiotic classes were considered (Extended Data Fig. 22). However, there are significant deviations from this general pattern. For instance, although SPR-206 is a derivative of Polymixin-B, these two antibiotics do not share any common DNA fragments responsible for conferring resistance. Similarly, while the chemical structure of cefiderocol resembles other cephalosporins studied, the addition of a chlorocatechol group transforms it into a siderophore^32^. Consequently, the majority of DNA fragments conferring resistance to cefiderocol are unique to this antibiotic. In summary, these analyses indicate that certain alterations in chemical structure that affect the mode of action or uptake of the antibiotic can lead to major changes in the associated resistance mechanisms.

### Health risk analysis of resistance genes

Next, we evaluated whether the detected putative ARGs could pose threats to public health. Inspired by an earlier study^33^, we considered three major ARG criteria: i) gene mobility, ii) presence in microbiomes associated with the human body, and iii) bacterial host pathogenicity. ARGs were designated as “high risk” if they fulfilled at least two of the three criteria.

Using established methods, gene mobility was defined by evidence for recent horizontal gene transfer events in nature and the presence of the ARGs on natural plasmids derived from diverse environments (Methods). Altogether, 20.7% of the putative ARGs were found to be carried by plasmids or have been subjected to recent horizontal gene transfer events (Fig. 5A; Extended Data Fig. 24, Supplementary Table 12). Next, we asked for the abundance of the identified ARGs in microbiomes associated with the human body. We identified close homologs of the detected ARGS by our functional metagenomic screen in the non-redundant Global Microbial Gene Catalogue (GMGCv1). The catalog summarizes results from over 13 000 publicly available metagenomes across 14 major habitats, including microbiomes from the human body, domestic animal, waste-water, freshwater, and built environments.

Most microbial genes in the catalog are specific to a single habitat^34^. By contrast, 27.6% of the putative ARGs detected in the resistance screens to new antimicrobials were found in multiple habitats, further indicating their potential mobility. In addition, when only habitats associated with the human body (human gut/oral/skin/nose/blood plasma/vagina microbiomes) were considered, this figure rises to 32.7%, indicating that these microbiomes could be a rich source of resistance genes to new antibiotics (Extended Data Fig. 23). Reassuringly, ARGs associated with the human body were also more prevalent in human-related abiotic habitats (wastewater/built environment) than other ARGs (Fisher’s test, Odds ratio: 125, P < 0.0001). Finally, 36.6% of the ARGs are already present in the genomes of bacterial pathogens with critical clinical importance (Extended Data Fig. 23, see Methods and Supplementary Table 12).

In sum, 24.5 % of the 642 ARGs were designated as “high-risk” (Supplementary Table 12): these ARGs are anticipated to possess the greatest potential for catalyzing multidrug resistance in pathogens through a combination of hazardous traits: broad host compatibility enabled by mobility, alongside enrichment in human microbiomes and in bacterial pathogens. However, a significant variation was observed in the frequency of “high-risk” ARGs across antibiotics (Fig. 5C). A notable example is apramycin sulfate (APR), an antibiotic with extensive use in veterinary medicine for decades, and is now in clinical trials for application in humans. Only 2 out of the 63 putative ARGs (3.2%) detected against this antibiotic were designated as high risk, due to a shortage of evidence for their mobility and presence in bacterial pathogens.

**Fig. 5C.**
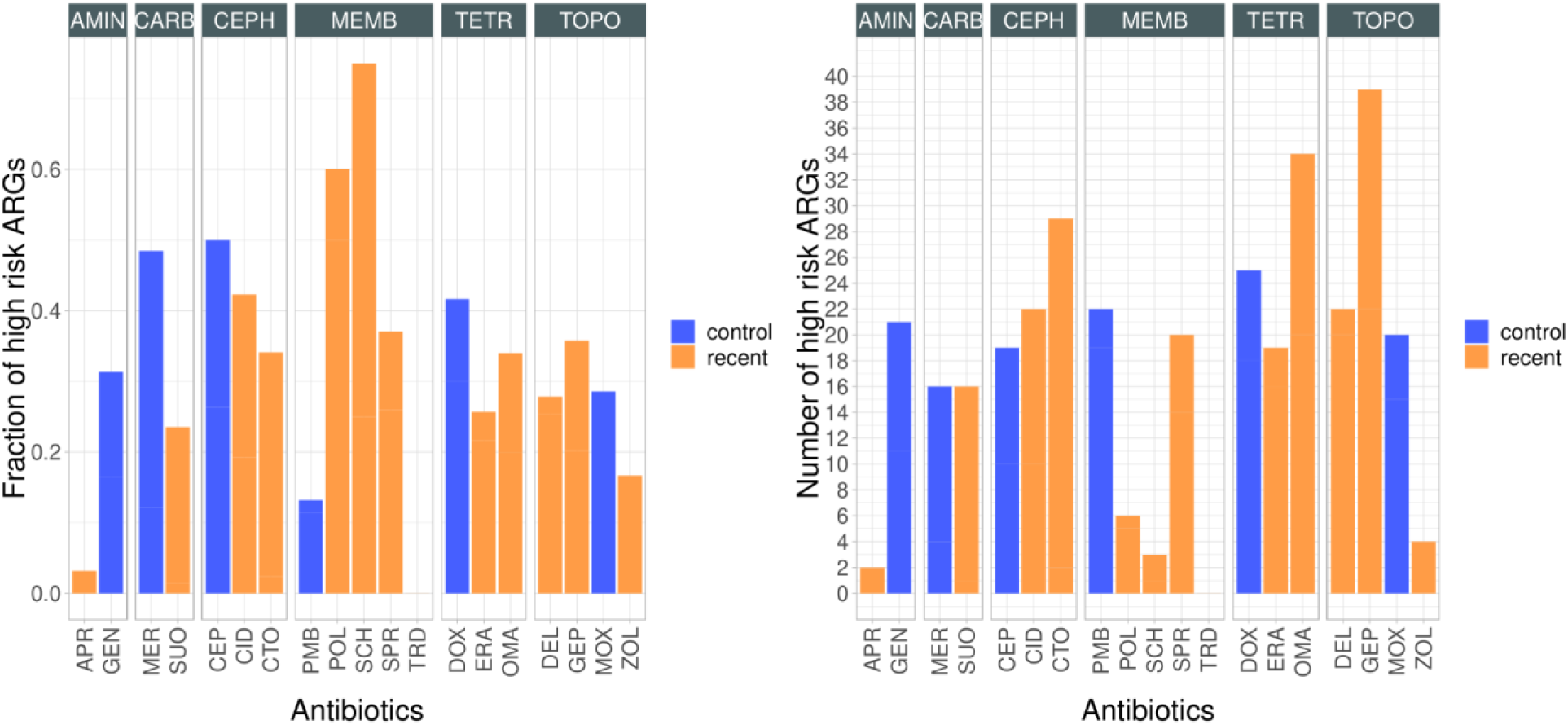
Health risk analysis of putative ARGs. The figure shows the fraction (left panel) and total number (right panel) of health risk ARGs across functional screens against different antibiotics. ARGs were designated as “high risk” if they fulfilled at least two of the following three criteria: i) mobile, presence in ii) human-associated microbiomes, and iii) human pathogens. Blue and orange colors depict control and recent antibiotics, respectively. The analysis revealed a significant variation in the fraction of high-risk ORFs across the antibiotics tested (proportion test, P < 0.05), indicating that certain antibiotics are more likely to be associated with high-risk ORFs compared to others. Antibiotic classes: TOPO (topoisomerase inhibitors), TETR (tetracyclines), AMIN (aminoglycosides), CARB (carbapenems), CEPH (cephalosporins), and MEMB (membrane-targeting antibiotics.

By sharp contrast, several “high-risk” ARGs were detected against new antibiotics, such as sulopenem (N=16), cefiderocol (N=22) and ceftobiprol (N=26). These high-risk ARGs included several β-lactamases, such as New-Delhi-metallo (NDM) and veron-integron metallo-β- lactamases (VIM) (Extended Data Fig. 24). Given the prior expectation of cefiderocol’s lower propensity for resistance development, the high number of high-risk ARGs against cefiderocol is especially alarming.

### Habitat-specificity of resistance genes in Escherichia coli

Finally, we studied the phylogenetic and ecological distribution of the putative ARGs across *E. coli* natural isolates, which occupy a wide range of ecological niches, and display highly variable genomic contents^35^. Specifically, we used genomic data on 16,000 *E. coli* strains that have been categorized into three habitats of isolation (agriculture, human or wild animal hosts) and 11 phylogenetic groups^36^. Overall, 48 putative ARGs have close homologs in the studied *E. coli* genomes (Supplementary Table 12). These genes are generally present in less than 10% of the investigated genomes, suggesting that they are part of a mobile subset of the *E. coli* pangenome (Extended Data Fig. 25). Additionally, they are detectable in the genomes of bacteria isolated from multiple habitats (Extended Data Fig. 26). Notably, natural *E. coli* genomes with at least one putative ARG against certain ‘recent’ antibiotics (such as cefiderocol, ceftobiprole and omadacycline) are frequent in all three main habitats considered (Extended Data Fig. 27, Extended Data Fig. 28). These patterns are not due to exceptionally successful colonization of particular clones, as these putative ARGs are present in *E. coli* genomes belonging to multiple phylogroups (Extended Data Fig. 29) and were isolated from diverse geographical locations (Extended Data Fig. 30).

### Comparison of resistance mechanisms by genomic mutations and mobile resistance genes

A unique aspect of our study is that it allows comparison of the resistance mechanisms identified in the laboratory evolution and functional metagenomic screens. Several main patterns emerged from this study:

The resistance mechanisms developed by genomic mutations and antibiotic resistance genes (ARGs) differ substantially from each other (Fig. 6A). To explore the distribution of different canonical resistance mechanisms across antibiotics, we first pulled information on the mutated genes from the CARD and ResFinder databases^37,38^. 476 unique non-synonymous mutations in our dataset have been associated with genes with established roles in antibiotic resistance, which corresponds to over 30% of all detected cases (Supplementary Table 6). When only genes mutated repeatedly across genetic backgrounds and antibiotic treatments were considered, this figure rises to 48% (Supplementary Table 7). Using the same databases, 36% of the DNA fragments identified in the functional metagenomics screens carry genes that have previously been associated with antibiotic resistance (Supplementary Table 12). The identified resistance mechanisms include antibiotic efflux, reduced membrane permeability, antibiotic target alteration, antibiotic target protection and enzymatic inactivation.

**Fig. 6A.**
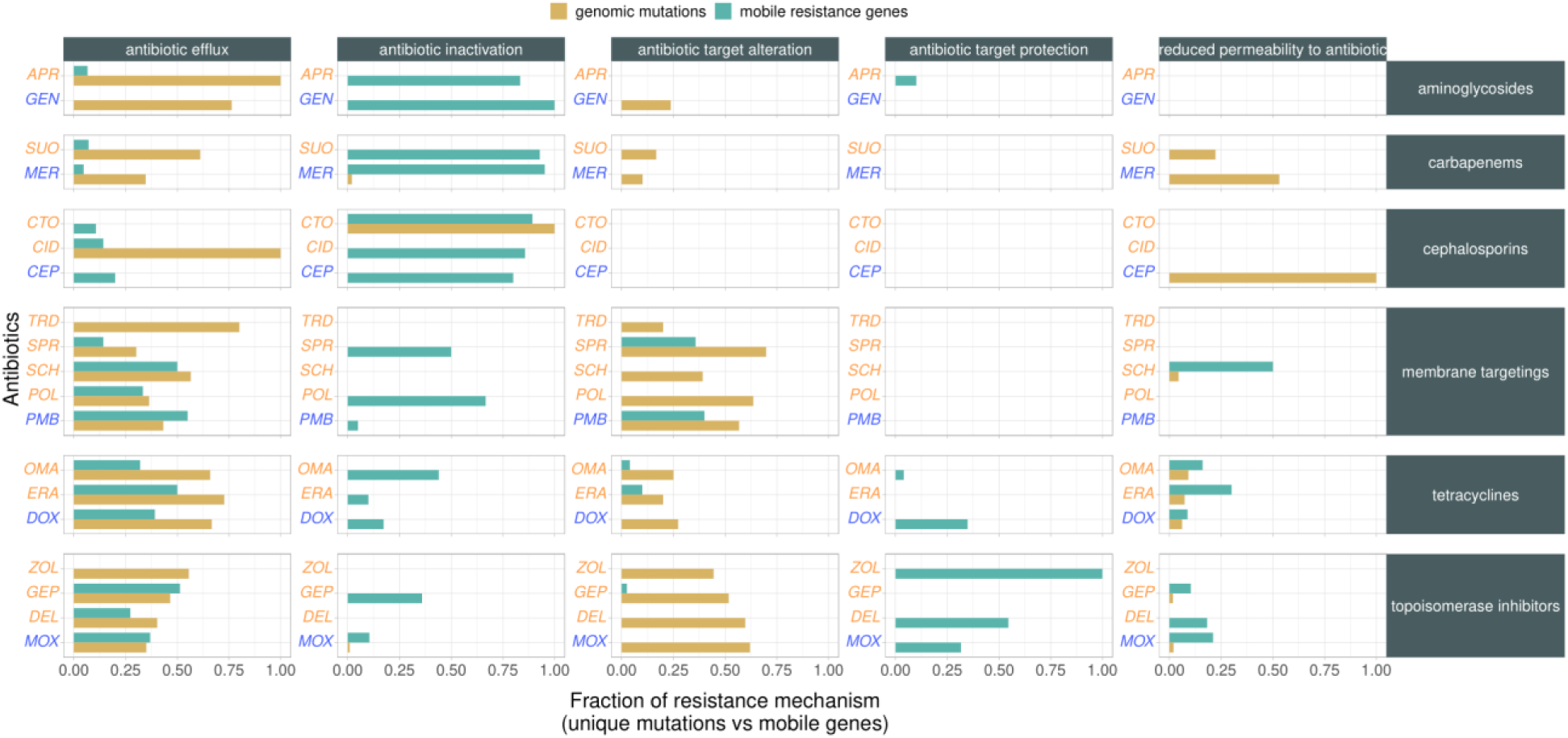
Comparison of the distribution of resistance mechanisms across antibiotics between assays. The figure shows the distribution of resistance mechanisms across different antibiotics (blue and orange indicate control and recent antibiotics, respectively). The fraction of resistance mechanisms are color-coded: those that rely on genomic mutations (ALE/FoR) and ARGs (functional metagenomics) are colored brow and green, respectively. Genetic elements were assigned to five major resistance mechanisms (vertical panels) based on homology to genes featured in the CARD and ResFinder databases.

In general, antibiotic efflux and target alteration were the two most ubiquitous resistance mechanisms derived by genomic mutations (Extended Data Fig. 31, Extended Data Fig. 32, Extended Data Fig. 33), while enzymatic alterations were far more prominent among hits derived from functional genomic screens (Extended Data Fig. 34). In addition, resistance by transfer of efflux pumps was rare for specific new antibiotics (e.g. SPR-206 or zoliflodacin, Extended Data Fig. 34). Antibiotic target protection, a mechanism wherein a resistance protein directly interacts with an antibiotic target to shield it from inhibition, was observed exclusively through functional genomic screens. Among the recognized proteins falling into this category is the quinolone resistance (Qnr) protein family. These pentapeptide repeat proteins facilitate diminished susceptibility to topoisomerase inhibitors, including compounds presently undergoing clinical development. They function by binding to and safeguarding the cellular targets, specifically type II topoisomerases, from the effects of antibiotics^39^.

The prevalence of efflux-mediated resistance displayed substantial heterogeneity across lineages derived from different strains and species (Extended Data Fig. 31). To investigate this issue in more detail, we studied 85 representative laboratory-evolved lines derived from frequency-of-resistance assays, derived from multi-drug resistant and antibiotic sensitive strains of *E. coli*, *K. pneumoniae,* and *A. baumannii*. These lines generally harbored 1-2 mutations only, providing a more direct link between the observed mutations and any potential physiological changes in efflux. In total, 40 out of the 85 studied laboratory-evolved lines displayed reduced intracellular levels of the fluorescent dye Hoechst 33342, indicating enhanced efflux or enhanced membrane permeability (Extended Data Fig. 35). Reassuringly, lines carrying mutations in established efflux pumps displayed reduced intracellular accumulation of the dye (P=0.019, Welch Two Sample T-test). For example, *basS* was repeatedly mutated in *E.coli* in response to polymyxin-B treatment. BasSR is a typical two-component signal transduction system (TCS), that regulates susceptibility to peptide-antibiotics by shaping the transcriptional level of the *emrD* multidrug efflux pump^40^. In other cases (Extended Data Fig. 36), the functional association between the mutated gene and enhanced efflux is not fully established yet. Intriguingly, none of the laboratory-evolved lineages derived from the MDR-resistant *E. coli* strain carried established efflux-related mutations or displayed reduced efflux compared to the corresponding ancestor (Extended Data Fig. 35B). The principles driving the evolution of strain-specific resistance mechanisms will be the subject of a future study.

Additionally, many mutations were found in genes not previously associated with antibiotic resistance. These mutated genes have a broad range of biological functions, including cell wall/membrane biogenesis, lipid metabolism, post-translational protein modifications, recombination and DNA repair, transcription, and translation (Extended Data Fig. 37). Some of these genes may contribute to the evolution of drug resistance by influencing antibiotic tolerance or by promoting the development of new resistance pathways. For a summary of these, see Supplementary Note 4.

### Integrating evidence on resistance to new antibiotic candidates

An ideal antibiotic candidate is expected to meet several essential criteria: i) a broad antibacterial spectrum to ensure effectiveness against a wide array of pathogens; ii) low tendency for resistance development through genomic mutations, iii) scarcity of both intrinsic and horizontally transferred mobile antibiotic resistance genes, and iv) a low prevalence of associated resistance mechanisms in human-associated microbiomes and bacterial pathogens. Unfortunately, none of the compounds investigated in this study simultaneously satisfied all these requirements (see Fig. 6B). By synthesizing several collected data, we calculated an average metric value that served for the ranking of new antibiotic candidates based on their resistance profiles and which showed significant heterogeneity across antibiotic classes (Fig. 6B, Kruskal-Wallis test, P < 0.05). According to this ranking, recent antibiotics targeting bacterial membranes are anticipated to exhibit reduced susceptibility to resistance development in natural settings, in comparison to tetracyclines and topoisomerase inhibitors (Dunn post-hoc test with Benjamini-Hochberg correction for multiple comparisons, P < 0.05). However, there remains significant scope for improvement in their efficacy.

**Fig. 6B.**
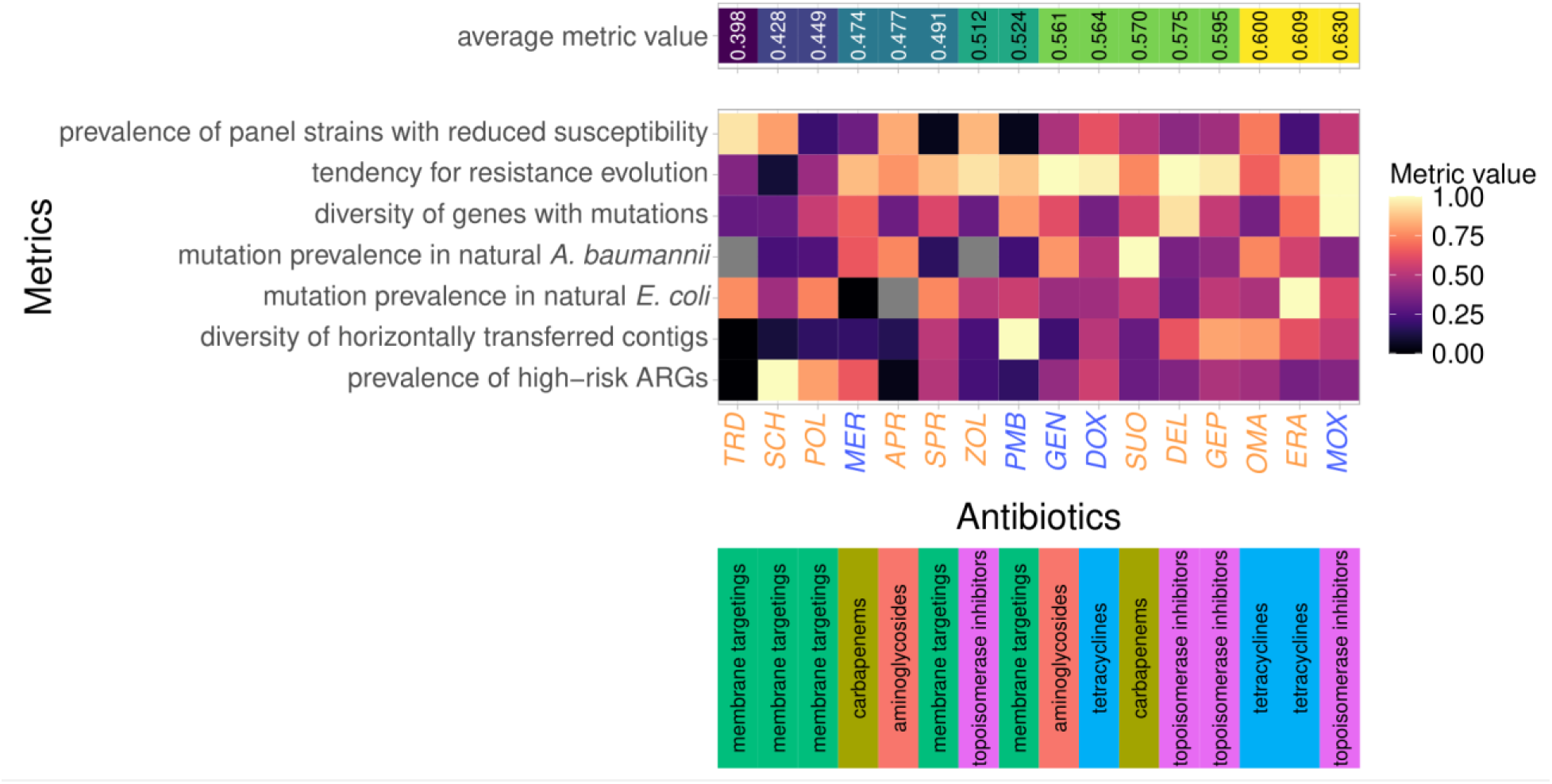
Resisto-plane for each antibiotic studied in this work. The heatmap shows various metrics for each antibiotic studied throughout the work. The metrics include: i) **prevalence of panel strains with reduced susceptibility** - fraction of bacterial strains from a selected pathogen panel with high initial MIC values (see Methods); ii) **tendency for resistance** - the fraction of adapted lines with a relative minimum inhibitory concentration (MIC) exceeding 16, representing the 25% quantile of all relative MIC values; iii) **diversity of genes with mutations** - the fraction of orthogroups exhibiting mutations during adaptive laboratory evolution (ALE) or frequency-of-resistance (FoR) assays with each antibiotic, adjusted by the total number of mutated orthogroups; iv-v) **mutation prevalence in natural *E. coli* or *A. baumannii* strains** - the fraction of laboratory observed adaptive mutations that are already present in natural *E. coli* and *A. baumannii* strains, respectively, vi) **diversity of horizontally transferred contigs** - the count of unique DNA fragments per antibiotic, normalized by the total contig count in functional metagenomics studies;and vii) **prevalence of high-risk ARGs** - the ratio of ARGs considered high-risk, based on meeting at least two of three specified health-risk criteria (see Methods), among all ARGs detected for each antibiotic. Grey color denotes missing value, due to initial resistance in the studied species. Antibiotics are ordered by the average metric value, blue and orange indicate control and recent antibiotics, respectively. The panel below the heatmap denote the class of the antibiotics analyzed.

Below, we offer a comprehensive evaluation of the strengths and weaknesses associated with each group of antibiotic candidates. Specifically, we’ve gathered data from past research asserting that the new antibiotic candidates either resist bacterial adaptation or circumvent established resistance mechanisms. Yet, our findings suggest that these conclusions were based on limited data and are therefore premature. This discrepancy arises from our novel research approach that integrates state-of-the-art methods and multiple bacterial species.

Antibiotics targeting multiple cellular functions are generally expected to be less prone to bacterial resistance. For example, it has been stated that SCH-79797, a dual-targeting antibiotic effectively kills a wide range of bacterial pathogens without detectable resistance^21^, but the underlying data was limited. By contrast to these claims, we found that a relatively low but significant resistance emerged via mutations of regulatory genes of efflux mechanisms (*acrR*, *adeN*, and *adeS*). Similarly, prior papers claimed that the chance of high-level delafloxacin and gepotidacin resistance via target mutations is limited due to the dual-targeting nature of these antibiotics^26,41^. Additionally, these antibiotics were supposed to be poor substrates of established bacterial efflux pumps^42^. In our studies, however, high levels of resistance to these antibiotics evolved via target mutations and mutations often occurred in established efflux pumps (*acrB*) or their corresponding regulatory genes (*acrR*, *adeN*, and *nfxB*).

A similar tendency was evident regarding DNA segments that provide resistance when transferred to new bacterial hosts. Omadacycline, for example, was previously stated to evade resistance via tetracycline-specific efflux pumps, based on the lack of cross-resistance^43^. In our study, however, contigs containing tetracycline-specific efflux pumps *tetA* and *tetX* came up as hits against omadacycline.

Due to its unique mechanism of uptake, resistance to cefiderocol was supposed to be relatively immune to the development of resistance^32^. However, *baeS and crP,* two regulatory genes involved in antibiotic efflux were mutated in response to cefiderocol. Indeed, prior works showed that mutations in these genes constitutively activate the BaeSR two-component regulatory system to increase the expression of the MdtABC efflux pump^44^. In addition, in our functional metagenomic screen, carbapenemases NDM-15, 22, and 27 provided resistance against cefiderocol. Similarly, sulopenem (SUO) is a broad-spectrum thiopenem β-lactam antibiotic being developed to treat infections caused by multidrug-resistant and cephalosporin-resistant bacteria belonging to the *Enterobacteriales* group^45^. However, it was prone to resistance in multiple screens. In particular, functional metagenomics identified several DNA fragments carrying New-Delhi-metallo (NDM)- β-lactamase genes (Supplementary Table 8, Supplementary Table 12).

Eravacycline (ERA) was specifically designed to overcome resistance to common tetracycline-specific efflux and ribosomal protection mechanisms^46^. Although eravacycline has a relatively broad antimicrobial spectrum, resistance to this compound evolves rapidly in the laboratory through modification of efflux pump activities (Fig. 3A, 3B, Extended Data Fig. 35).

Finally, it was claimed that resistance is limited to new antibiotics with unique targets, such as Tridecaptin M152-P3 and POL-7306. Tridecaptin M152-P3 (TRD) is a non-ribosomally synthesized lipopeptide antibiotic candidate that binds Lipid II and thereby disrupts bacterial membrane potential^47^. The essential bacterial membrane-anchored cell-wall precursor Lipid II has been considered a promising target for antibiotics with limited resistance. Indeed, no resistance-conferring DNA fragment was detected against Tridecaptin M152-P3 in any of the metagenomic libraries and host species (Fig. 6B). However, Tridecaptin M152-P3 was particularly prone to resistance by genomic mutations: up to 32-fold increments in resistance levels were detected in laboratory evolved lines via mutations in genes that shape membrane integrity (e.g. *lptD* or *lptX*). In addition, mutations in *phoQ* previously stated to provide no cross-resistance between POL-7306 and colistin^48^, appeared in multiple POL-7306 resistant lines. Resistance to another peptide-based antibiotic, SPR-206 (SPR) readily emerged by genomic mutations. SPR-206 is in clinical trials with potent activity against a wide range of multidrug-resistant bacteria^22^. However, an increase in resistance level as high as 128-fold emerged in *K. pneumoniae* because of single mutations in the two-component BasS/BasR regulatory system.

Overall, the need for more thorough resistance studies during early development phases is glaringly obvious. To gain an accurate picture of bacterial resistance potential, a broader approach incorporating diverse strains and multiple complementary assays is necessary.

## Discussion

Prioritizing potential antibiotics that are less prone to resistance is crucial for advancing antibiotic development. For this purpose, we conducted a thorough investigation by combining various approaches in multiple clinically relevant pathogens: (i) susceptibility testing across a large panel of pathogens, (ii) high-throughput adaptive laboratory evolution experiments, (iii) functional metagenomics with two species, (iv) comprehensive genome sequencing of numerous antibiotic-adapted strains, and (v) extensive bioinformatics analyses and database searches to identify mutations and horizontally transferred genetic elements conferring resistance. We assessed 13 recently developed antibiotic candidates and 6 control antibiotics against various bacterial pathogens to explore and contrast their resistance evolution patterns (Fig. 1).

Our work showed that bacterial resistance generally evolves rapidly against both control and recent antibacterial compounds in the laboratory, regardless of the length of antibiotic exposure and the methodological approaches (Fig 3B, Fig 5A, Extended Data Fig. 19). These patterns also hold for compounds that have novel or dual modes of action and have been previously suggested to be relatively immune to bacterial resistance in the laboratory (Fig. 3A and 3B). Notably, genomic mutations that accumulated during the course of laboratory evolution are likely clinically relevant, as they are prevalent in the genomes of clinical bacterial isolates (Fig. 4C). These results predict that resistance to new antibiotics will arise through selection for pre-existing resistant strains via de novo mutations as well as with horizontally transferable genetic elements (Extended Data Fig. 26 and 27). These findings raise the question of why the resistance genes and mutations to antibiotic candidates that have not yet reached the market are already present in the environment.

One possibility is that due to overlap in resistance mechanisms, prolonged antibiotic exposure both in clinics and agriculture has selected for resistance mechanisms that simultaneously reduce susceptibility to antibacterial compounds still in development. This possibility is further supported by the observation that antibiotics with similar structures tend to share resistance-conferring genetic mechanisms, as evidenced in functional metagenomics screens (Fig. 5B). This trend holds even when considering antibiotic pairs belonging to different classes (Extended Data Fig. 22). Overall, these observations suggest that minor chemical modifications are insufficient to circumvent established resistance mechanisms. The general lack of efforts towards novelty in the clinical antibiotic development pipeline is supported by our data as the majority of the tested antibiotics were highly prone to resistance in our integrated resisto-plane analysis. Contrastingly, all three antibiotics that reached the lowest overall score (e.g. SCH79797, POL-7306, Tridecaptin M152-P3) in the analysis, have novel modes of action (Fig. 6B) Properties that make these antibiotic candidates less resistance-prone will be analyzed in detail in a future work.

Several lines of evidence suggest that the genetic mechanisms conferring resistance in clinical isolates to conventional antibiotics could render new antibiotic candidates ineffective. First, there is a strong positive correlation between the fraction of resistance to recent antibacterial compounds and that of control antibiotics across pathogenic isolates (Fig. 2C). Second, several genes mutated repeatedly across different antibiotic treatments in the evolved lines (Fig. 4A). Third, metagenomics screens revealed an overlap in the set of resistance-conferring DNA fragments between new antibacterial compounds and clinically employed antibiotics (Fig. 5B). This pattern also holds for antibiotic pairs with different modes of action, such as topoisomerase inhibitors and tetracyclines. Finally, deep-scanning mutagenesis established specific mutational combinations that provide cross-resistance between control and recent topoisomerase inhibitors (Extended Data Fig. 17).

Our work highlights the concern that antibiotic development is currently dependent solely on susceptibility indicators in various bacterial pathogens. While indeed many of the novel antibiotics have improved antibacterial spectrum compared to their predecessors, our studies show that this is rarely paired with favorable resistance properties. Specifically, eravacycline shows great antibacterial activity against a panel of bacterial pathogens (Extended Data Fig. 3), however, it is especially prone to resistance by genomic mutations and horizontal gene transfer (Fig. 3B, Fig. 5A).

Future application of the same antibiotic to initially susceptible pathogens can have different outcomes depending on their capacity to evolve resistance^19,49^. Indeed, we have found that the level of resistance achieved during adaptive laboratory evolution was highly contingent upon the bacterial species and strains studied, particularly in an antibiotic-specific manner (Extended Data Fig. 9, 10 and 11). This variability may stem from strain specific differences in initial susceptibility to a given antibiotic, presence of efflux pumps and/or the influence of specific ’potentiator’ genes, which facilitate new mutational pathways towards resistance through epistatic interactions with resistance mutations. These hypotheses will be studied thoroughly in future work, which might aid in developing species-specific therapeutic options to counter the rapid development of resistance^50^. Our work also highlights the risk that antibiotic development programs waste considerable resources on antibiotic candidates prone to resistance if they concentrate solely on a single bacterial species or only on resistance emergence arising from genomic mutations. An important limitation of this study is the lack of systematic investigation of trade-offs of antibiotic resistance, especially on bacterial fitness and virulence, an issue that will be covered in future work. Future works should also decipher the exact role of the identified mutations by re-introducing them individually and in combinations into a wild-type genetic background, and studying the impact of the susceptibility of the resulting mutant strains to new antibiotics. In addition, resistance against Gram-positive specific antibiotic candidates and combination therapies involving new antibiotics will be studied elsewhere.

In sum, the framework provided in this work highlights the importance of testing resistance evolution and the underlying mechanisms rigorously with complementary methods and in multiple relevant bacterial species. Application of such a framework is feasible and advisable to new antibiotics before acceptance for clinical usage as it would allow us to more accurately assess both the immediate efficacy and the long-term utility of antibiotic candidates, as well as to understand the potential health risks associated with resistance development. Although our findings indicate that none of the tested compounds meets all the criteria for an ideal future antibiotic, they also highlight opportunities for improving certain critical properties (see Fig. 6B). This underscores the pressing need for innovative approaches in the discovery and optimization of new antibiotics, particularly those that address the challenges of efficacy and resistance.

## Methods

### Strains, antibiotics and media

This study focused on multiple bacterial strains, as follows. We tested the activity spectrum of the antibiotics in this study on a set of 40 clinically relevant pathogenic strains of four species (*E. coli, K. pneumoniae, A. baumannii*, and *P. aeruginosa*; see the whole list of pathogens in Supplementary Table 2). For the FoR and ALE experiments, 2 strains per species were chosen: *E. coli* ATCC 25922, *K. pneumoniae* ATCC 10031, *A. baumannii* ATCC 17978, and *P. aeruginosa* ATCC BAA-3107 as sensitive (SEN) strains, and *E. coli* NCTC13846, *K. pneumoniae* ATCC 700603, *A. baumannii* ATCC BAA-1605 and *P. aeruginosa* LESB58 as multi-drug resistant (MDR) strains. In the case of *E. coli,* we chose the ATCC25922 strain as sensitive strain, due to its widespread usage in the literature, and an MCR-1 carrying *NCTC 13846* as the MDR strain, due to the high interests in the impact of this mobile resistance gene on colistin resistance^51^. For the other 3 species, sensitive and MDR strains were selected based on the highest number of control antibiotics they showed sensitivity or resistance to, respectively, with an additional criterion for MDR strains, that they should be part of an official strain collection (Extended Data Fig. 1). Functional metagenomic screens were performed with *E. coli* ATCC 25922 and *K. pneumoniae* ATCC 10031 strains. Deep-scanning mutagenesis (DIvERGE) was performed with *E. coli* K-12 MG 1655 and *K. pneumoniae* ATCC 10031 strains.

Nineteen antibiotics were applied in this study from six different antibiotic families, thirteen newly developed (‘recent’) antibiotics, which are in different phases of clinical trials and six conventional (‘contol’) antibiotics with long clinical history, from each antibiotic family. For name, abbreviation and further details see Table 1 and Supplementary Table 1. Antibiotics were custom synthesized or were purchased from several distributors (Supplementary Table 1). Upon preparation, each antibiotic stock solution was filter-sterilized and kept at −20 °C until usage. For more details on ‘recent’ and ‘control’ antibiotics, see Supplementary Note 1, Table 1 and Supplementary Table 1. For data on clinical breakpoints and peak plasma concentrations, see Supplementary Table 1.

Unless otherwise indicated, cation-adjusted Mueller-Hinton Broth 2 (MHB, Millipore) medium was used throughout the study, except for cefiderocol (CID) and a folate biosynthesis inhibitor, SCH79797 (SCH). Following EUCAST recommendation on cefiderocol, iron-depleted MHB media was used^52^. To maximize antibacterial activity of SCH, based on prior experience with folate biosynthesis inhibitor antibiotics^53^, Minimal Salt (MS) medium was used (1 g/L (NH_4_)_2_SO_4_, 3 g/L KH_2_PO_4_ and 7 g/L K_2_HPO_4_) supplemented with 1.2 mM Na_3_C_6_H_5_O_7_ × 2H_2_O, 0.4 mM MgSO_4_, 0.54 μg/mL FeCl_3_, 1 μg/mL thiamine hydrochloride, 0.2% Casamino-acids and 0.2% glucose).

### High-throughput MIC measurements

A standard serial broth microdilution technique^54^ was used to determine MICs, as suggested by the Clinical and Laboratory Standards Institute (CLSI) guidelines. A robotic liquid handling system was used to automatically prepare 11 to 16-step serial dilution in 384-well microtiter plates. 5 × 10^5^ bacterial cells per mL were inoculated into each well containing 60 µl medium. Bacterial cultures were incubated at 37°C with continuous shaking (300 RPM) for 18 hours (2 replicates from each). Cell growth was monitored by measuring the optical density (OD600 values, using Biotek Synergy microplate reader). MIC was defined as the antibiotic concentration of complete growth inhibition (i.e., OD600 < 0.05). The same protocol was used to estimate antibiotic susceptibility of laboratory evolved lineages. Relative MIC was calculated as follows: log_2_(MICevolved / MICancestor). Increase in MIC was calculated as follows: log_10_(MICevolved – MICancestor).

We aimed to perform both frequency-of-resistance (FoR) and adaptive laboratory evolution (ALE) assays with all selected 8 bacterial strains, however, 34% of all antibiotic-strain combinations (N = 52) were excluded from further experiments due to modest initial drug efficacy (i.e., MIC > 4 µg/ml, rendering them less relevant for clinical use) (Supplementary Table 3). Prevalence of panel strains with reduced susceptibility against a certain antibiotic was estimated by calculating the fraction of panel strains with high initial MIC values.

### Frequency-of-resistance assays

To estimate the frequency of spontaneous mutations that confer resistance in a microbial population, frequency of resistance (FoR) assay was used. Using standard protocols^14–17^, approximately 10^10^ cells from stationary-phase cultures were plated to antibiotic-containing MHB broth plates. Before plating, bacteria were grown overnight in MHB medium at 37 °C, 250 RPM, collected by centrifugation, and washed once in equal volumes of MHB broth. From this concentrated bacteria suspension ∼10^10^ cells were plated to agar plates containing the selective drug at the desired concentration (i.e., 2×, 4×, 8× and 20× MIC of each given antibiotic). Unless otherwise indicated (see the section above on high-throughput MIC measurements), MHB agar medium was used throughout the study. All experiments were performed in 3 replicates. Plates were grown at 37 °C for 48 hours. Total colony forming units (CFUs) were determined simultaneously in each experiment by plating appropriate dilutions to antibiotic-free MHB agar plates. Resistance frequencies for each bacterial strain were calculated by dividing the number of formed colonies by the initial viable cell count. Ten bacterial colonies from the highest antibiotic concentration were selected for further MIC measurements and whole genome sequence analysis.

### High-throughput adaptive laboratory evolution

A previously established protocol^55,56^ was employed for adaptive laboratory evolution, with the aim to ensure that populations with the highest level of resistance were propagated further. Starting with antibiotic concentration resulting in ∼50% growth inhibition, 10 parallel populations per antibiotic-ancestor strain combination were grown for 72 h, at 37 °C and continuous shaking (300 RPM). As rapid degradation has been observed for *β*-lactams and cephalosporins in liquid laboratory media^57^, ceftobiprole, cefiderocol and cefepime were not subjected to adaptive laboratory evolution. Unless otherwise indicated, MHB broth medium was used. After each incubation period, 20 μl of 370 of each bacterial culture was transferred to four new independent wells containing freshly prepared medium containing different antibiotic concentrations (0.5×, 1×, 1.5×, and 2.5× the concentration of the previous step). A chess-board layout was used on the plate to monitor potential cross-contamination events. Cell growth was monitored prior to each transfer by measuring the optical density at 600 nm (OD600 value, Biotek Synergy 2 microplate reader). Only populations with the highest drug concentration (and reaching OD600 > 0.2) were selected for further transfer. The evolution experiment was generally continued for 20 transfers, resulting in a total of 728 evolved lines (78 lines were omitted because of limited growth).

### Whole genome sequencing

To identify potential antibiotic resistance conferring mutations, we generally selected 2 to 5 lines from frequency-of-resistance (FoR) and laboratory evolution (ALE) experiments, respectively, for whole genome sequencing. Resistant populations were grown overnight in antibiotic-free medium. DNA isolation from overnight cultures was performed with GenElute Bacterial Genomic DNA Kit (Sigma), according to the manufacturer’s instructions. DNA was eluted 120 µl RNAse free sterile water, in 2 elution steps. 60 µl of the eluted DNA was then concentrated by DNA Clean and Concentrator Kit (Zymo), according to the manufacturer’s instructions. Final DNA concentration was measured with a Qubit Fluorometer and concentration was set to 1 ng/ml in each sample. Sequencing libraries from isolated genomic DNA were prepared with Nextera XT DNA library preparation kit (Illumina) following the manufacturer’s instructions. The sequencing libraries were sequenced on an Illumina NextSeq 500 sequencer using 2*150 PE mid or high output flow cells to generate 2 × 150 bp paired-end reads.

To determine and annotate the variants, we mapped the sequencing reads to their corresponding reference genomes using an established method (Burrows-Wheeler Aligner)^58^. From the aligned reads, PCR duplicates were removed with the Picard Mark Duplicates tool (see http://broadinstitute.github.io/picard/). We removed every read that has been aligned with more than 6 mismatches (disregarding insertions and deletions). The SNPs and INDELs were called with Freebayes^59^ with the following parameters: -p 5--min-base-quality 28. The identified variants were filtered by the vcffilter tool from vcflib^60^ using the following parameters –f ’QUAL > 100’. To avoid missing rare but valid hits, we did not set a lower limit for the prevalence of rare variants. If necessary, mutations were also manually inspected within the aligned reads using IGV^61^ to reduce BWA alignment or freebayes artefacts. Finally, the variants were annotated with SnpEf, and kept only those that were not present in the ancestor. We filtered out mutations that appeared in more than nine lines, as these variants are likely already present in the ancestor as well. Furthermore, mutations that appeared in less than nine but more than six lines were manually inspected to exclude sequencing artefacts. We also excluded mutations that affect 40 bp or longer repetitive regions, resulting in a filtered set of mutations. Lines containing mutations in *the mutL, mutS* or *mutY* genes, or lines with more than 19 mutations, defined by outlier filtering (Q3 + 3*IQR), were considered hypermutators and were subsequently discarded.

To analyze the presence / absence of mutations across genes and strain backgrounds, we first complemented existing gene functional annotation for the 8 bacterial strain backgrounds as follows. Nucleotide sequence and annotation files of 6 strains (*E. coli* ATCC 25922, *K. pneumoniae* ATCC 10031, *A. baumannii* ATCC 17978, *P. aeruginosa* ATCC BAA-3107, *K. pneumoniae* ATCC 700603, and *A. baumannii* ATCC BAA-1605) were downloaded from the ATCC database (https://www.atcc.org/). For *P. aeruginosa* LESB 58 and *E. coli* NCTC13846 strains, genomic data was downloaded from NCBI Nucleotide (accession numbers FM209186.1 and NZ_UFZG00000000.1). Next, all genes in the GeneBank files, including hypothetical ones, were functionally annotated using PANNZER2^62,63^. To compare the sets of mutated genes across strain backgrounds, we determined genes that are shared across different strains by identifying groups of orthologous genes using OrthoFinder (version:2.5.4)^64,65^.

### Bioinformatics analysis of mutations promoting growth in the laboratory based on prior works

We compiled a comprehensive list of 104 genes associated with medium adaptation in *E. coli*, as identified in two previous studies^66,67^. First, we examined our evolved *E. coli* strains for mutations within these genes. The DNA sequences of these genes from the *E. coli* strain MG1655 were retrieved from EcoCyc (version 26.0)^68^. We then aligned the sequences to the amino acid sequences of proteins in our reference genomes (SEN and MDR, separately) using the xblast tool (implemented in the rBLAST R package^69^). For each gene, we selected the alignment with the highest bit-score, requiring a sequence identity of at least 80% and coverage of at least 80%. This approach resulted in 92 of the 104 genes being matched in each reference genome. Among these, 8 genes in the MDR strains exhibited 13 mutations, while 7 genes in the SEN strains contained 15 mutations, totaling 11 genes with 28 mutations in at least one strain. As a next step, we investigated non-coding mutations in our evolved strains to ascertain if any were located in or adjacent to operons overlapping with the genes implicated in medium adaptation. This analysis did not reveal any non-coding mutations associated with the genes of interest.

### Comparison of variants to public genomes of bacterial isolates

We assessed whether amino acid substitutions occurring in the FoR and ALE samples are present in natural populations of *E. coli* and *A. baumannii* as follows. We compiled a comprehensive genomic dataset for *E. coli* strains by downloading assembled genome sequences or un-assembled reads, and metadata, from four sources: i) the JGI Integrated Microbial Genomes & Microbiomes (IMG) database^70^ on 29 Jan 2020, ii) the NCBI RefSeq prokaryotic collection [Available from: https://www.ncbi.nlm.nih.gov/refseq/] on 29 Jan 2020; iii) genomes that were analyzed in the study of Moradigaravand et al.^71^; and iv) genomes that were analyzed in the study of Johnson et al.^72^.

After trimming the adapters with the Cutadapt v3.2 program^73^, we de novo assembled the next-generation sequencing short reads (downloaded from the Sequence Read Archive (SRA) database^74^) of genomes of source iii) and iv) using the SPAdes v3.14.1 software^75^. Then, we applied the BUSCO v5.0.0. workflow^76^ to exclude genome sequences with less than 95% of the BUSCO genes, indicating inadequate completeness / quality. When multiple genome sequences belonged to the same BioSample ID, only the one with the highest BUSCO score, longest sequence and fewest contigs were kept, while all metadata of the original sequences were merged. This resulted in 20,786 *E. coli* genomes (Data S1) for which gene prediction was performed using Prodigal (Version: 2.6.3,^77^) to obtain protein coding gene annotations that are consistent across the genomes. ORFs with less than 100 AAs were filtered out. Strains were classified as pathogens and non-pathogens based on their genomic metadata. For *A. baumannii* strains, we downloaded all available assembled genome sequences from the NCBI RefSeq prokaryotic genome collection (Available from: https://www.ncbi.nlm.nih.gov/refseq/) on 12 Sep 2022. Then we applied genome filtering with 95 percentage completeness using BUSCO v5.4.6. and protein prediction using Prodigal as described above for *E. coli*. This resulted in 15,185 *A. baumannii* genomes (Data S1).

Next, we searched for the presence of each amino acid changing SNP across the *E. coli* and Acinetobacter genome collections as follows. First, we performed sequence similarity search of each gene carrying a given variant using DIAMOND blast (version 2.0.2,^78^) using an e-value of 0.00001 with 90% coverage and 90% identity to identify homologs among the genomes. In the next step, we performed multiple sequence alignment using MAFFT with the --retree 2 option (version: 7.475,^79^). Then, we analyzed the amino acid frequency across the alignments in all mutated positions. All *E. coli* and *A. baumannii* variants that were present in the corresponding species’ genome collection and appeared more than once in our FoR and ALE samples were selected for further analysis.

### Functional metagenomic screens

Resistance conferring DNA fragments in the environment were identified by functional selection of metagenomic libraries. In a previous work^28^, we created metagenomic libraries to obtain environmental and clinical resistomes, including i) river sediment and soil samples from seven antibiotic polluted industrial sites in the close vicinity of antibiotic production plants in India^80^ (anthropogenic soil microbiome), ii) fecal samples from 10 European individuals who had not taken any antibiotics for at least 1 year before sample donation (that is, gut microbiome) and iii) samples from a pool of 68 multi-drug resistant bacteria isolated in healthcare facilities or obtained from strain collections (see Supplementary Table 11). For full details on library construction, see Apjok and colleagues^28^.

Briefly, environmental and genomic DNA was isolated using the DNeasy PowerSoil Kit (Qiagen) and the GenElute Bacterial Genomic DNA Kit (Sigma), respectively. Environmental and genomic DNA was enzymatically fragmented followed by the size selection of 1.5-5 kb long fragments. Metagenomic inserts were cloned into a medium-copy-number plasmid, containing 10-10 nt long barcodes flanking the insert (uptag and downtag). Library sizes ranged from 2-6 million clones with an average insert size of 2 kb.

Libraries were introduced into *K. pneumoniae* ATCC 10031 and *E. coli* ATCC 25922 by bacteriophage transduction (DEEPMINE)^28^ and electroporation, respectively. DEEPMINE employs hybrid T7 bacteriophage transducing particles to alter phage host-specificity and efficiency for functional metagenomics in target clinical bacterial strains.

In the current study, we followed previously described protocols with two minor modifications. First, transducing hybrid phages were generated with a T7 phage lacking the gp11-gp12-gp17 genes, constructed as previously^81^. Second, we used a new phage tail donor plasmid for complementing the deleted phage tail genes. This plasmid was cloned using the ΦSG-JL2 phage tail coding genes, the packaging signal region of T7 phage and the pK19 plasmid backbone based on a previous work^82^.

Functional selections for antibiotic resistance were performed on MHB agar plates containing a concentration gradient of the antimicrobial compounds^83,84^. Cells containing the metagenomic libraries were plated in a cell number covering at least 10 times the size of the corresponding metagenomic library. Plates were incubated at 37°C for 24 h. For each functional selection, a control plate was prepared with the same number of cells containing the metagenomic plasmid without a cloned DNA fragment in its multi-cloning site. These control plates showed the inhibitory zone of the antimicrobial compound. To isolate the resistant clones from the libraries, sporadic colonies were identified above the inhibitory zone based on the control plate by visual inspection. Colonies were then collected for plasmid isolation (Thermo Scientific GeneJET Plasmid Miniprep Kit). Metagenomic inserts in the resistant hits were sequenced by two complementary sequencing methods. First, random 10 nt barcodes flanking the metagenomic inserts on the resistant plasmids from each selection experiment were PCR amplified. For this, we used primers that contain 2x8 nt long barcodes specific for each selection experiment. Amplicons were pooled, size selected on agarose gel and sequenced by Illumina. Second, metagenomic inserts and their flanking 10 nt uptag and downtag barcodes were sequenced by Nanopore.

### Annotation of antibiotic resistance genes (ARGs)

Consensus insert sequences from Nanopore sequencing were matched with the respective selection experiment using the data from Illumina sequencing. First, sequencing reads from Illumina sequencing were demultiplexed using the 2x8 nt long barcodes specific to the selection experiment, and then the demultiplexed reads were matched with the consensus insert sequences using the random 10 nt long barcodes specific to the metagenomic inserts. To reduce redundancy and spurious matches, the list of metagenomic contigs were filtered 1) to unique barcodes, keeping barcodes with the highest nanopore read count and 2) to contigs that were supported by at least 8 nanopore reads and 5 illumina reads. Prediction of ARGs within these contigs was based on ORF prediction using Prodigal v2.6.3,^77^, followed by searching the annotated ORFs within the CARD and ResFinder databases^37,38^. Searches were performed using blastx from NCBI BLAST v2.12.0^85^ with 10^-5^ E-value threshold and otherwise default settings. ORFs were clustered at 95% identity and coverage using CD-HIT v4.8.1,^86^ and only one representative ORF was kept for each cluster. The inserts were classified based on whether or not any ARGs were found in them, and whether or not at least one of these ARGs was associated with the antibiotic being tested in that particular selection experiment. Close orthologues of the host-specific proteins were excluded from further analyses, by performing a blastp search of each ORFs on host proteomes (https://www.uniprot.org/proteomes/UP000001734, https://www.uniprot.org/proteomes/UP000029103, downloaded on 2022.11.24.) and removing each ORFs with higher than 80% sequence similarity. The potential origin of the inserts was assessed by searching the Nanopore contigs within the NCBI Prokaryotic RefSeq Genomes database^87^ using blastn from NCBI BLAST v2.12.0 with default settings and resolving taxids to hierarchical classifications using R^88^ and the taxizedb package^89–91^.

### Catalogue of mobile ARGs

A mobile gene catalogue (that is, a database of recently transferred DNA sequences between bacterial species^92^) was created previously^28^. Briefly, 1377 genomes of diverse human-related bacterial species from Integrated Microbial Genomes and Microbiomes database^92^ and 1417 genomes of Gram negative ESKAPE pathogens from NCBI RefSeq database were downloaded. Using NCBI blastn 2.10.1+,^85^ we searched the nucleotide sequences shared between genomes belonging to different species. The parameters for filtering the NCBI blastn 2.10.1+ blast results were the following: minimum percentage of identity, 99%; minimum alignment length, 500; maximum alignment length, 20,000. Then, to generate the mobile gene catalogue, we compared them with the merged CARD 3.1.0^37^ and ResFinder (d48a0fe) databases^38^ using diamond v2.0.4.142^78^. Natural plasmid sequences were identified by downloading 27,939 complete plasmid sequences from the PLSDB database (version 2020-11-19,^93^). Then, the representative sequences of the isolated 114 ARG clusters were BLASTN searched both in the mobile gene catalogue and in natural plasmid sequences with an identity and coverage threshold of 90%. Those ARGs were considered as mobile which were present in the mobile gene catalogue and/or in natural plasmid sequences.

### Detecting ARGs present in human-associated microbiome and human pathogens

To identify close homologs of the ARGs discovered in our functional metagenomic screens, we utilized the Global Microbial Gene Catalogue (GMGCv1)^94^. This extensive, non-redundant database comprises over 2.3 billion unigenes, derived from more than 13,000 metagenomes across 14 major habitats, and includes detailed phylogenetic origin information. We applied BLASTN^95^ search to compare the nucleotide sequences of the ORFs from our screens against all unigenes in the GMGCv1, using a stringent identity and coverage threshold of 90%. ARGs were considered human body-associated if they showed sequence homology to unigenes present in at least 5 samples in at least one of the following environments: human gut, oral cavity, skin, nose, blood plasma, or vagina. To further investigate the association of the detected ARGs with human pathogens, we analyzed i) their presence in the clinical metagenomic library, and ii) their phylogenetic relationships to pathogens, specifically focusing on ESKAPE pathogens (*Enterococcus faecium, Staphylococcus aureus, Klebsiella pneumoniae, Acinetobacter baumannii, Pseudomonas aeruginosa, Enterobacter spp*) and those listed in the WHO priority lists (*Acinetobacter baumannii, Pseudomonas aeruginosa, Enterobacteriaceae, Enterococcus faecium, Staphylococcus aureus, Helicobacter pylori, Campylobacter, Salmonella, Neisseria gonorrhoeae, Streptococcus pneumoniae, Haemophilus influenzae,* and *Shigella*) by leveraging species information metadata from the GMGCv1 database for each BLASTN hit.

### Detecting ARGs across E. coli phylogroups, host species types and geographic regions

Host type, geographic location and phylogroup were determined for a dataset of 16,272 E. coli genomes in previous work^36^. In brief, the initial complete dataset of 26,881 E. coli genomes was retrieved from the NCBI RefSeq database in February 2022, and filtered for genomes with complete metadata. Clermont phylogrouping^96^ was performed in silico using the EzClermont command-line tool^97^, while host and location metadata were retrieved and categorised using the Bio.Entrez utilities from Biopython v1.77. All genomes were sorted into the following host species categories: ‘Human’, ‘Agricultural/Domestic animals’, and ‘Wild animals’. This was achieved using regular expressions constructed by manually reviewing text in the ‘host’ field of the biosample data for each accession number. Geographic locations were split into 20 subregions according to Natural Earth data^98^. A local blastp search was performed for this collection of *E. coli* genomes against a database of the predicted ARG ORFs identified in functional metagenomic screens, using default parameters. ARGs with both 90% amino acid identity and 90% query coverage per subject, and present in no more than 10% of the examined *E. coli* genomes were analyzed further.

### DIvERGE mutagenesis

We performed deep-scanning mutagenesis in the target genes of moxifloxacin, an established topoisomerase inhibitor. The quinolone resistance-determining regions (QRDR)^25^ of the *gyrA* and *parC* genes were subjected to a single round of mutagenesis using DIvERGE in *E. coli* K-12 MG 1655 and *K. pneumoniae* ATCC 10031. A previously described workflow^18^ was employed with minor modifications. Briefly, cells carrying pORTMAGE311B plasmid (Addgene no. 120418) were inoculated into 2 ml LB medium plus 50 μg/ml kanamycin and were grown at 37°C and continuous shaking (250 RPM) for 12 hours. From this starter culture, 500 μl stationary-phase culture was propagated in 50 ml of the same fresh medium under identical conditions. Induction was initiated at a fixed population density (OD600 =0.4) by adding 50 μl of 1 M *m*-toluic acid (dissolved in 96% ethyl alcohol; Sigma-Aldrich) for 30 to 45 minutes at 37°C. After induction, cells were cooled on ice for 15 minutes. Next, cells were washed three times with sterile ice-cold ultrapure distilled water. Finally, the cell pellet was resuspended in 800 μl sterile ultrapure distilled water and kept on ice until electroporation.

To perform DIvERGE mutagenesis, the corresponding *gyrA* QRDR- and *parC* QRDR-targeting oligonucleotides were equimolarly mixed. 2 μl of the 500 μl oligonucleotide mixture was added to 40 μl electrocompetent cells in 5 parallel samples. The used oligonucleotides are listed in Supplementary Table 10. Following electroporation, the parallel samples were pooled and were suspended into 25 ml fresh LB medium to allow for cell recovery (37°C and 250 RPM). After 60 minutes of recovery period, an additional 25 ml LB medium was added, and cells were grown for an additional 24 hours.

To select clones with reduced susceptibility to moxifloxacin, 500 μl of each mutant cell libraries were spread onto moxifloxacin-containing MHB agar plates. The plates were incubated at 37°C for 48 h. Finally, 20-20 antibiotic resistant clones were selected randomly and analyzed further by capillary sequencing.

### Efflux activity measurements

Measuring the accumulation of the fluorescent Hoechst dye is known as a robust and rapid method for monitoring efflux activity/membrane permeability in bacteria^99^. This method is based on the intracellular accumulation of the fluorescent probe Hoechst 33342 (Bisbenzimide H 33342, Sigma-Aldrich). Cells were grown overnight in MHB then 20 µL of the overnight culture was used to inoculate 2 mL of MHB liquid medium and then the cells were grown to mid-exponential phase (OD_600_ = 0.4–0.6). Bacterial cultures were harvested by centrifugation at 4,500 × *g* for 30 min. Next, cells were washed and resuspended in the buffer, containing 5 m*M* HEPES (pH 7.0) and 5 m*M* glucose. The optical density (OD_600_) of the cell suspensions was adjusted to 0.1, and 0.18 mL of each suspension was transferred to 96-well plates (CellCarrier-96 Black Optically Clear Bottom, supplied by Sigma-Aldrich). Plates were incubated in a Synergy H1 microplate reader at 37°C, and 25 μM Hoechst 33342 was added to each well. The ancestor strain was treated with an efflux inhibitor agent (Phenylalanine-arginine β-naphthylamide (PAβN)) that served as a positive control. The OD_600_ and fluorescence curves were recorded for 2 h with 75 s delays between readings and 2.5 min reading intervals. Fluorescence reading was performed from the top of the wells using excitation and emission filters of 355 and 460 nm, respectively. To estimate changes in efflux activity, we employed a two-step process: (i) we measured the optical density-normalized fluorescence signal over a fixed timeframe (from 7.5 to 120 min) to monitor the intracellular accumulation of a fluorescent probe, and (ii) we calculated the change in normalized fluorescence signal by dividing the signal at the final timepoint (120 min) by that at the initial timepoint (7.5 min). Relative efflux activity of the tested strains was determined by normalizing the reached raw values to those of their respective ancestral strains, and taking its inverse.

## Acknowledgments

The authors thank Adrienn Kobl for her help with the visual elements of Fig. 1, which was created using BioRender, owning a full licence to publish.

This work was supported by:

The European Research Counsil ERC-2023-ADG 101142626 FutureAntibiotics (CP)

The National Laboratory of Biotechnology Grant 2022-2.1.1-NL-2022-00008 (CP, BK, BP) National Research Development and Innovation Office ‘Élvonal’ Programme KKP 126506 (CP)

National Research, Development and Innovation Office ‘Élvonal’ program KKP KH125616 (BP)

The National Laboratory for Health Security RRF-2.3.1-21-2022-00006 (BP)

The European Union’s Horizon 2020 research and innovation programme under grant agreement no. 739593 (BK, BP, MM)

National Research, Development and Innovation Office grant FK-135245 (BK)

János Bolyai Research Fellowship from the Hungarian Academy of Sciences (BO/352/20) (BK)

New National Excellence Program of the Ministry of Human Capacities (UNKP-20-5-SZTE- 654 and UNKP-21- 5-SZTE-579) (BK)

Proof of Concept grant of the Eötvös Loránd Research Network (ELKH-PoC-2022-034) (B K)

The New National Excellence Program of the Ministry for Culture and Innovation - ÚNKP- 22-2-SZTE-220 and ÚNKP-23-3-SZTE-272 (MSCz)

The National Academy of Scientist Education under the sponsorship of the Hungarian Ministry of Culture and Innovation (VI/1697-4/2022/FÁFIN) (MSCz)

The National Research, Development and Innovation Office, Hungary (NKFIH) grant PD, grant number 131839 (EA)

The National Research, Development and Innovation Office, Hungary (NKFIH) grant FK- 142312 (MM)

The National Research, Development and Innovation Office, Hungary (NKFIH) KIM NKFIA TKP-2021-EGA-05 (SJ)

The National Research, Development and Innovation Office, Hungary (NKFIH) KIM NKFIA 2022-2.1.1-NL-2022-00005 (SJ) H2020-WIDESPREA-01-2016-2017-TeamingPhase2, GA:739593-HCEMM (SJ)

The National Research, Development and Innovation Office, Hungary (NKFIH) grant FK- 131961 (SJ)

The New National Excellence Program of the Ministry of Human Capacities Bolyai +, ÚNKP-22-5-SZTE-578-Bolyai+ (SJ)

The Janos Bolyai Research Fellowship from the Hungarian Academy of Sciences bo_656_20 (SJ)

The Lister Institute for Preventative Medicine (SvH)

The National Research, Development, and Innovation Office (PharmaLab, RRF-2.3.1-21-2022-00015 and TKP-31-8/PALY-2021) (LH)

## Author contributions

Conceptualization: LD, MSCz, PSz, ZF, BK, BP, CP

Methodology: LD, MSCz, PSz, ZF, DB, GG, SJ, ADunai, MSz, TSári, TStirling, BMV, EA, CC, MM, MZsE, GJ, SvH, EP, LP, LH, BK, BP, CP

Software: GG, TStirling, BMV, EA, CC, MM, GJ, SvH, EP

Validation: LD, MSCz, PSz, ZF, DB, GG, SJ, ADunai, MSz, TSári, TStirling, BMV, EA, CC, MM, MZsE, GJ, SvH, EP, LP

Formal analysis: LD, MSCz, PSz, ZF, DB, EM, TV, LS, BDV, GG, SJ, AD, MSz, TStirling, BMV, KK, EA, CC, MM, GJ, SvH, EP

Investigation: LD, MSCz, PSz, DB, EM, TV, LS, BDV, SJ, ADunai, MSz Resources: ADaraba, TSári, MZsE, LP, LH, BK, BP, CP

Data Curation: LD, MSCz, PSz, ZF, GG, TStirling, BMV, EA, CC, MM, GJ, SvH, EP Writing - Original Draft: LD, MSCz, PSz, ZF, DB, GG, SJ, ADaraba, MSz, TStirling,

BMV, EA, MM, BK, BP, CP

Writing - Review & Editing: LD, MSCz, PSz, ZF, ADaraba, BK, BP, CP Visualization: LD, MSCz, PSz, ZF, GG, TStirling, BMV, EA, CC, MM Supervision: LH, BK, BP, CP

Project administration: ADaraba, BK, BP, CP Funding acquisition: BK, BP, CP

## Data and materials availability

All data are available in the main text or the supplementary information. Illumina reads and Nanopore contigs for this study have been deposited in the European Nucleotide Archive (ENA) at EMBL-EBI under accession number PRJEB63210 (https://www.ebi.ac.uk/ena/browser/view/PRJEB63210).

## Supplementary Information

Supplementary Notes 1-4

Description of Supplementary Tables 1-12 (in Auxiliary Supplementary File) and Supplementary Data 1

---

**Extended Data Fig. 1.**
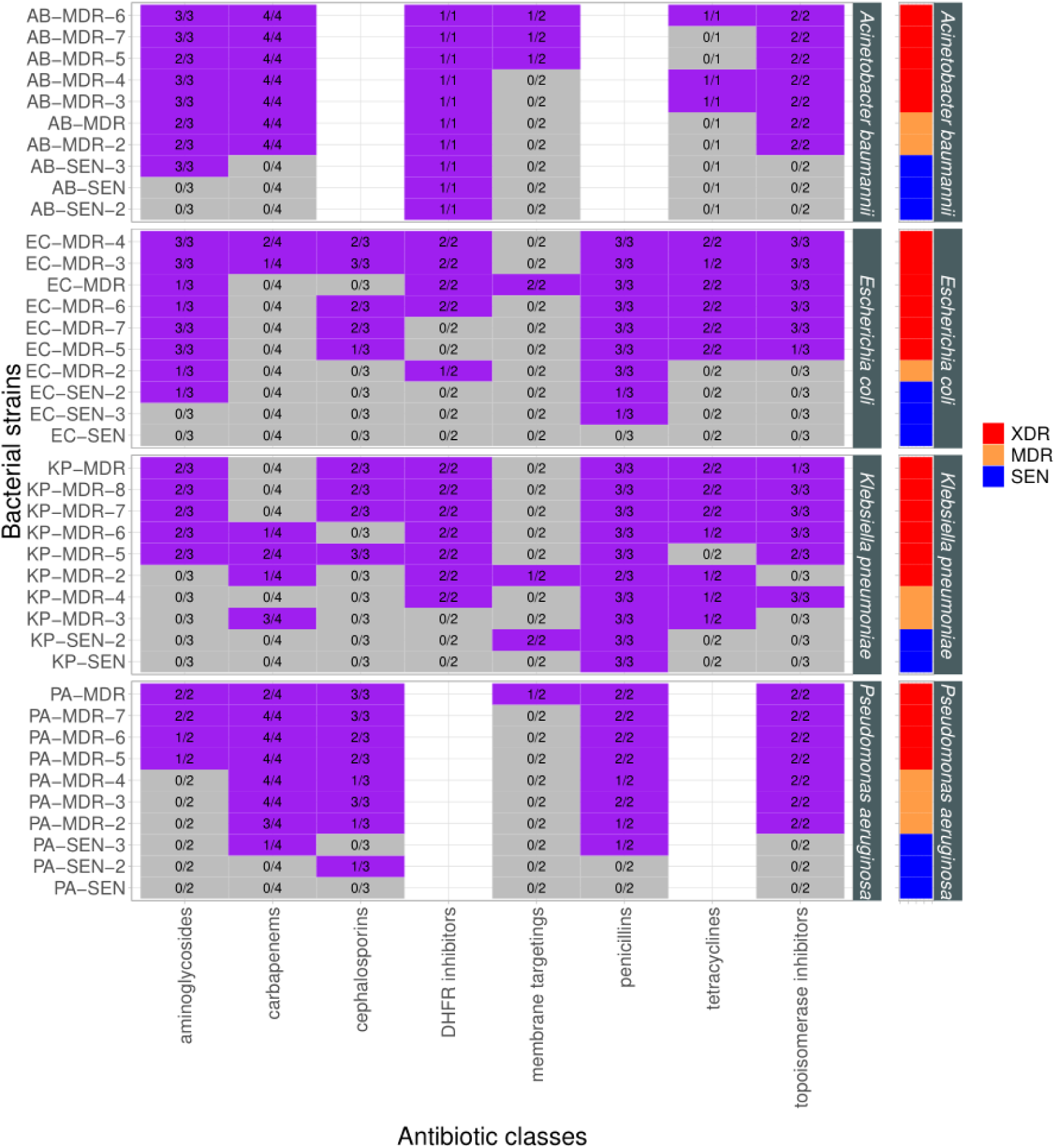
The heatmap shows the resistance profile of 40 bacterial strains against 8 different antibiotic classes. Strains resistant and sensitive to a given antibiotic class (based on clinical breakpoints) are represented by purple and gray, respectively. The numbers correspond to the number of drugs a strain is resistant to out of the number of drugs tested. The right panel classifies the strains based on their resistance profile as sensitive (SEN), multidrug-resistant (MDR), or extremely drug-resistant (XDR). Strains are classified as MDR, if being resistant to 3 or 4 antibiotic classes and XDR if being resistant to 5 or more antibiotic classes, respectively^101^. For abbreviations, see Supplementary Table 1 and Supplementary Table 2.

**Extended Data Fig. 2.**
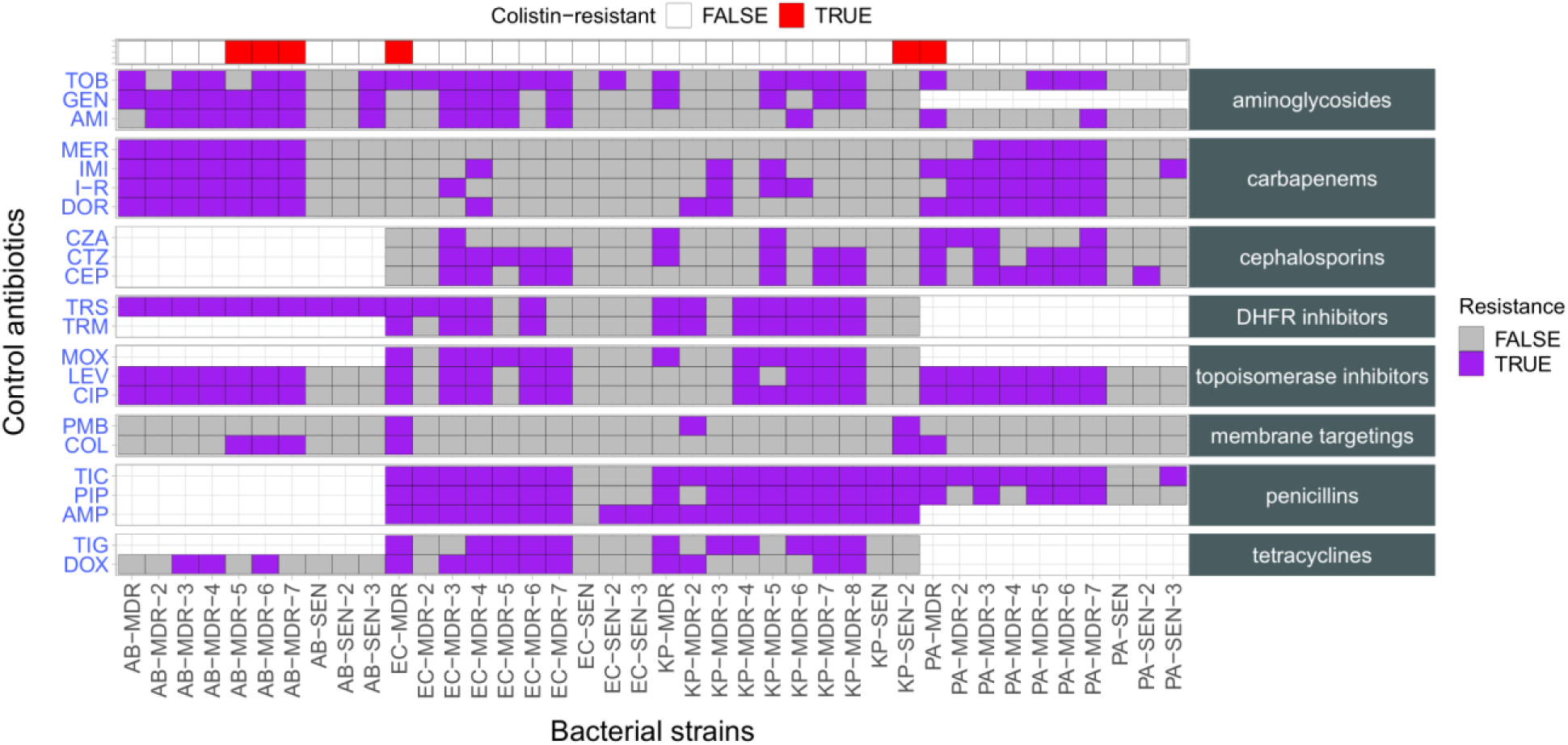
The heatmap shows the resistance profile of 40 bacterial strains against control drugs. Antibiotics are grouped by the antibiotic classes they belong to. Bacterials strains are ordered by the fraction of resistance against control antibiotics. Strains resistant and sensitive to a given antibiotic are represented by purple and gray, respectively. Resistance is defined as having an MIC value greater than the clinical breakpoint in a given species. Accordingly, missing values indicate the absence of clinical breakpoints in certain combinations of species and antibiotics. Strains that are colistin-resistant are indicated in the top panel. For abbreviations, see Table 1 and Supplementary Table 2.

**Extended Data Fig. 3.**
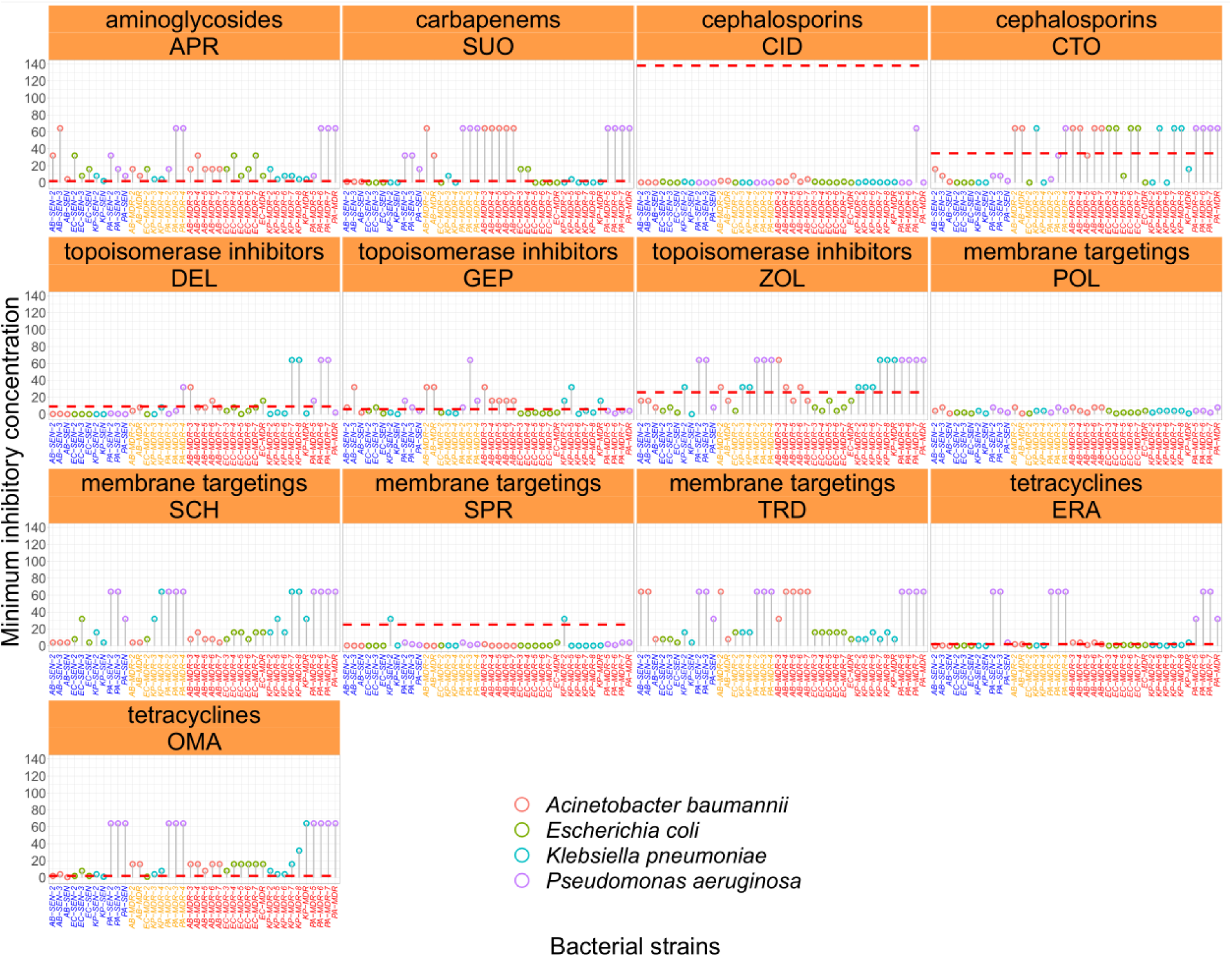
The figure shows the antibiotic susceptibility of 40 strains to ‘recent’ antibiotics. Blue, orange, and red labels indicate sensitive (SEN), multidrug-resistant (MDR), or extremely drug-resistant (XDR) strains, respectively. The color of the points marks the bacterial species. Dashed line indicates the estimated antibiotic specific peak-plasma concentration (see Supplementary Table 1). For abbreviations, see Table 1 and Supplementary Table 2.

**Extended Data Fig. 4.**
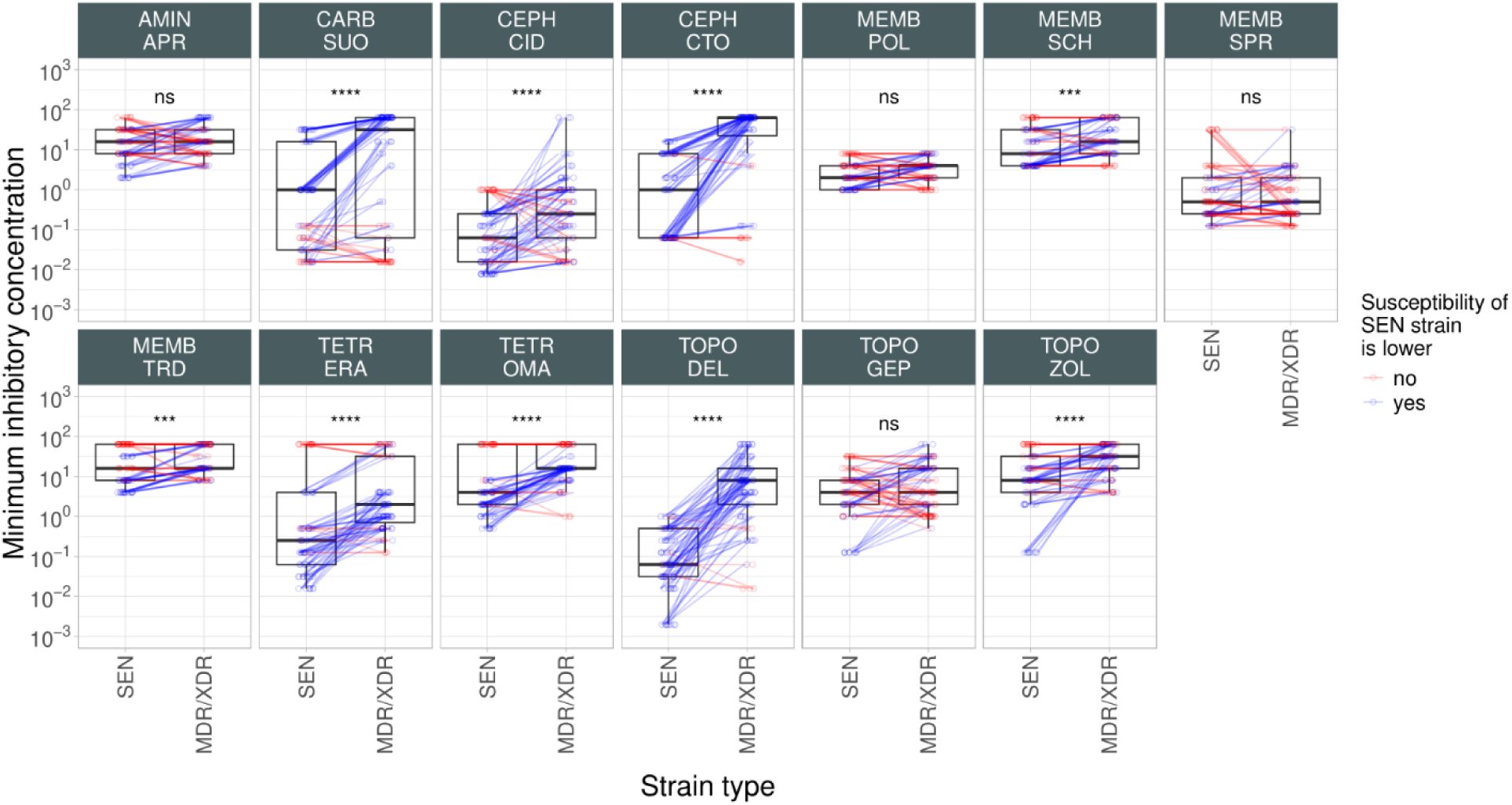
Comparison of minimum inhibitory concentration (MIC) of recent antibiotics for sensitive and multidrug-resistant/extensively drug-resistant strains. The figure shows the MIC values (on a log_10_ scale) of recent antibiotics across all tested bacterial strains. Each vertical panel represents a specific antibiotic class and antibiotic, as indicated at the top. Individual points depict strain-antibiotic pairs, with lines connecting paired data points representing MIC values of one antibiotic for one sensitive (SEN) and one multidrug-resistant/extensively drug-resistant (MDR/XDR) strain for the same species. Blue and red colors indicate cases where the MIC of a single antibiotic for a SEN strain is lower or not than that of the MDR/XDR strain for the same species. Boxplots display the median, first, and third quartiles, with whiskers indicating the 5th and 95th percentiles. Paired Wilcoxon rank sum analysis was performed to assess significant difference in sensitivity between antibiotic sensitive (SEN) and MDR/XDR bacterial strains. ****/*** indicates P < 0.0001/0.001, ns indicates that the P value is non-significant. For antibiotic abbreviations, see Table 1. Antibiotic classes: TOPO (topoisomerase inhibitors), TETR (tetracyclines), AMIN (aminoglycosides), CARB (carbapenems), CEPH (cephalosporins), and MEMB (membrane-targeting antibiotics.

**Extended Data Fig. 5.**
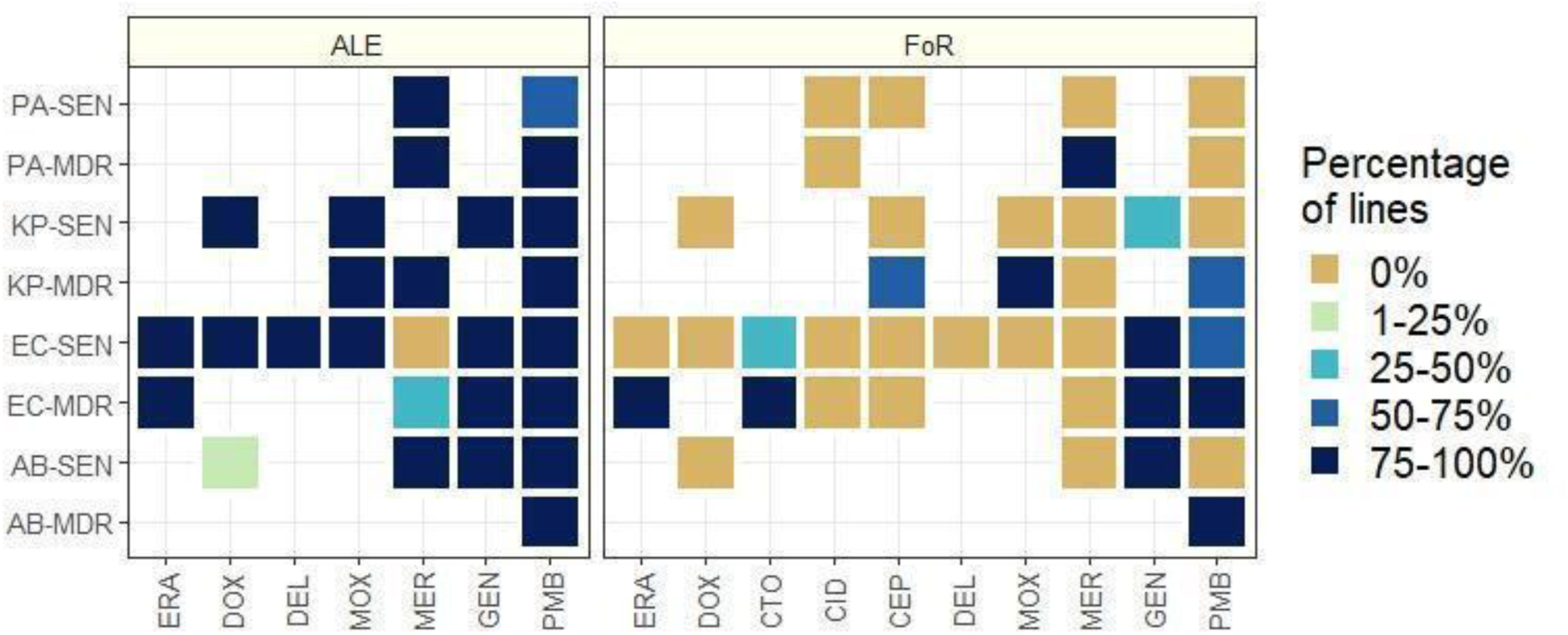
Percentage of mutant lines reaching the clinical breakpoint. The heatmap shows the percentage of lines reaching the clinical breakpoint. Unavailable clinical breakpoints are indicated by white. For the antibiotic abbreviation and clinical breakpoints see Supplementary Table 1, for strain abbreviation see Supplementary Table 2.

**Extended Data Fig. 6.**
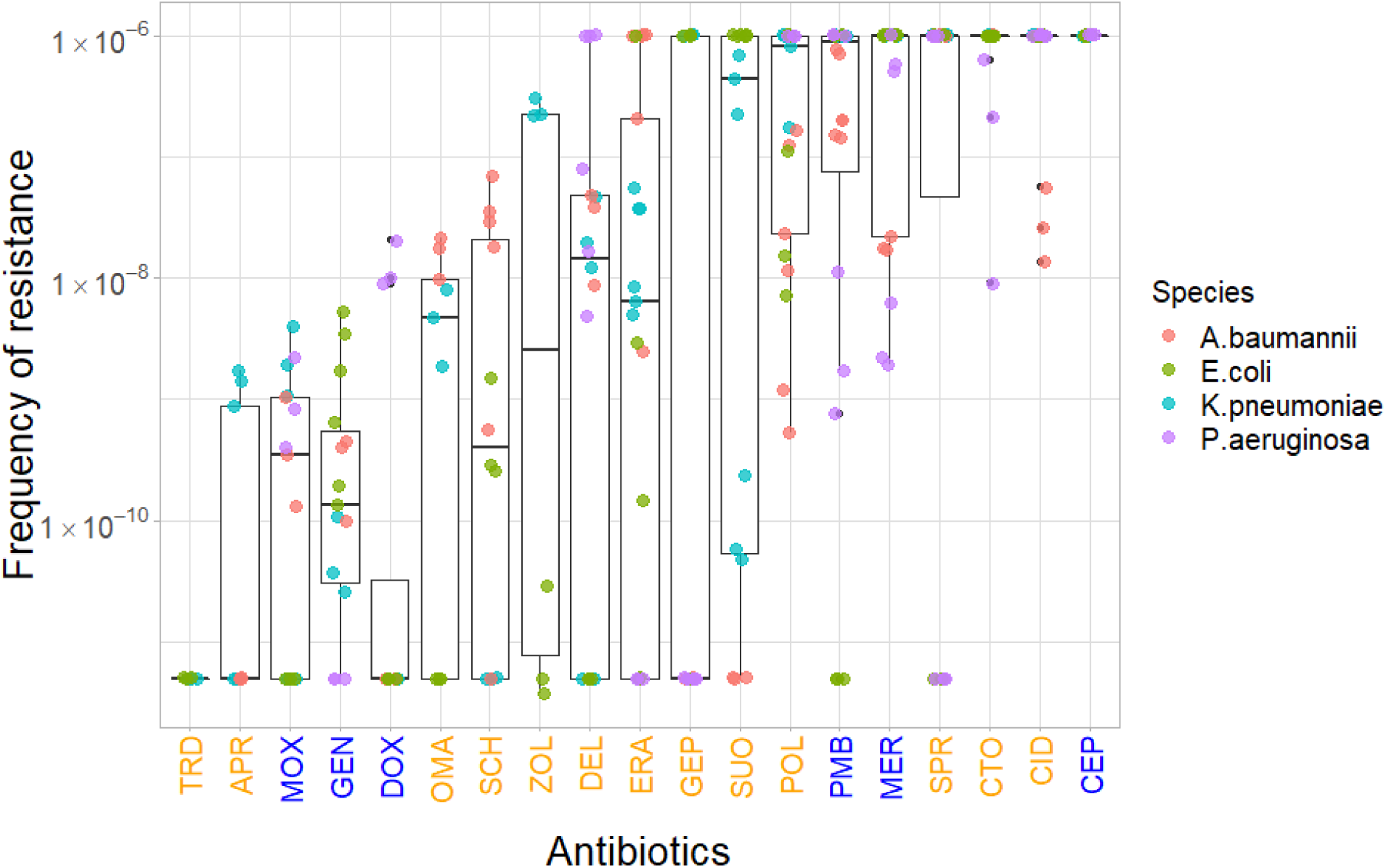
Frequency of resistance of evolved lines adapted to different antibiotics. The frequency of resistance at 8xMIC antibiotic concentrations was calculated for all antibiotics, shown as the number of mutations per cell per generation. Each data point represents a distinct mutant line derived from the frequency of resistance assays (FoR), species are denoted by different colors. The label color on the x-axis shows the generation of the different antibiotics (blue stands for control, orange for recent antibiotics). The boxplots show the median, first, and third quartiles, with whiskers showing the 5th and 95th percentiles. There is highly significant heterogeneity in the frequency of resistance across antibiotics (Kruskal-Wallis test, P < 0.00001), but no statistical difference was found between control and recent antibiotics (Wilcoxon rank-sum-test, P = 0.9) when all antibiotics were considered. For antibiotic abbreviations, see Table 1.

**Extended Data Fig. 7A.**
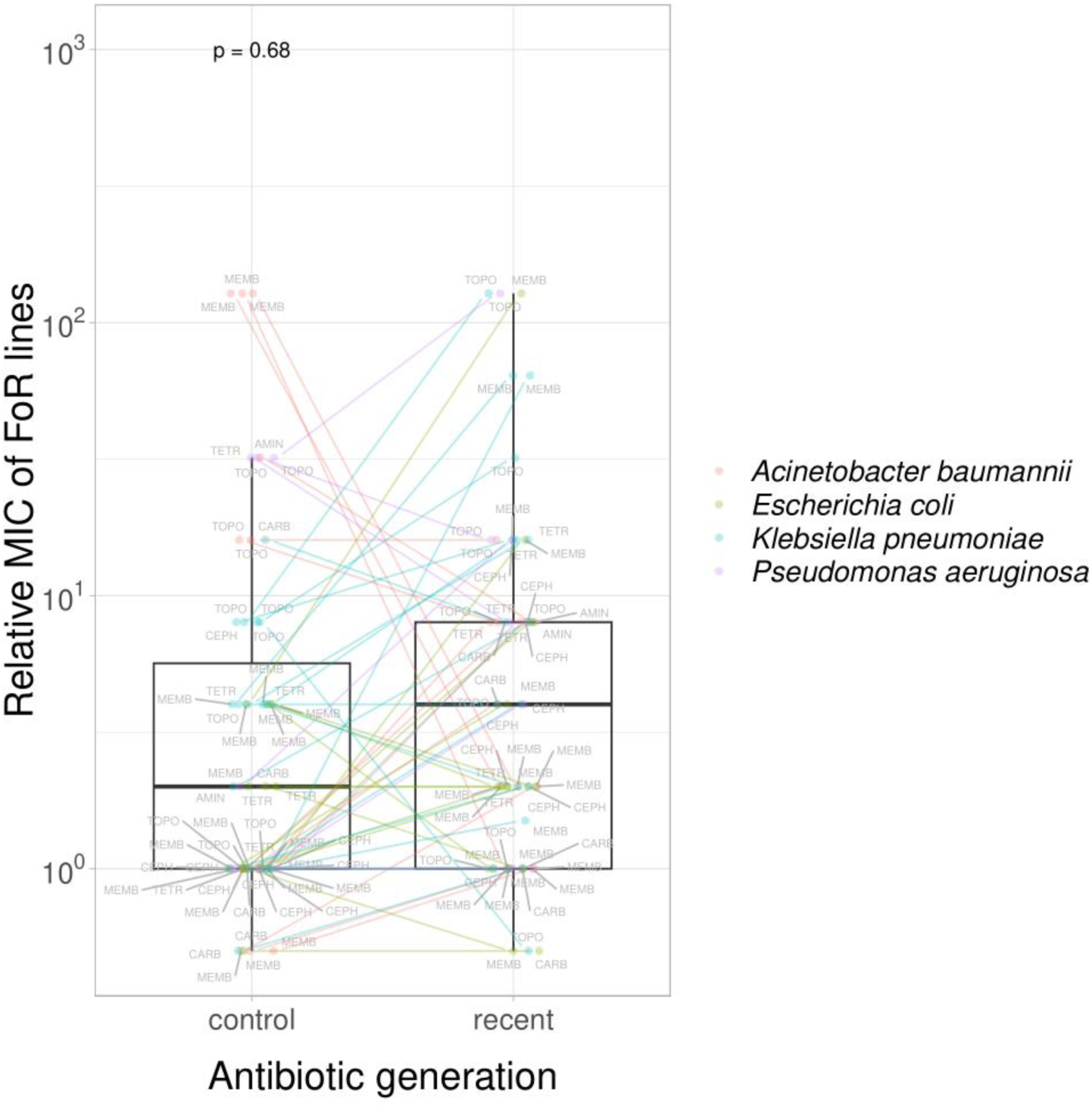
Control and recent antibiotics are equally prone to resistance in frequency-of-resistance assays. The boxplot shows the relative MIC of control and recent antibiotics from frequency-of-resistance (FoR) assay. The lines connect different generations of antibiotic belonging to the same antibiotic class (labels). We found no significant difference between the control and recent antibiotics (paired T-test, P = 0.68).

**Extended Data Fig. 7B.**
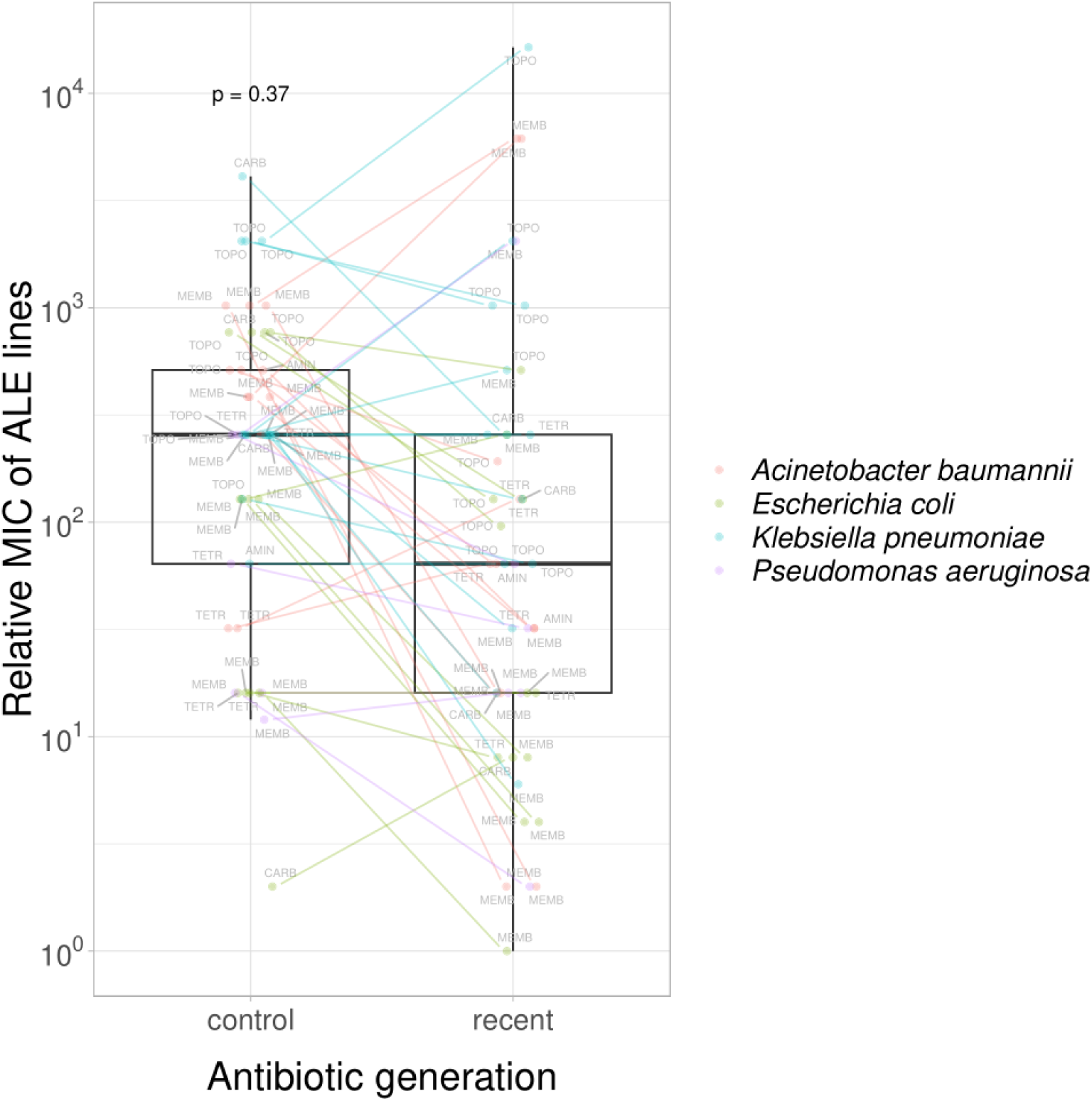
Control and recent antibiotics are equally prone to resistance in adaptive laboratory evolution. The boxplot shows the relative MIC of control and recent antibiotics from adaptive laboratory evolution (ALE). The lines connect different generations of antibiotic belonging to the same antibiotic class (labels). We found no significant difference between the control and recent antibiotics (paired T-test, P = 0.37).

**Extended Data Fig. 8A.**
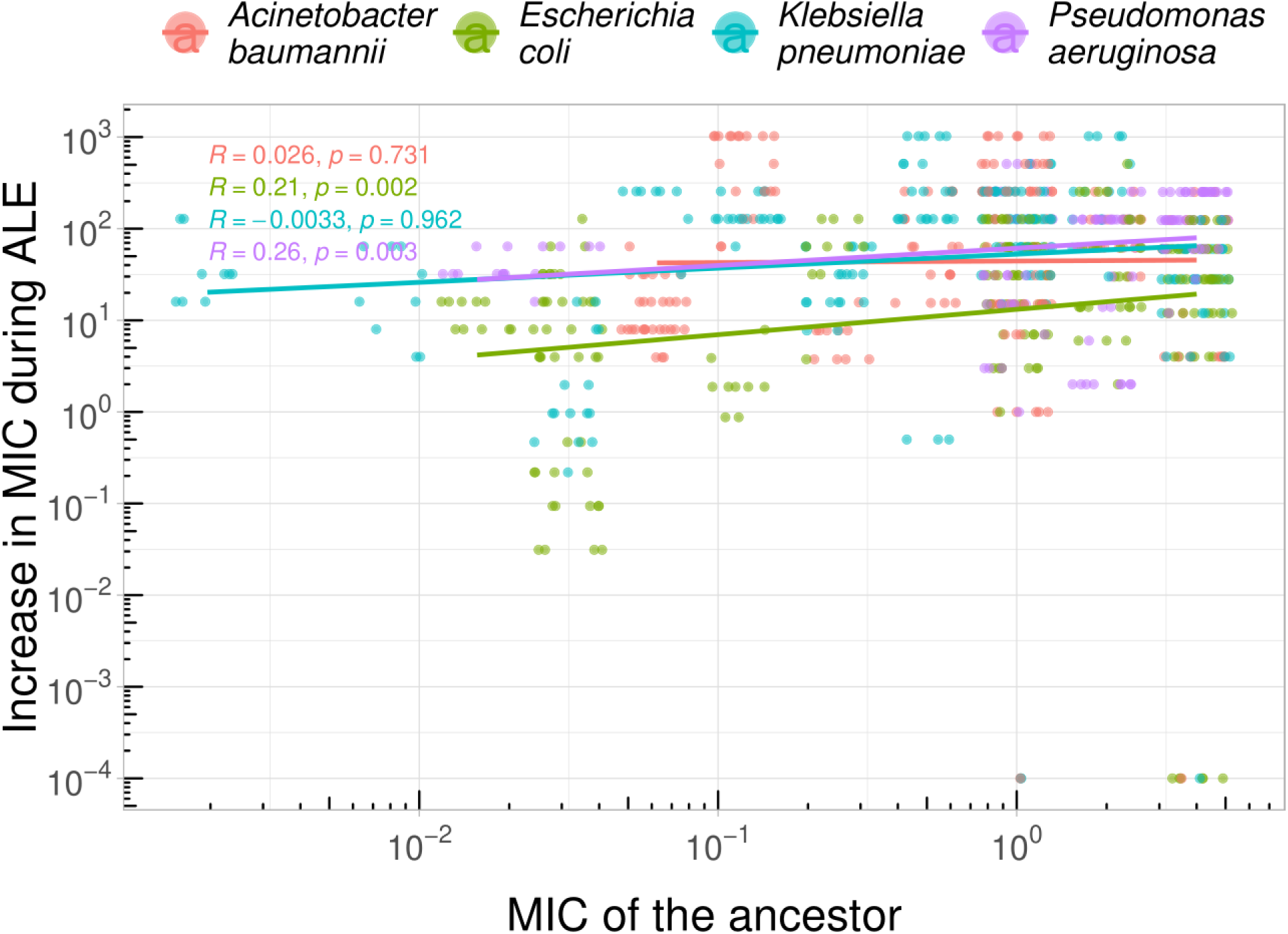
Correlation of the initial MIC and increase in resistance levels during adaptive laboratory evolution. The scatterplot shows the correlation between initial resistance level (MIC of the ancestor) and the increase in MIC (both on log10-scale) during adaptive laboratory evolution (ALE) for the four bacterial species. Increase in MIC was calculated by subtracting the initial MIC from the final MIC. Each point corresponds to an adapted line- antibiotic pair. Spearman’s correlation values between the two variables across all adapted lines of each species are displayed in the figure. Colors indicate the 4 bacterial species studied.

**Extended Data Fig. 8B.**
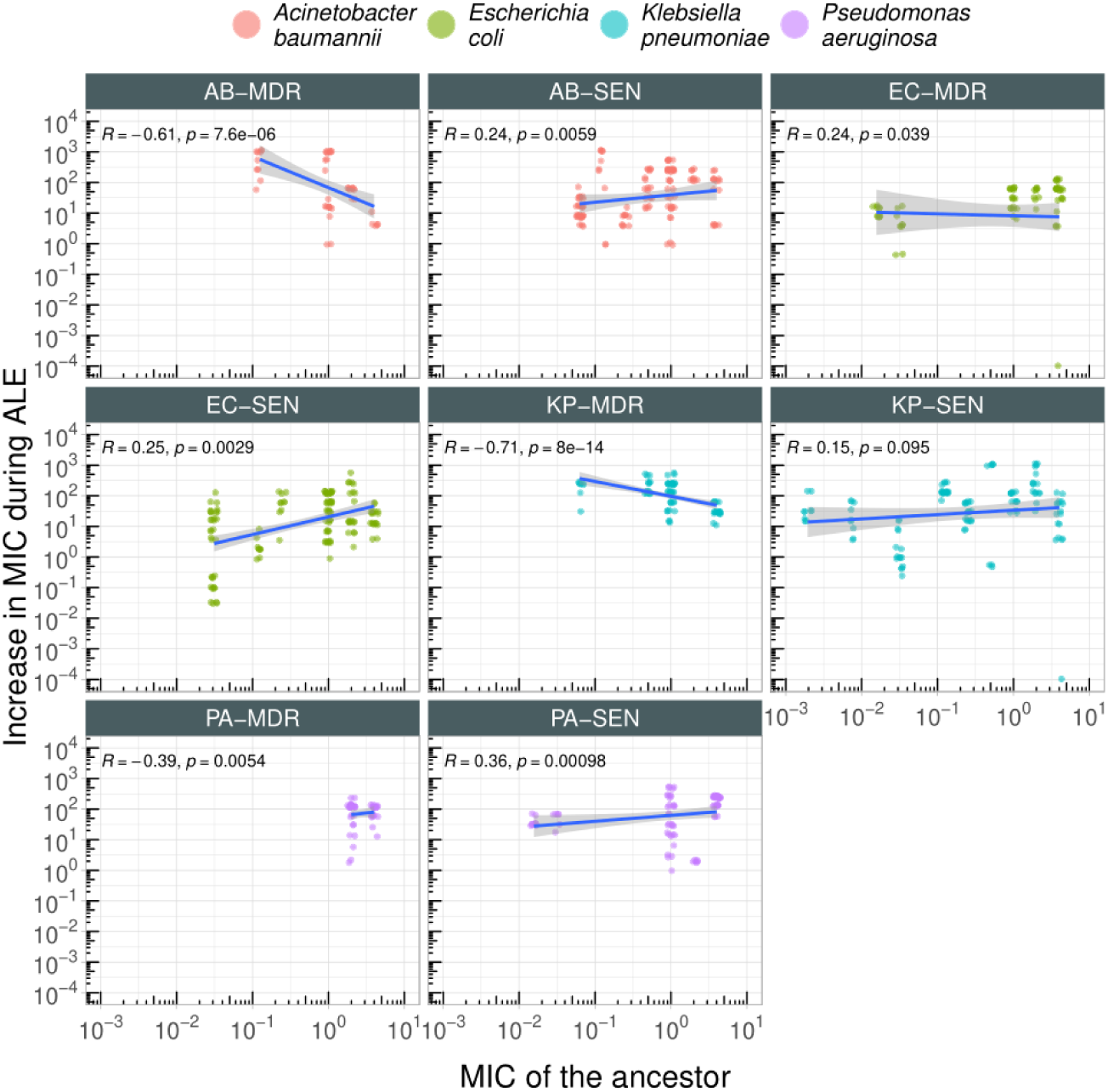
Correlation analysis between the initial MIC and the increase in MIC during adaptive laboratory evolution for all genomic backgrounds. The scatterplots shows the initial MIC and the increase in MIC (both on log10-scale) during adaptive laboratory evolution for each 8 studied bacterial strain (indicated in the top of each panel). Increase in MIC was calculated by subtracting the initial MIC from the final MIC. Each point corresponds to an adapted line- antibiotic pair. Spearman’s correlation coeffiicents and p-values are indicated within each panel. Colors indicate the 4 bacterial species studied. For abbreviations, see Supplementary Table 2.

**Extended Data Fig. 9.**
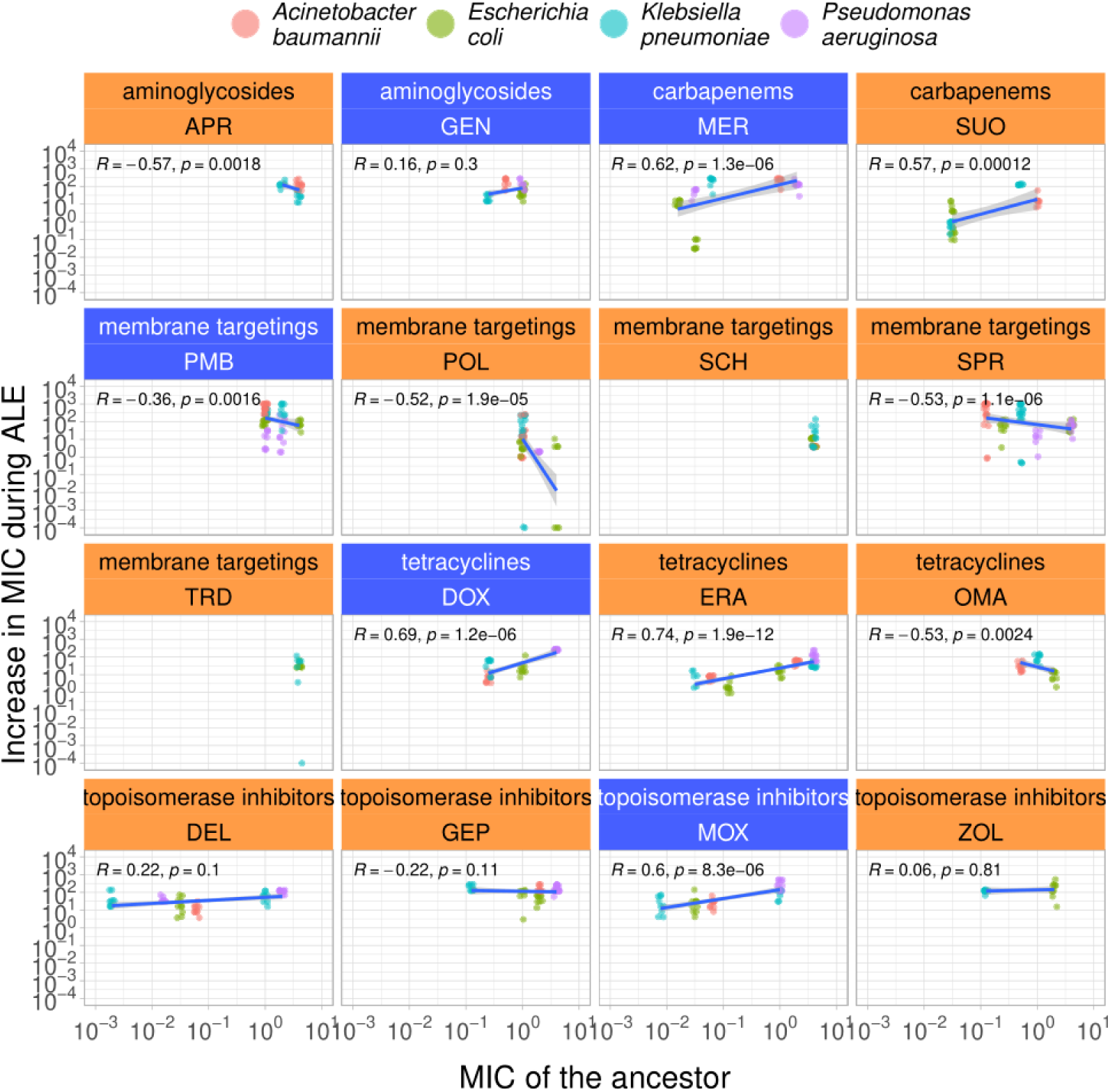
Correlation analysis between the initial MIC and the increase in MIC during adaptive laboratory evolution for all tested antibiotics. The scatterplots shows the initial MIC and the increase in MIC (both on log10-scale) during ALE for all antibiotic studied (indicated in the top of each panel). Increase in MIC was calculated by subtracting the initial MIC from the final MIC. Each point corresponds to an adapted line-antibiotic pair. Spearman’s correlation coefficients and p-values are indicated within each panel (absence of values in certain panels is due to the lack of variability in the initial MIC). Colors indicate the 4 bacterial species studied. For abbreviations, see Table 1.

**Extended Data Fig. 10.**
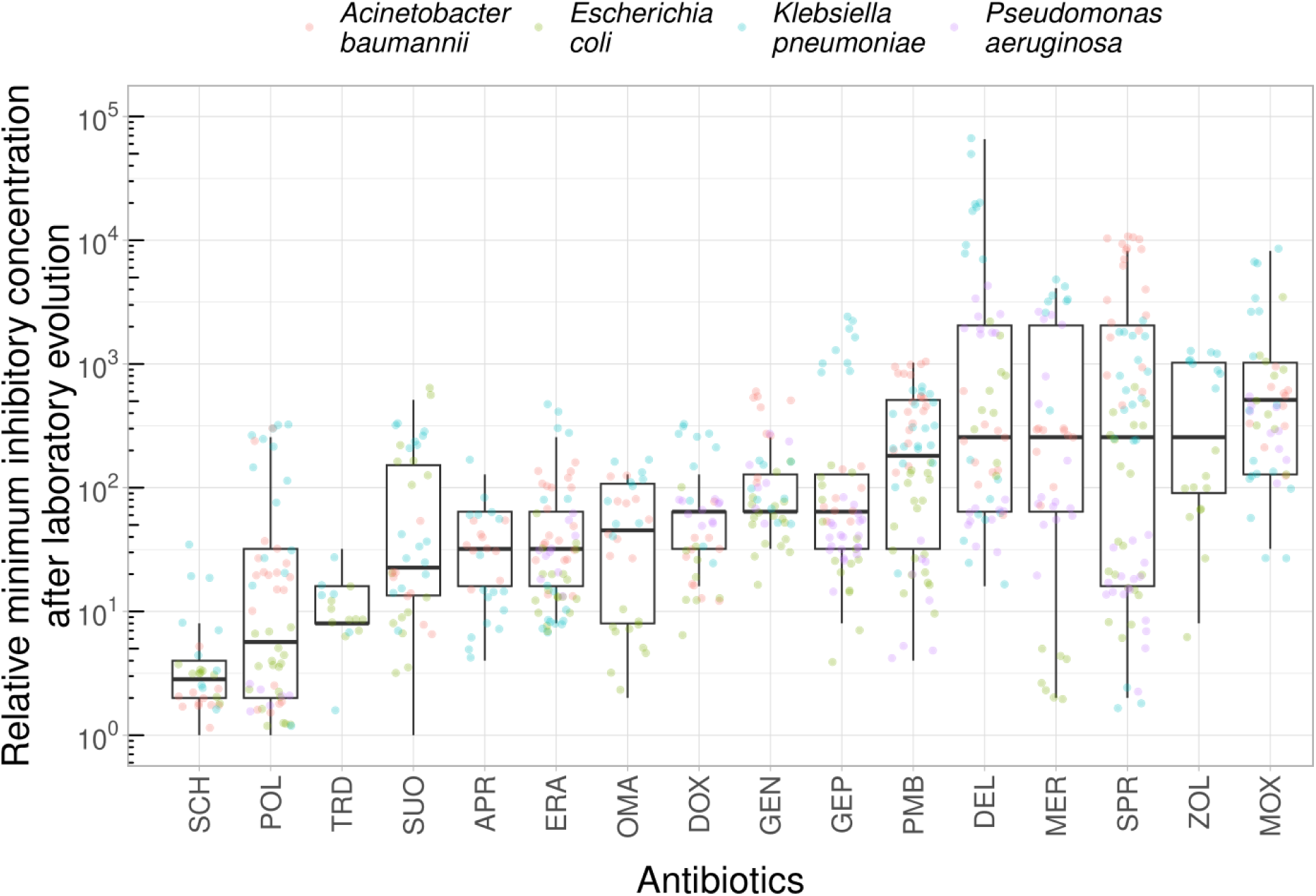
Relative MIC of laboratory evolved lines across all antibiotics. Relative MIC is the MIC of the evolved line divided by the MIC of the corresponding ancestor. Each point is a laboratory evolved line from ALE, the colors indicate the bacterial species. The boxplots show the median, first and third quartiles, with whiskers showing the 5th and 95th percentiles. There is a highly significant heterogeneity in relative MIC across antibiotics (Kruskal-Wallis chi-squared = 281.03, df = 15, P < 2.2e-16). For antibiotic abbreviations, see Table 1.

**Extended Data Fig. 11.**
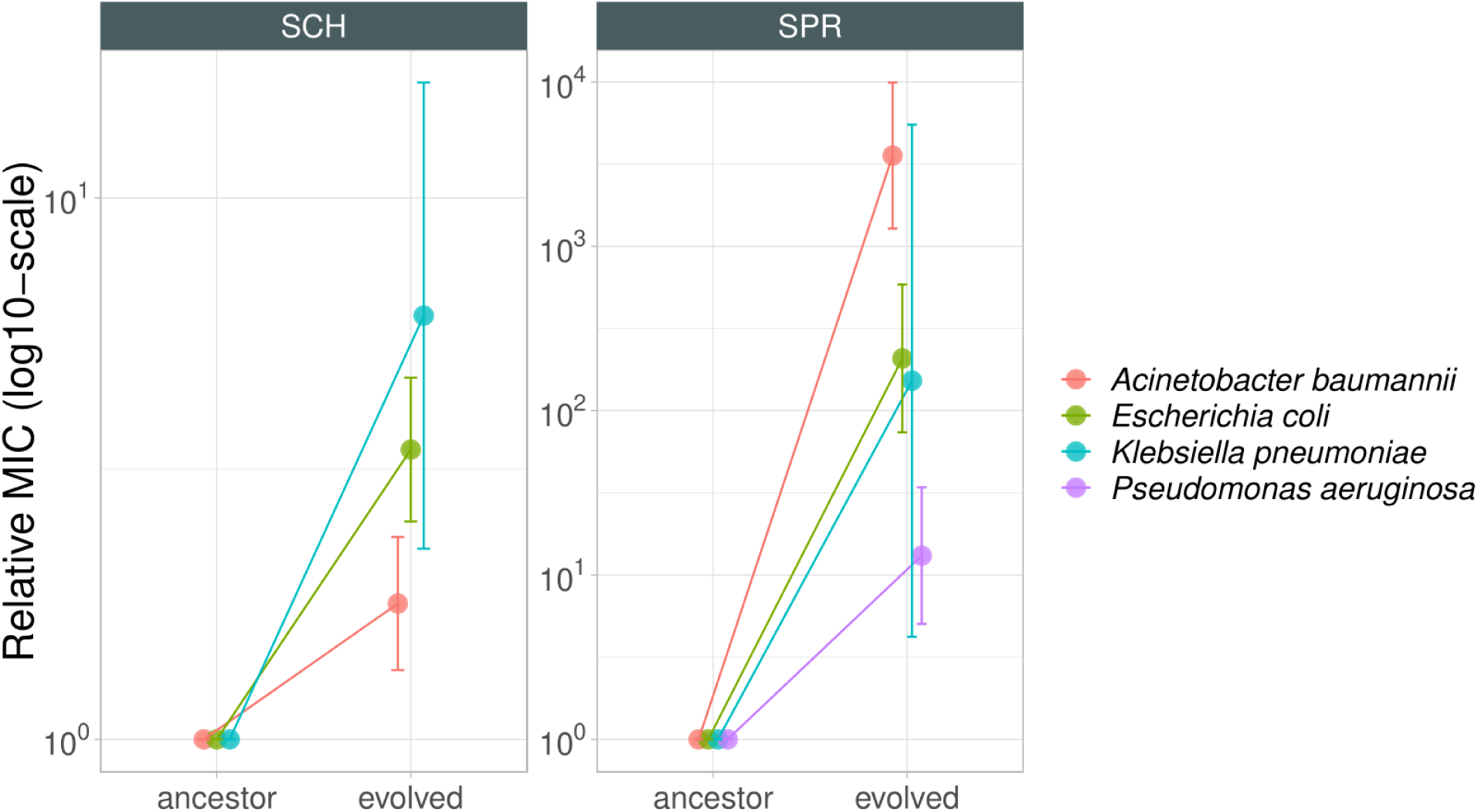
Species-specific differences in resistance evolution. The figure shows the mean relative MIC of lines adapted against SCH7979 (SCH, left panel) and SPR-206 (SPR, right panel), based on at least 6 parallel evolved lines derived from initially antibiotic sensitive (SEN) genetic background. The color of the data points represents the bacterial species. Error bars represent the standard deviation of the relative MIC of all lines per species derived from ALE. We found significant heterogeneity in the relative MIC reached in the evolved lines across the four species (Kruskal-Wallis test, P < 0.01 for both antibiotics). Post-hoc comparison (Dunn’s test) revealed significant differences in relative MIC for SCH7979 between *A. baumannii* and either *E. coli* or *K. pneumoniae* adapted strains (P < 0.05), whereas the difference between *E. coli* and *K. pneumoniae* was not significant (P = 0.58). In case of SPR-206, Dunn’s test revealed significant differences in relative MIC between *A. baumannii* and any other species (P < 0.05), whereas the difference between other pairs of species were not significant (P > 0.05).

**Extended Data Fig. 12.**
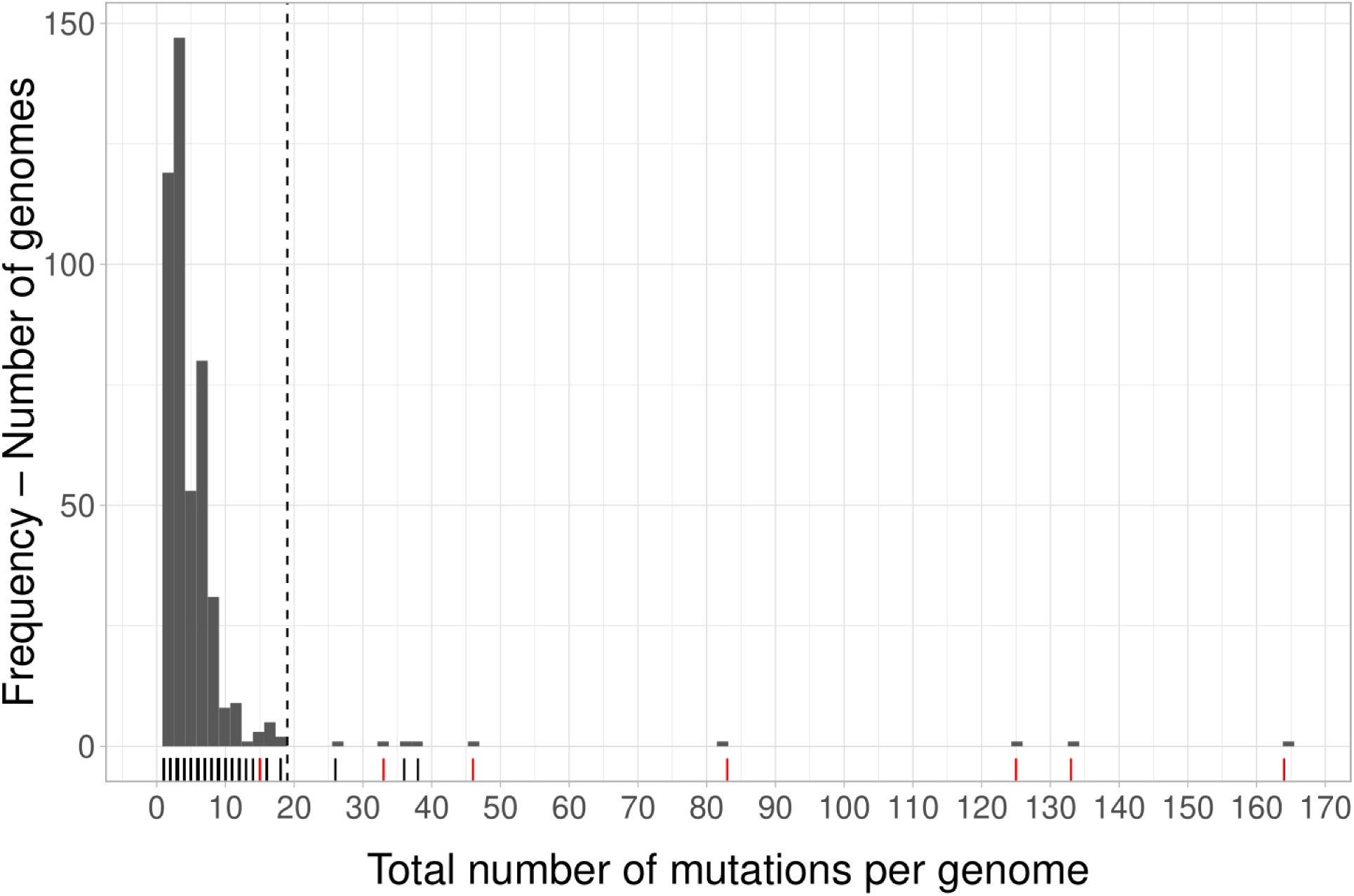
The histogram shows the distribution of the total number of mutations that has accumulated during the course of laboratory evolution. Resistant lineages derived from ALE (n=332) and FoR (n=135) assays are depicted here. 10 resistant lineages accumulated exceptionally large numbers of mutations (>18, see Methods), 6 of which accumulated mutations in methyl-directed mismatch repair (*mutL, mutS* or *mutY*). For these two reasons these lineages are likely to be hypermutator lines, and were excluded from further analysis.

**Extended Data Fig. 13.**
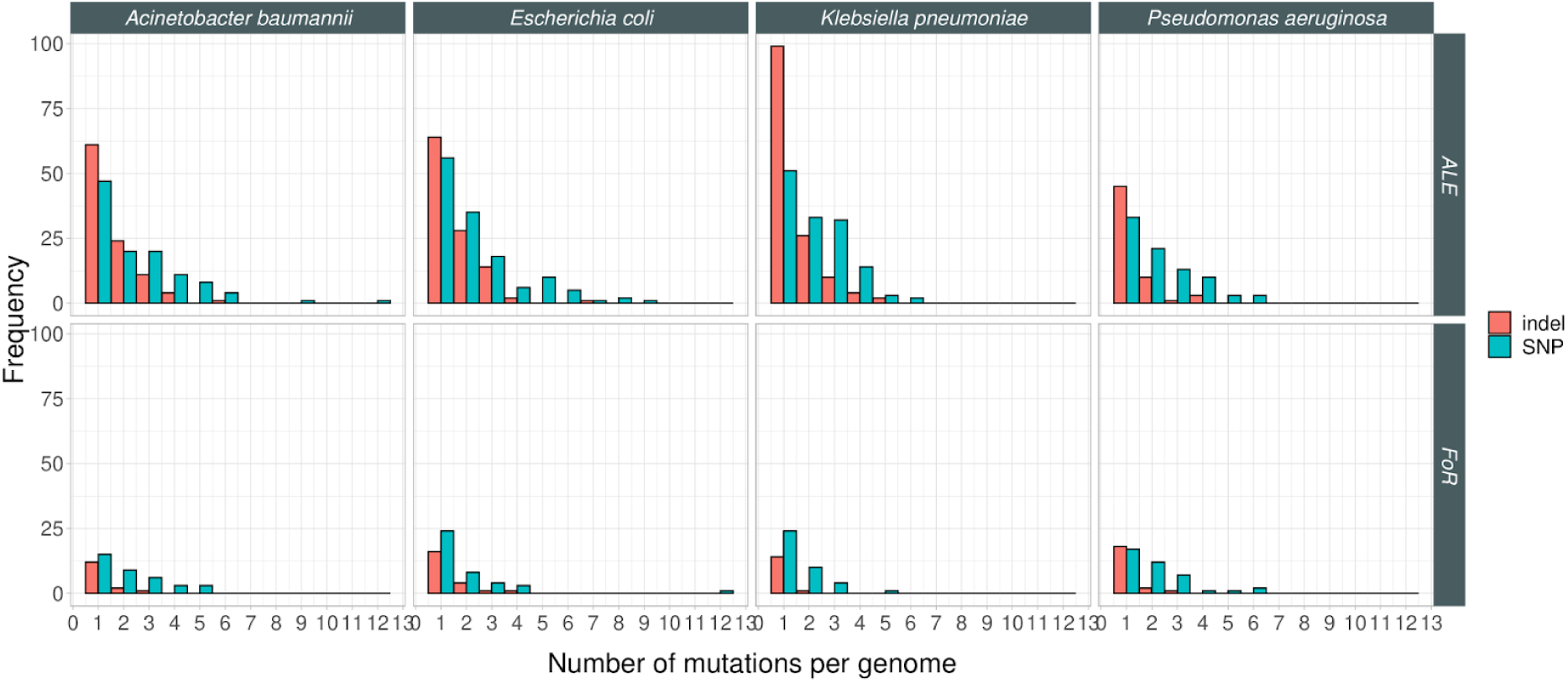
Distribution of SNPs and indels in resistant lines across four bacterial species. Red and green colors denote short indels and SNPs, respectively. As expected, lines derived from FoR have accumulated fewer mutations (Wilcoxon rank sum test, P < 2.2e-16), when compared to lines derived from ALE.

**Extended Data Fig. 14.**
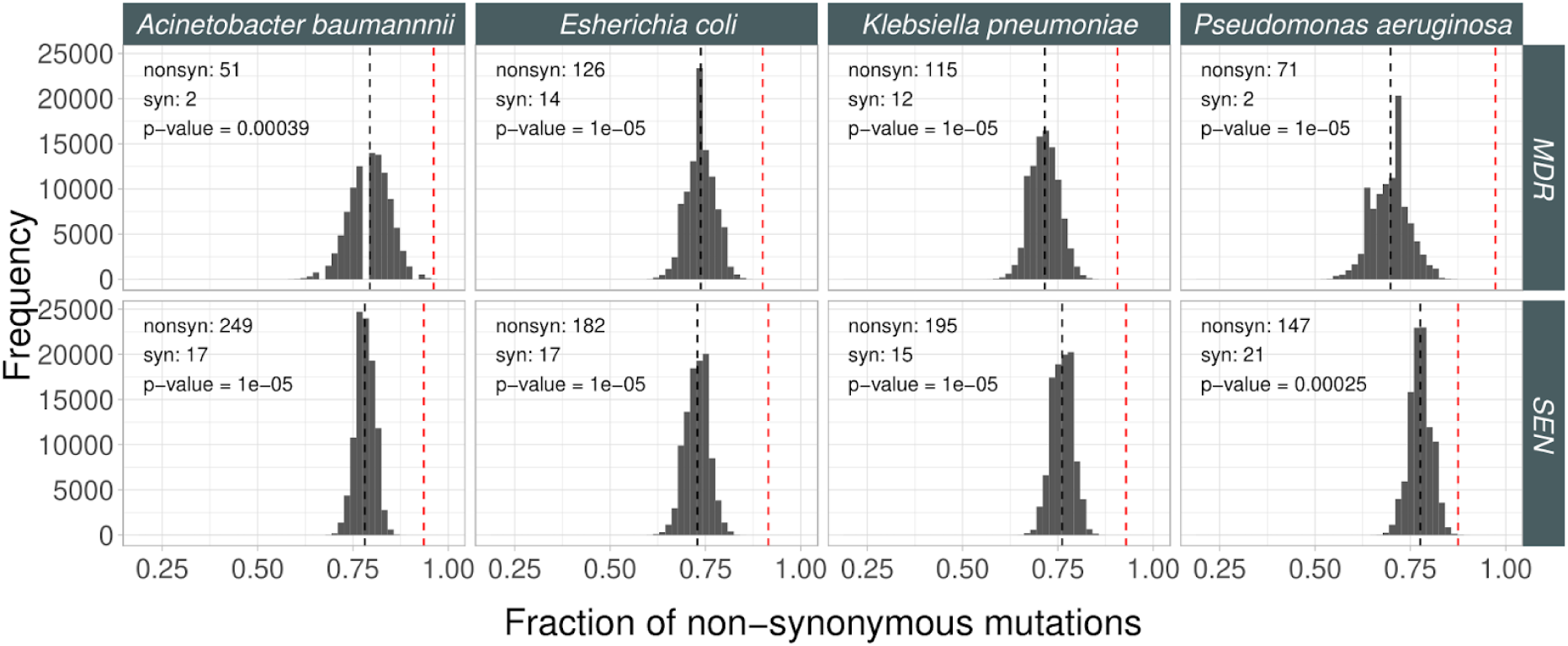
Fraction of non-synonymous mutations. To assess the signatures of adaptive evolution in our genomic samples from ALE and FoR assay, we tested whether the fraction of non-synonymous mutational events within all SNPs in the coding region was higher than expected based on a purely neutral model of evolution using an established method^101^. For each bacterial strain background, we identified all SNPs in the coding regions, counted the number of non-synonymous (*nonsyn*) and synonymous (*syn*) ones and calculated the observed fraction of non-synonymous mutations (red dashed line) as follows: nonsyn / nonsyn + syn, where nonsyn and syn are the number of non-synonymous and synonymous mutations, respectively. Next, we randomly generated the same number of SNPs at random coding positions along the genome as observed in the mutation dataset. We repeated this step 5000 times, then plotted the fraction of non-synonymous mutations as histograms for each species (columns) and strain type (rows). Next, we calculated the probability (P-value) that the fraction of nonsynonymous mutations was equal to or higher in the real data than that of in the randomly generated one.

**Extended Data Fig. 15.**
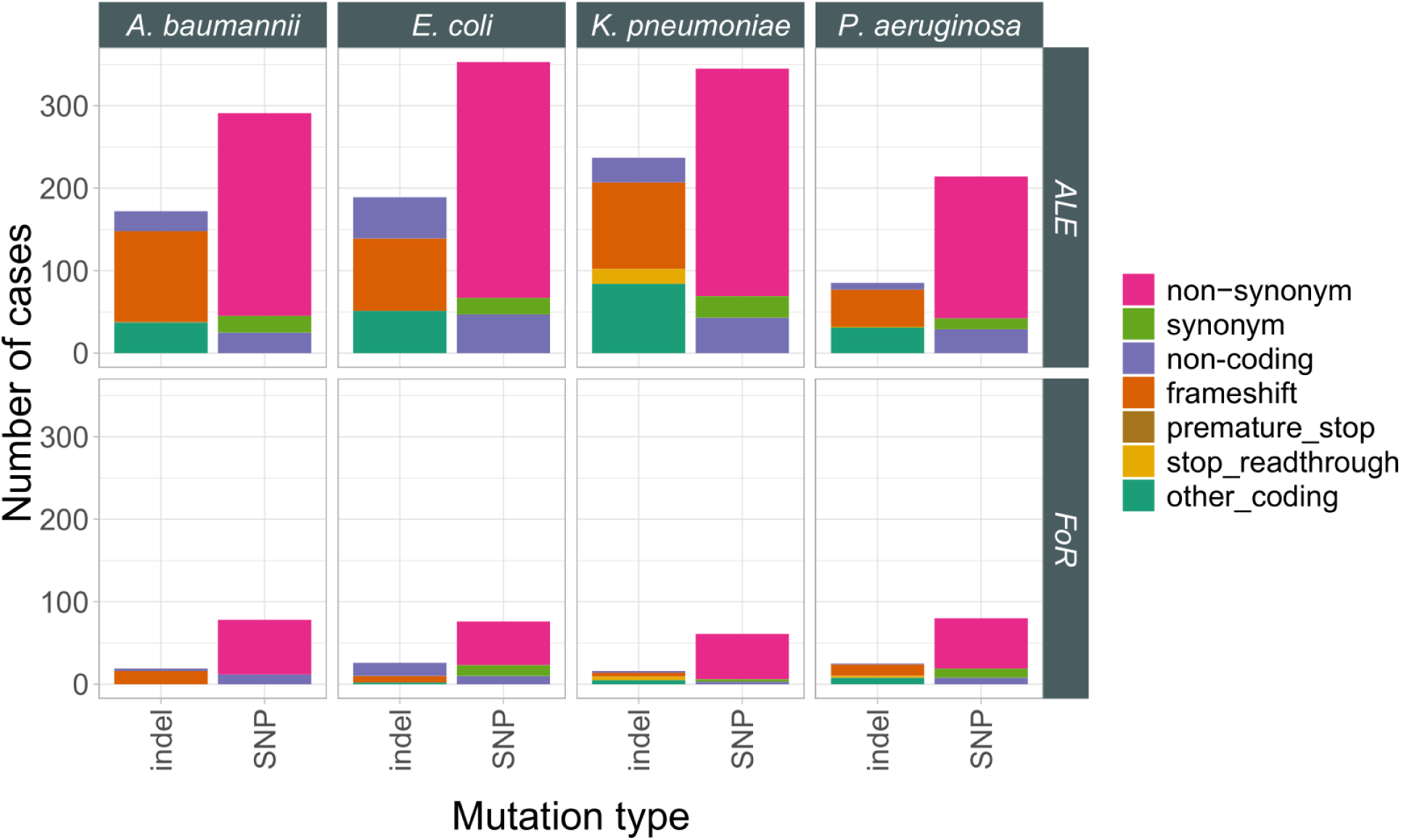
Distribution of different mutational events. Top and bottom row correspond to the adapted lines originating for laboratory evolution (ALE) and frequency of resistance (FoR) assays, respectively.

**Extended Data Fig. 16.**
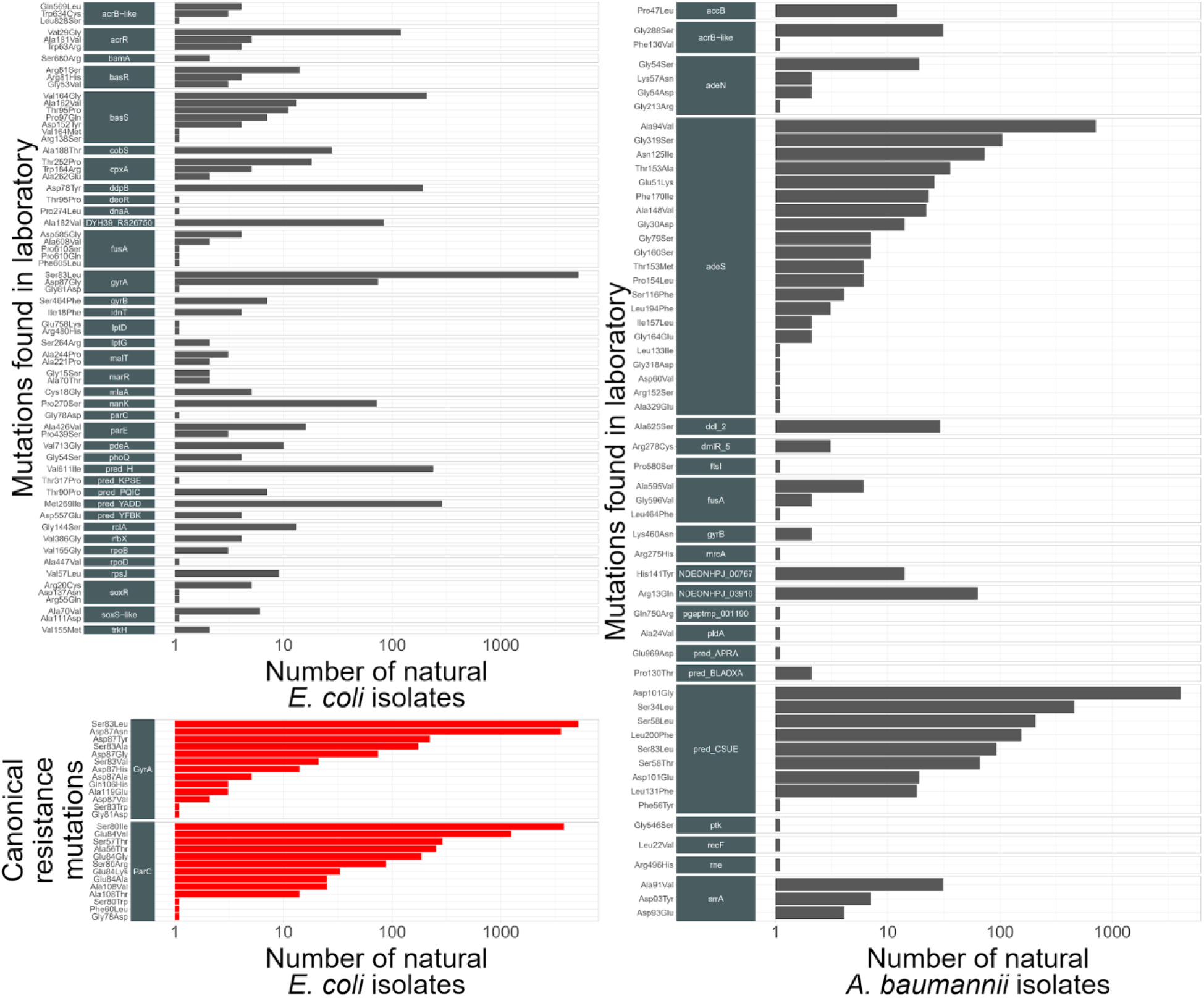
Non-synonymous mutations shared by laboratory evolved lines and natural isolates of *E. coli* and *A. baumannii*. The barplot shows the number of natural isolates with the same non-synonymous mutation that arose during laboratory evolution of resistance. The corresponding genes possessing non-synonymous mutations are labeled with dark gray strips. The bottom left panel shows the number of natural strains with canonical resistance mutations (source: Pointfinder^102^).

**Extended Data Fig. 17.**
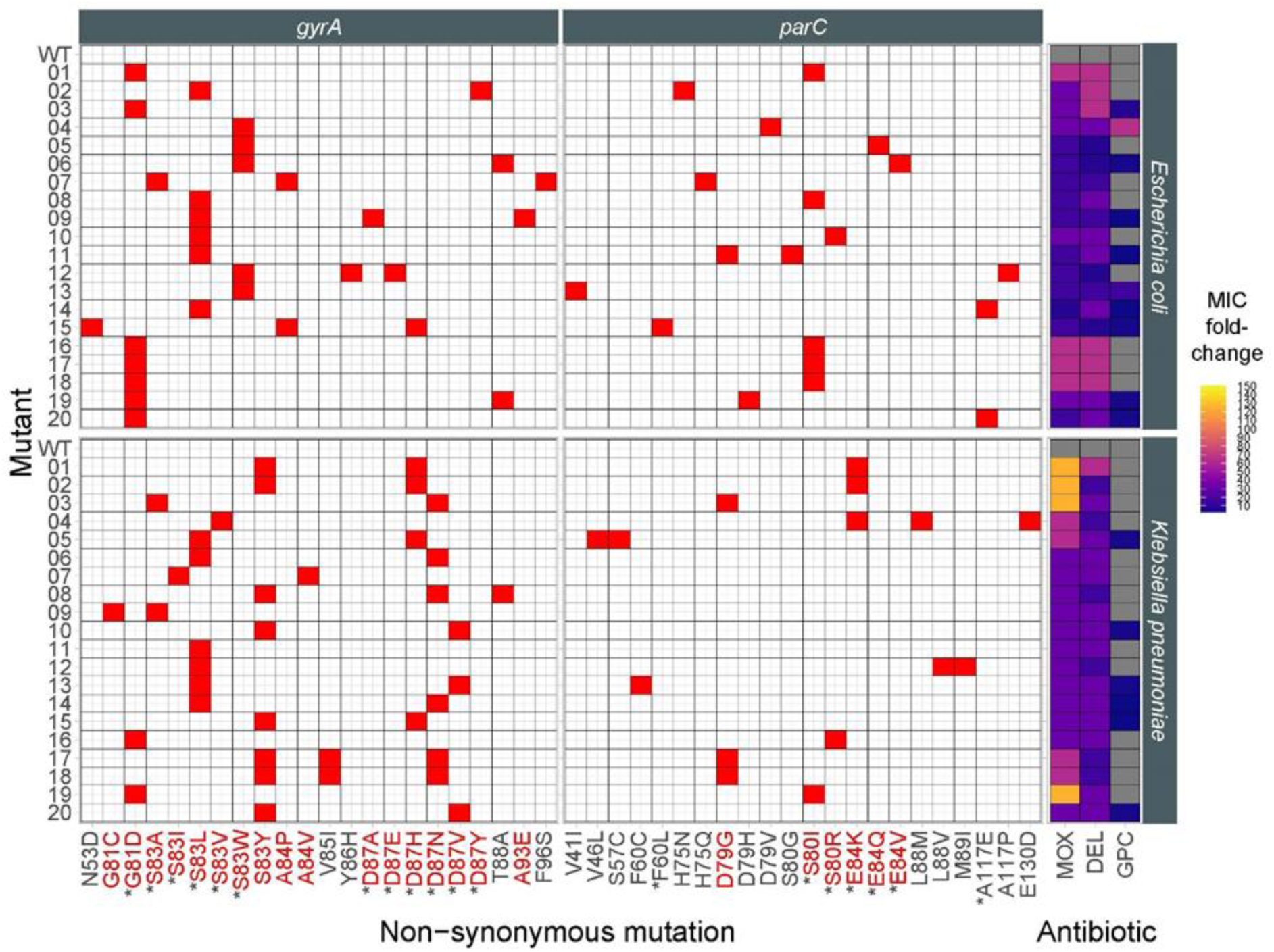
Mutations and the associated cross-resistance patterns across topoisomerase inhibitors. QRDR regions of the *gyrA* and *parC* genes of *Escherichia coli* MG 1655 K-12 and *Klebsiella pneumoniae ATCC* 10031 were subjected to targeted mutagenesis (DIvERGE) and selection for moxifloxacin resistance. The figure shows the mutations carried by the 20-20 strains isolated on moxifloxacin agar plates. The heatmap on the right shows the MIC fold change of the selected mutants to topoisomerase inhibitors, including moxifloxacin (MOX), delafloxacin (DEL) and gepotidacin (GEP). Grey color indicates no change in MIC compared to the wild type. Mutations indicated with red color have been shown previously as topoisomerase resistance-conferring mutations. Asterisks indicate mutations found in the genomes of clinical E*. coli* isolates.

**Extended Data Fig. 18.**
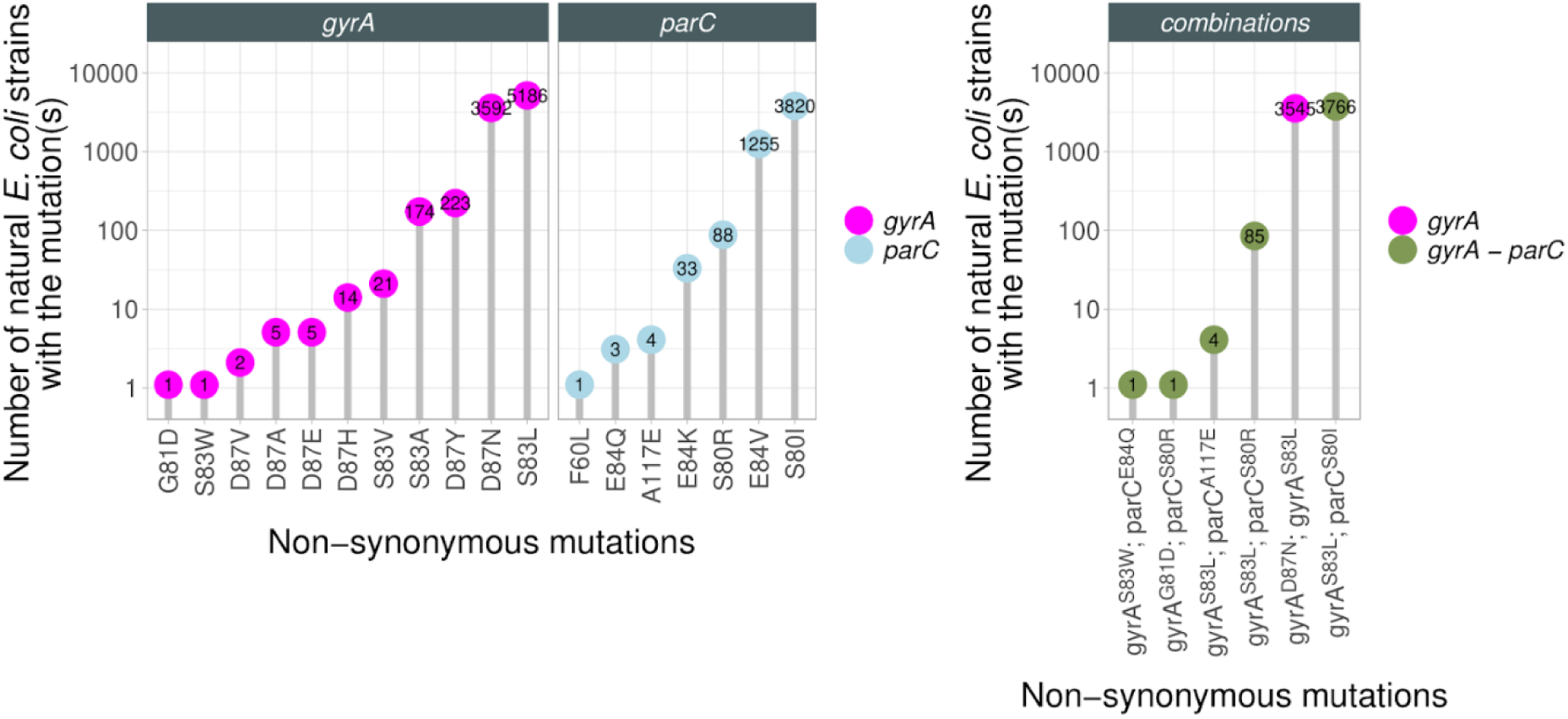
Moxifloxacin resistance mutations in the genomes of natural *E.coli* isolates. The figure shows mutations generated by DIvERGE experiment in *gyrA* (magenta, left panel) and *parC* (blue, middle panel) and their frequencies across the genomes of natural *E. coli* isolates. Combination of mutations (right panel) found in the same resistant clone are also depicted here (magenta: *gyrA-gyrA*, green: *gyrA-parC*). Mutations and combinations thereof in coding sequences were searched for in a dataset of 20,786 natural *E. coli* genomes (see Methods).

**Extended Data Fig. 19.**
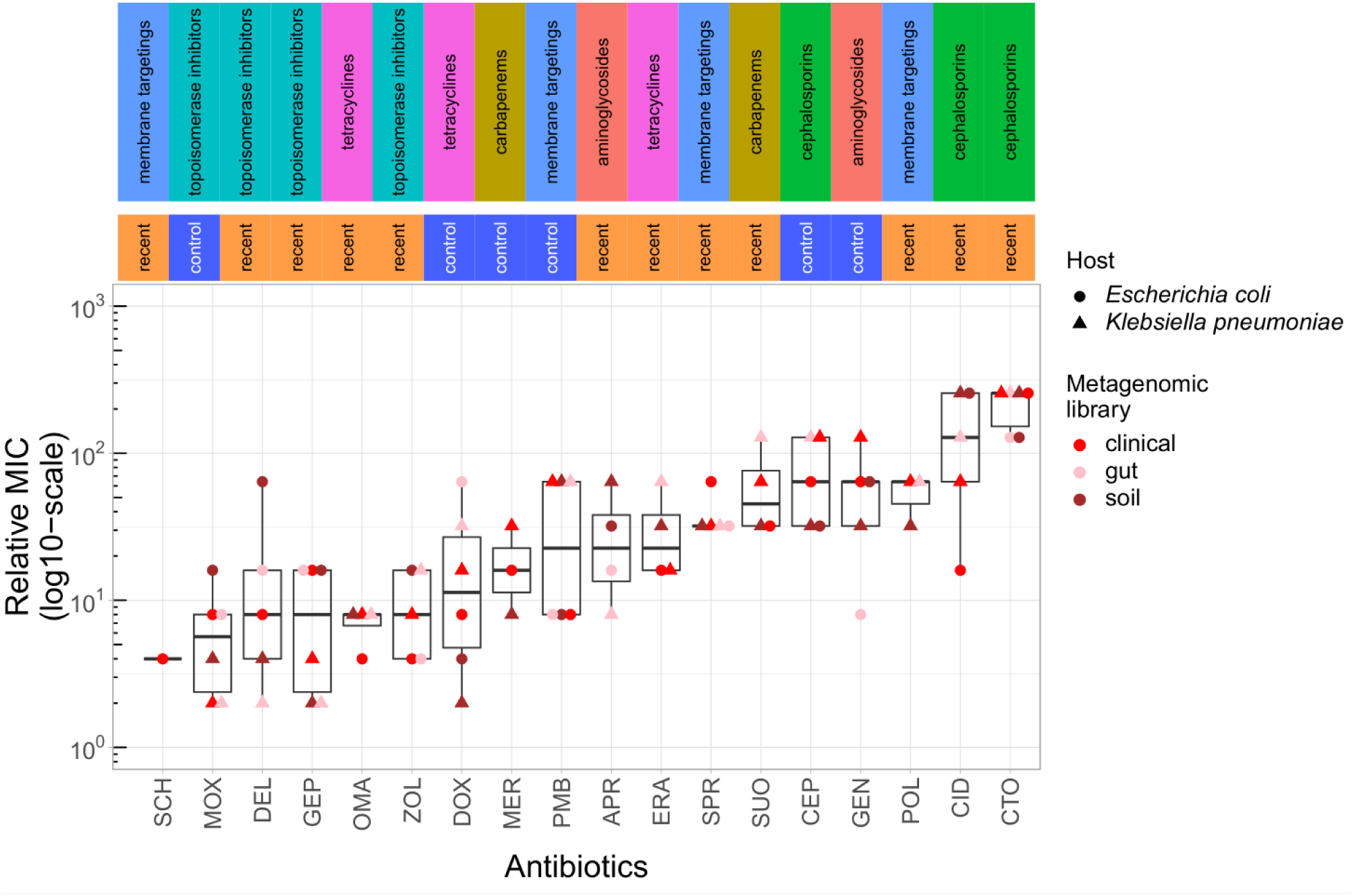
Functional selection of metagenomic libraries with 18 antibiotics resulted in reduced antibiotic susceptibility in the bacterial hosts. The figure shows the MIC fold change in the bacterial host provided by the DNA fragments associated with a given metagenomic library. Only resistant populations or populations with a relative MIC > 1 are depicted here. We observed a substantial, up to 256-fold variation in the resistance level across antibiotics (Kruskal-Wallis chi-squared = 1168.5, df = 18, P < 2.2e-16), whereas no significant difference was observed between the median relative MICs between recent antibiotics and their corresponding within-class controls (paired Wilcox-test, P = 0.7211). Boxplots show the median, first and third quartiles, with whiskers showing the 5th and 95th percentiles. For abbreviations, see Table 1.

**Extended Data Fig. 20.**
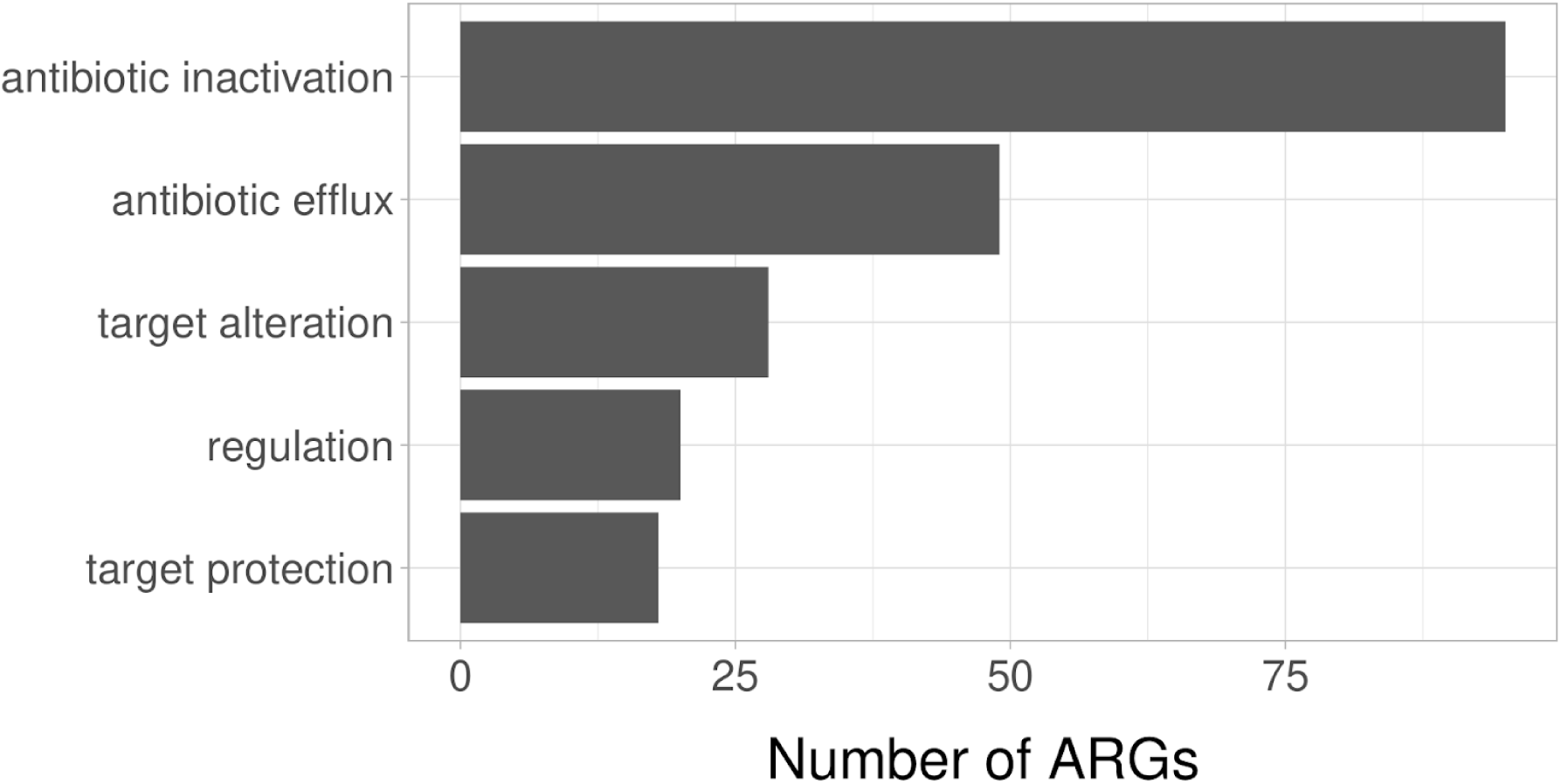
Prevalence of resistance mechanisms for ARGs. The barplot shows the number of ARGs linked to various resistance mechanisms and/or gene regulation.

**Extended Data Fig. 21.**
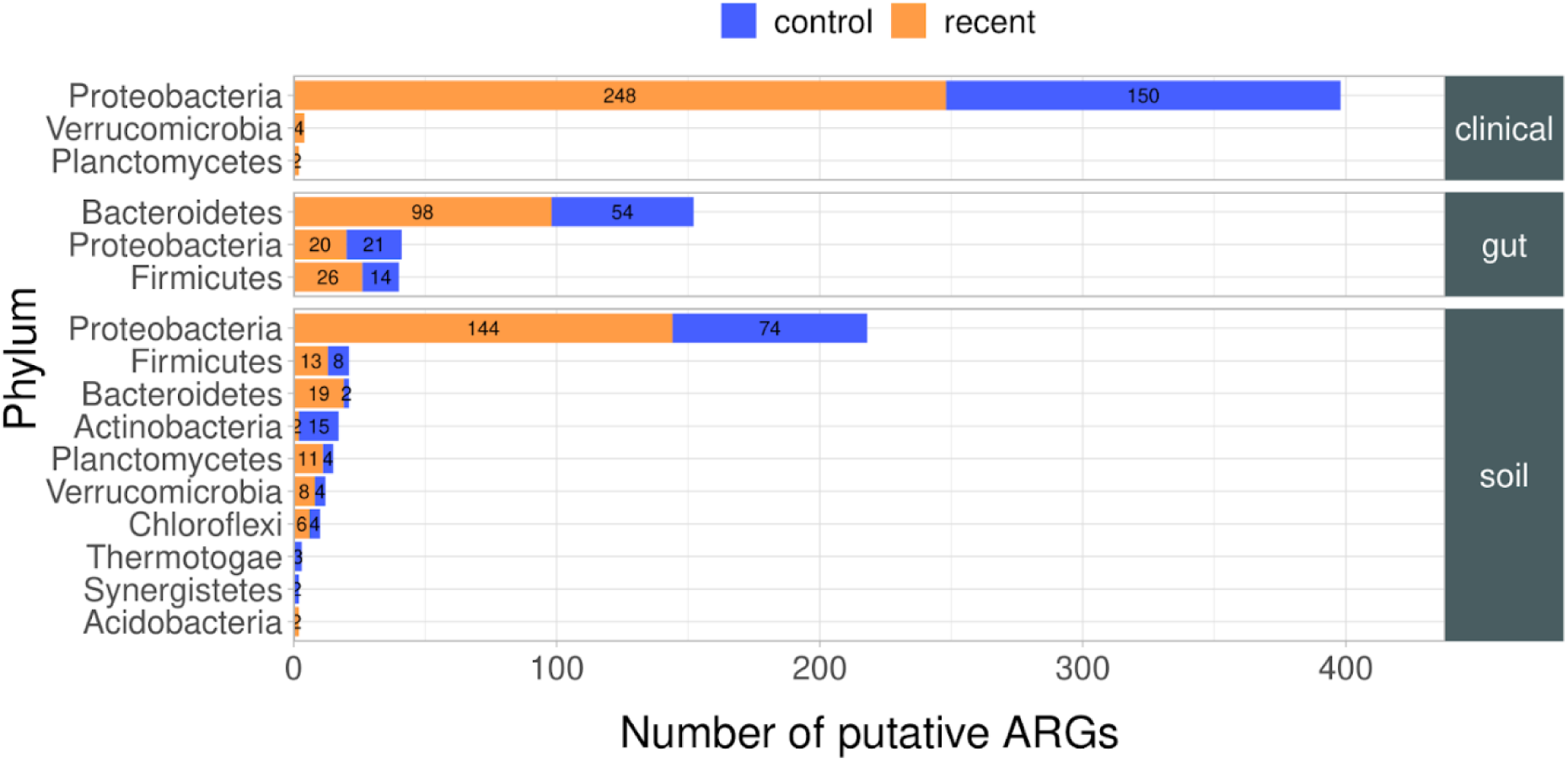
Phylogenetic origin of ARGs. The barplot shows the number of ARGs originating from various phyla. Blue and orange bars denote the number of ARGs detected in control and recent antibiotics, respectively.

**Extended Data Fig. 22.**
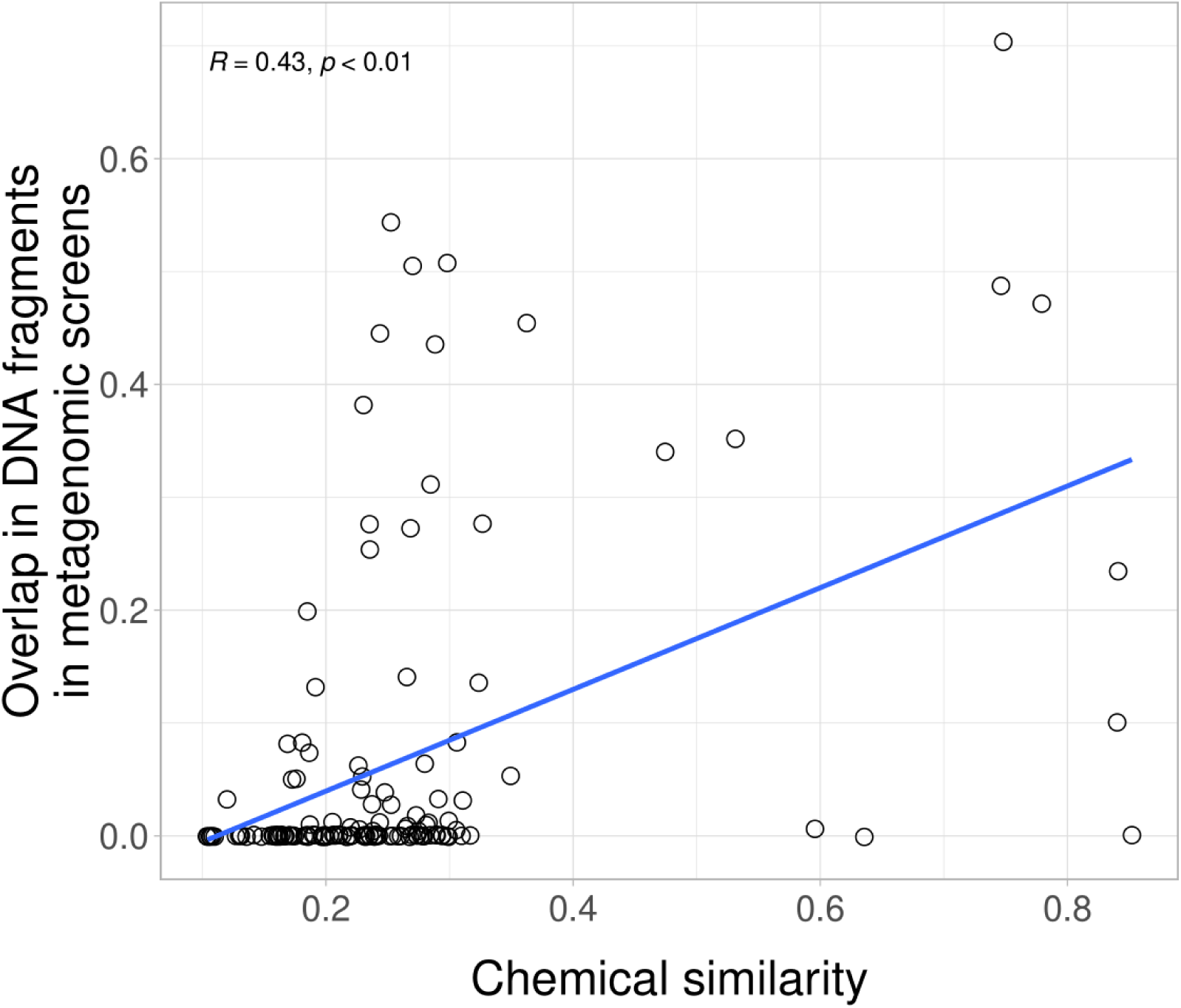
Chemical similarity and co-resistance between antibiotics. The scatterplot shows the association between the chemical similarity of antibiotic pairs and the overlap in resistance-conferring DNA fragments between antibiotic pairs from metagenomic screens. Chemical similarity between antibiotics was estimated by the Tanimoto similarity of their molecular fingerprints (SMILES)^24^. Overlap of resistance-conferring DNA fragments was estimated by the Jaccard similarity, as earlier^100^. There is a significant positive correlation between the two values (Spearman-rank correlation R = 0.43, P < 0.01), even when antibiotic pairs belonging to different antibiotic classes are considered (Spearman-rank correlation R = 0.31, P < 0.01).

**Extended Data Fig. 23.**
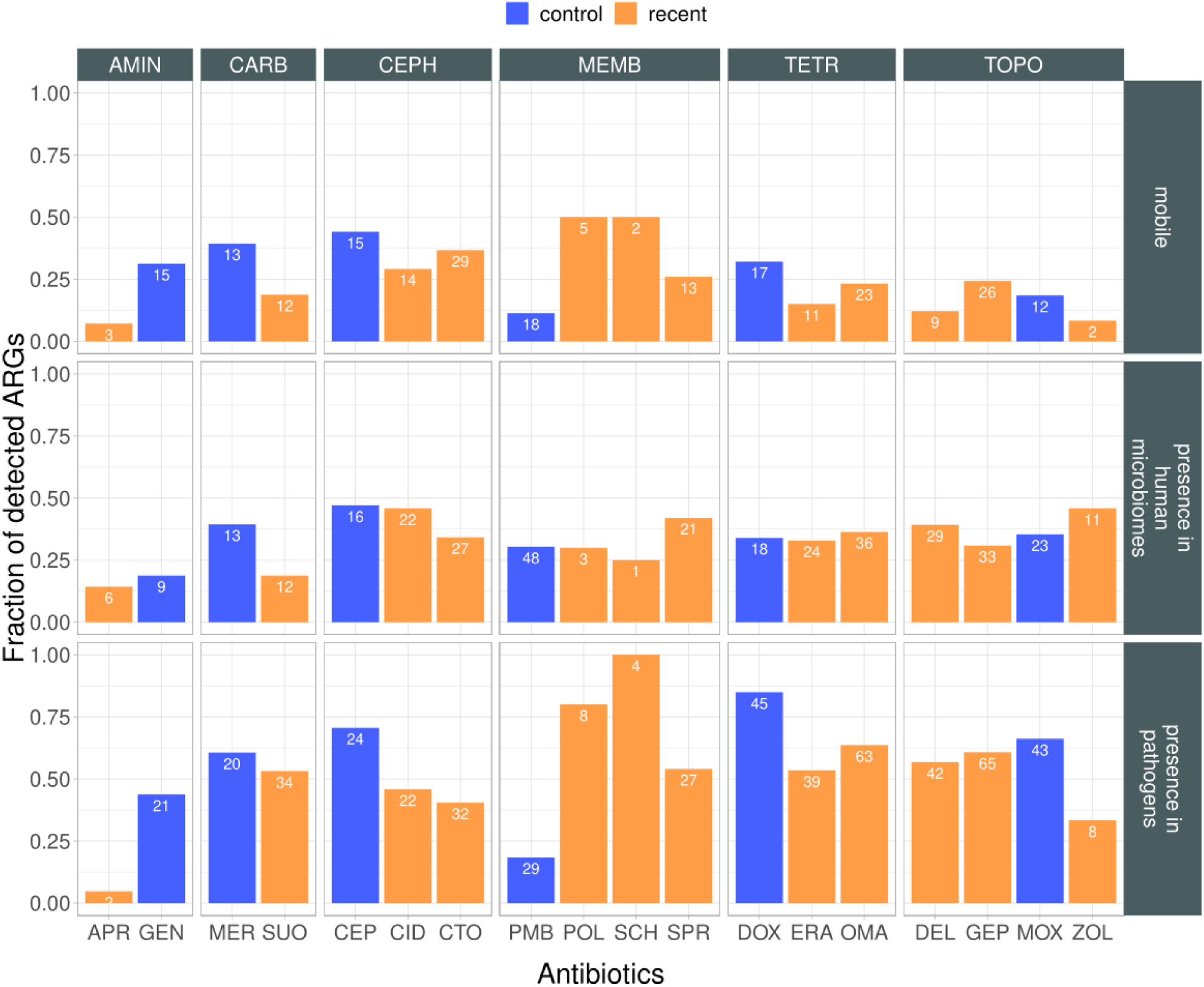
Antibiotic resistance genes and health-risk criteria. The figure shows the fraction of putative ARGs that meet the following criteria: i) gene mobility, ii) presence in microbiomes associated with the human body, and iii) bacterial host pathogenicity. Numbers within the bars indicate the total number of ARGs belonging to the specific category. There is significant variation in the frequency of ORFs across the antibiotics tested (Proportion test, p < 0.05) for each criteria, indicating that adaptations to certain antibiotics are more likely to meet a specific criterium. Abbreviations: TOPO: topoisomerase inhibitors, TETR: tetracyclines, AMIN: aminoglycosides, CARB: carbapenems, CEPH: cephalosporins, MEMB: membrane targeting antibiotics. For antibiotic abbreviations see Table 1.

**Extended Data Fig. 24.**
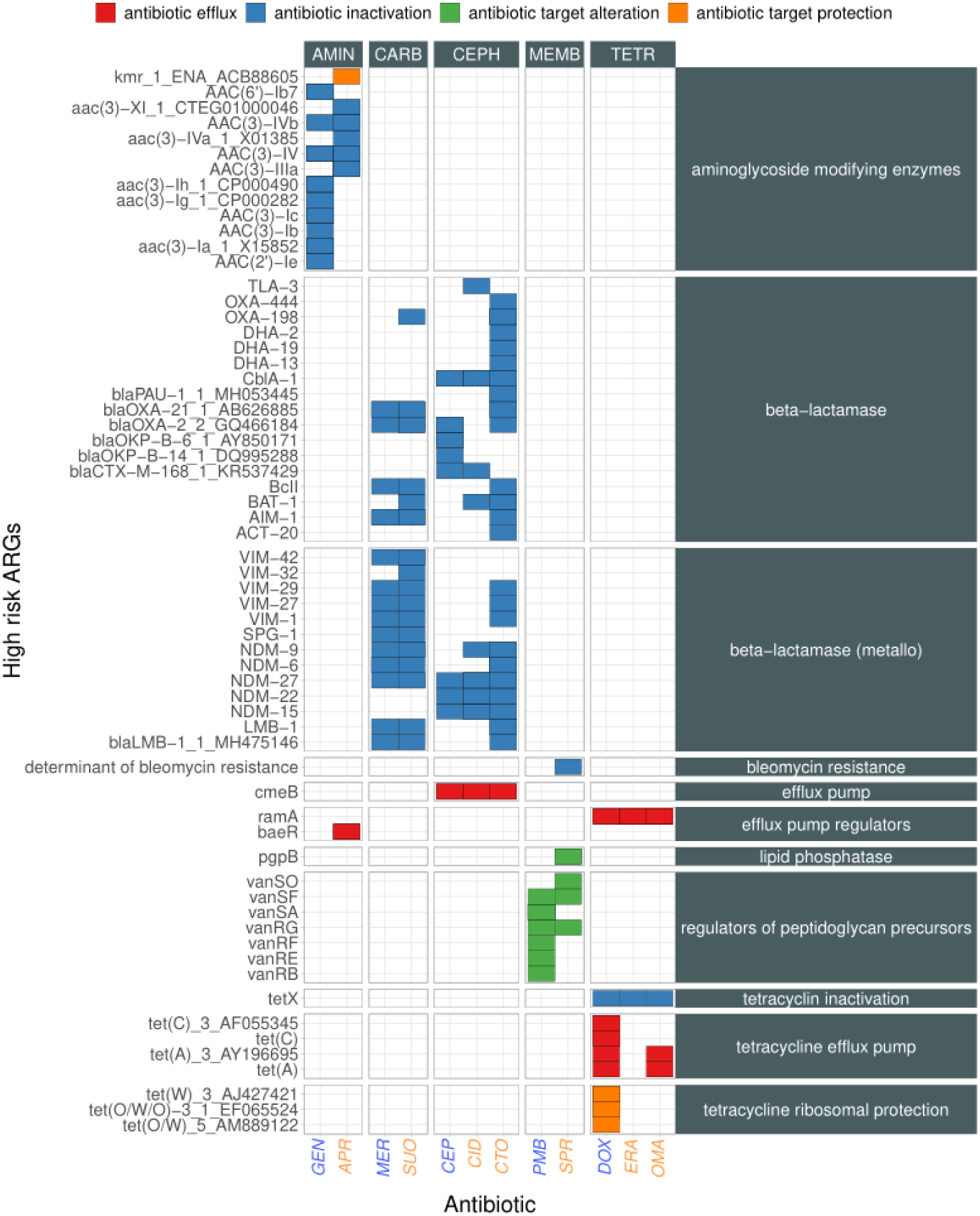
High-risk antibiotic resistance genes against recent antibiotics. The heatmap shows resistance genes identified in functional metagenomics screens. Blue and orange labels indicate control and recent antibiotics, respectively. Only genes that display high sequence similarity to already known antibiotic resistance genes are depicted here. The colors of the heatmap indicate different resistance mechanisms associated with a given antibiotic resistance gene. Horizontal panels depict major types of resistance genes. Antibiotic classes (vertical panels): TOPO (topoisomerase inhibitors), TETR (tetracyclines), AMIN (aminoglycosides), CARB (carbapenems), CEPH (cephalosporins), and MEMB (membrane-targeting antibiotics). For antibiotic abbreviations see Table 1.

**Extended Data Fig. 25.**
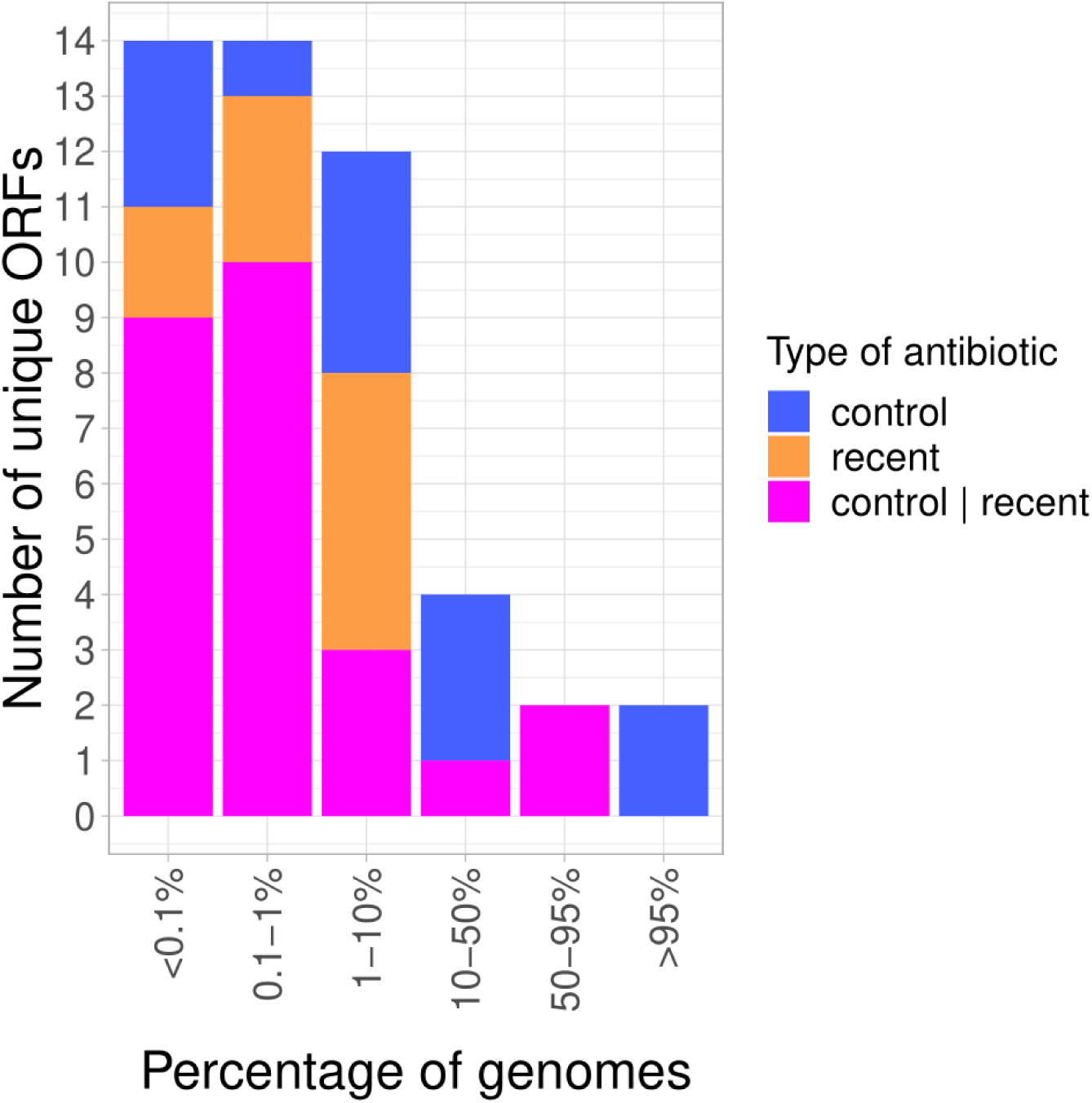
Most putative ARGs are present in only a small fraction of the investigated genomes. The barplot shows the distribution of putative ARGs, detected either against control (blue), recent (orange) or both types of antibiotics (magenta), in the investigated genomes of natural *E. coli* isolates.

**Extended Data Fig. 26.**
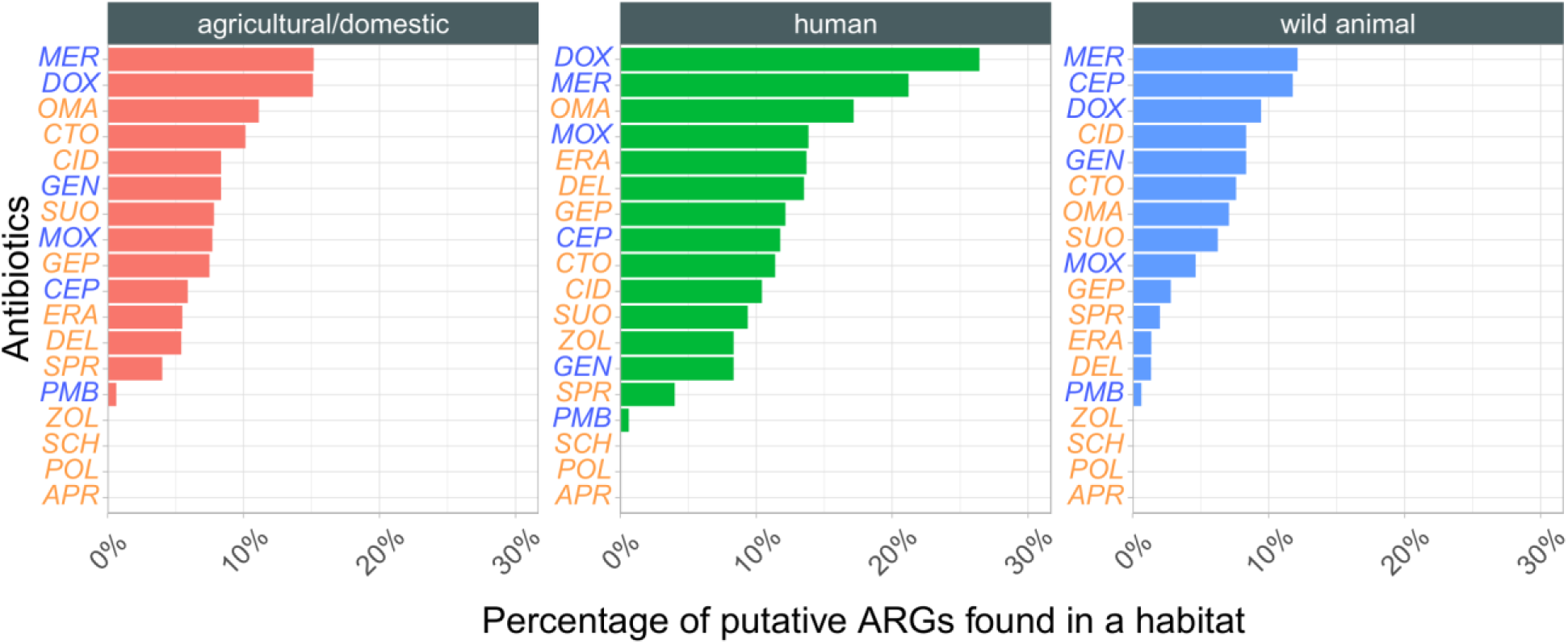
Putative ARGs against various antibiotics are present in natural *E. coli* genomes. The barplot shows the percentage of putative ARGs detected in natural genomes of *E. coli* in a given habitat across control (blue labels) or recent (orange labels) antibiotics. For each habitat, no significant difference was found between the fraction of putative ARGs conferring resistance to control and recent antibiotics (binomial regression model, P = 0.786/0.608/0.302 for agricultural/domestic, human and wild animal, respectively). For abbreviations, see Table 1.

**Extended Data Fig. 27.**
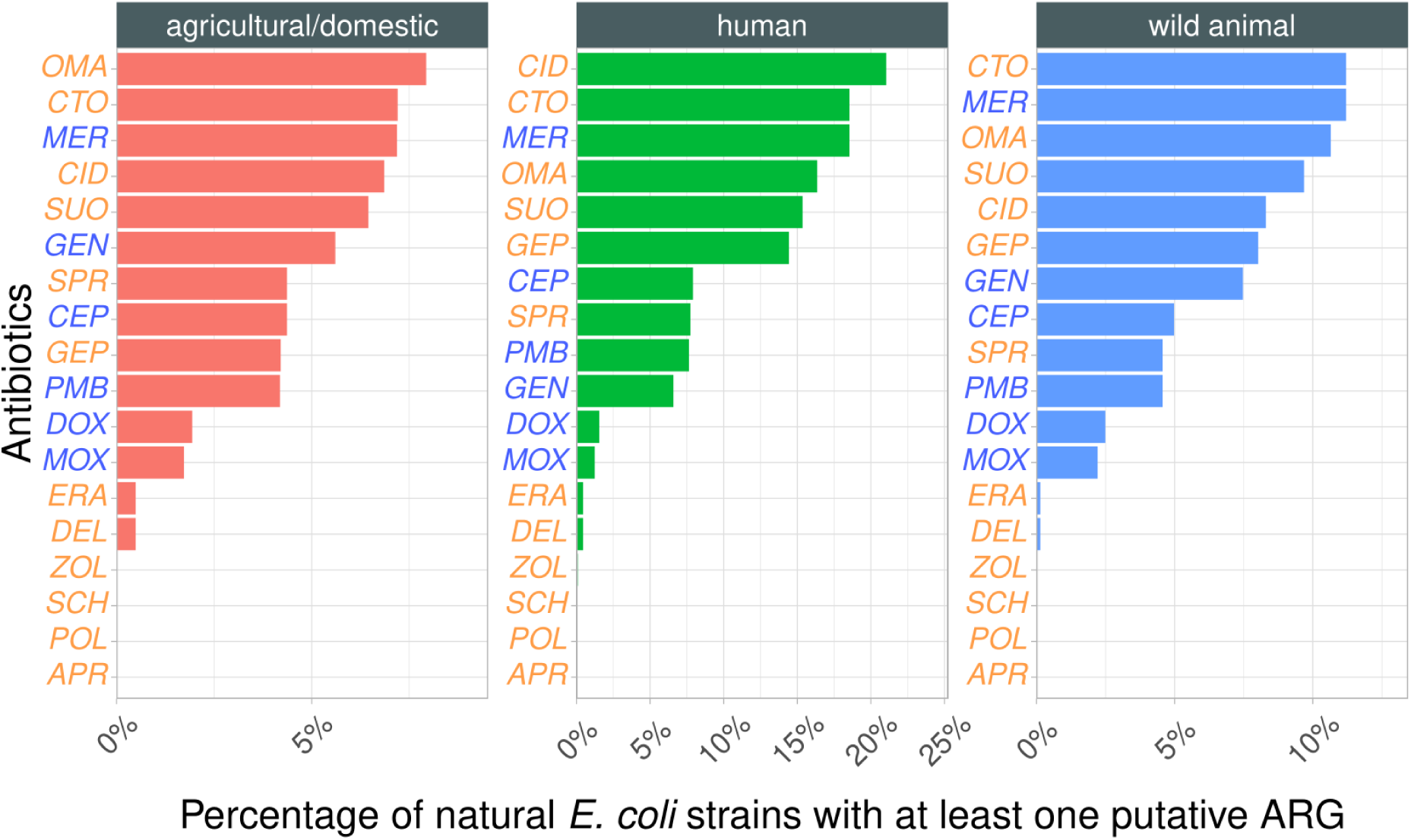
Natural *E. coli* genomes with at least one putative ARG against specific recent antibiotics. The barplot shows the percentage of natural *E. coli* genomes with at least one putative ARGs (present in less than 10% of the investigated genomes) against control (blue label) and recent (red label) antibiotics. For each habitat, we found significant differences between the percentage of natural *E. coli* strains with at least one such putative ARGs that confer resistance to control and recent antibiotics (binomial regression model, P < 0.01). For abbreviations, see Table 1.

**Extended Data Fig. 28.**
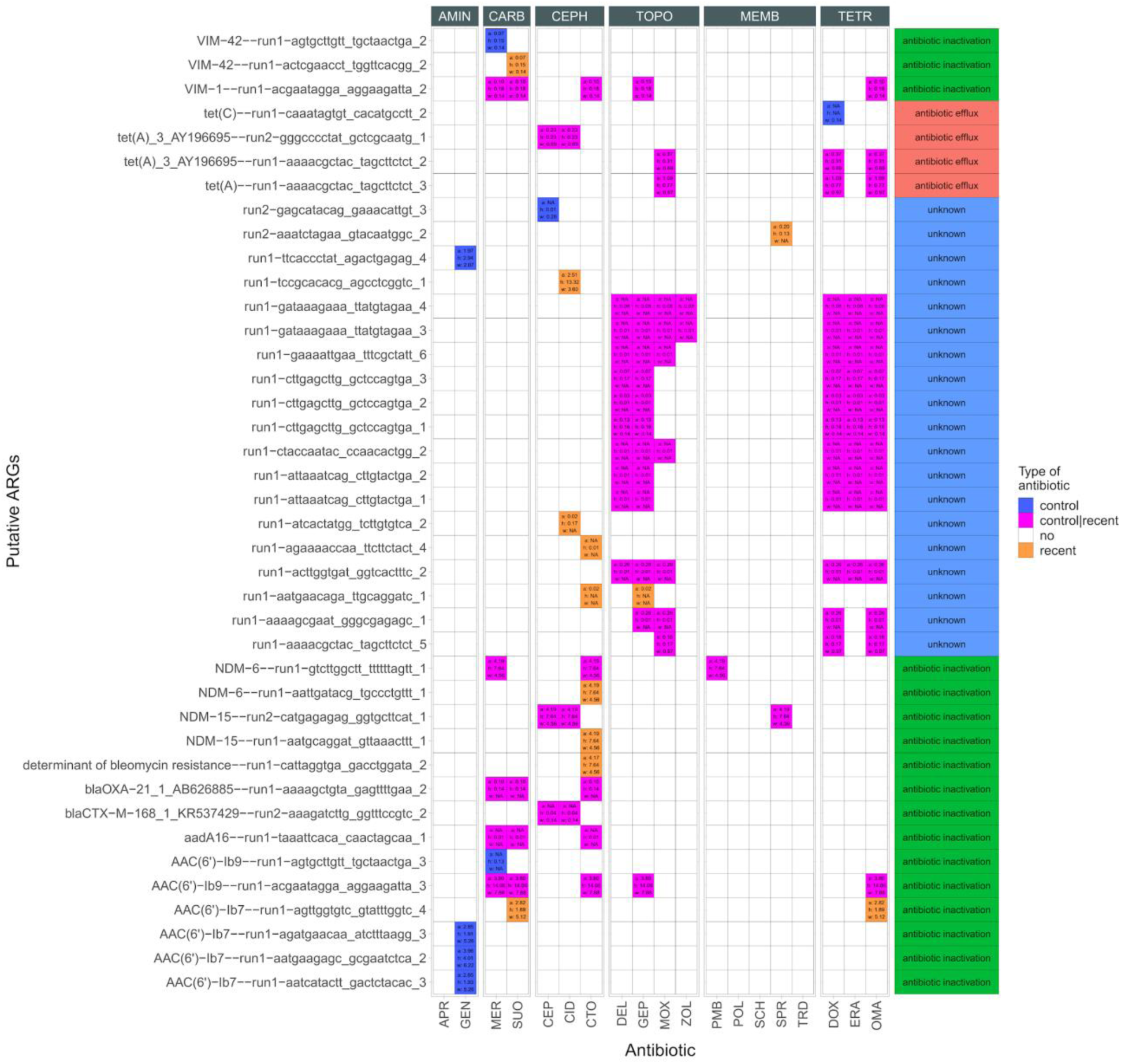
Various putative ARGs could confer resistance to multiple antibiotics. The heatmap shows the distribution of putative ARGs found in natural habitats across antibiotics. Numbers within a cell indicate the number of isolates across habitats (*a*, *h* and *w* denote agriculture, human and wild-animal, respectively). The right panel shows the identified resistance mechanism, if any, for each putative ARGs. ARGs where no sequence similarity was found are labeled with a unique identifier, whereas those that share sequence similarity with known resistance genes (present in either CARD or ResFinder) are labeled with the corresponding gene name.

**Extended Data Fig. 29.**
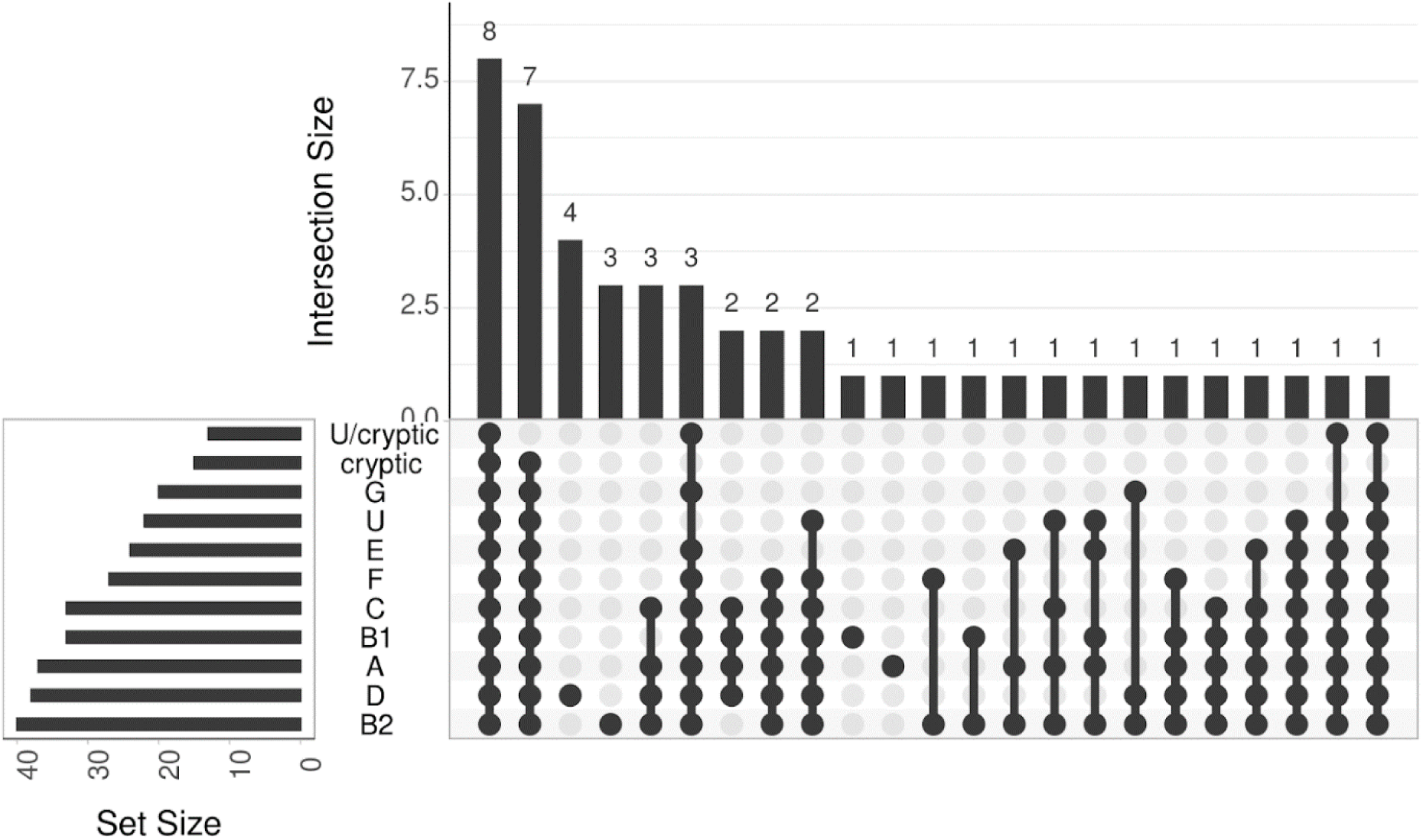
Phylogenetic origin of putative ARGs. The figure shows the overlap of putative ARGs across different phylogroups of *E. coli*. The plot shows 48 putative ARGs detected in at least one *E. coli* genome. Letters on the left (A, B1, B2, D, E, F, G, U, U/cryptic, and cryptic) denote phylogroups, according to a previous study^26^.

**Extended Data Fig. 30.**
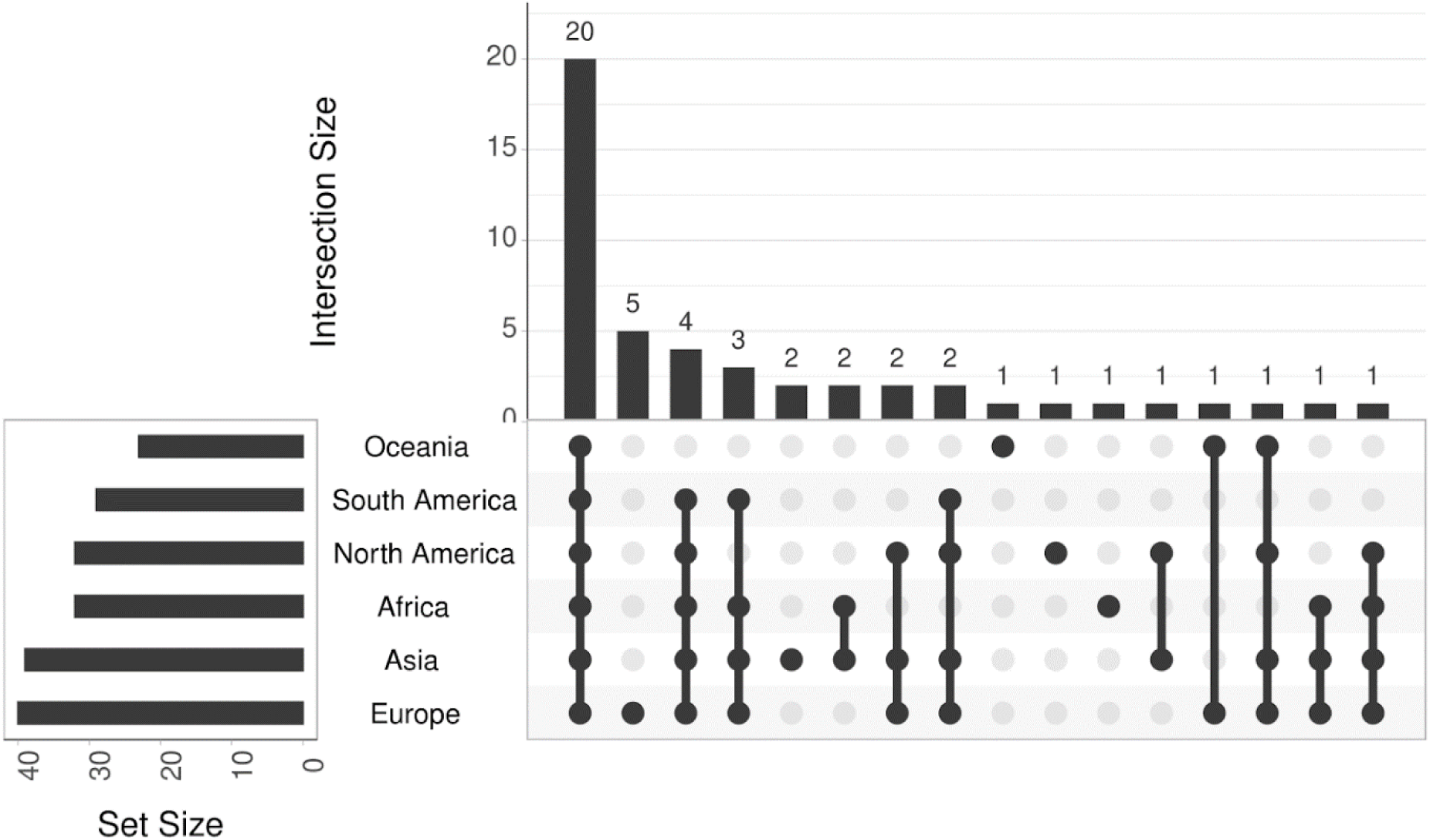
Geographical origin of putative ARGs. The figure shows the overlap of putative ARGs across *E. coli* from different continents.

**Extended Data Fig. 31.**
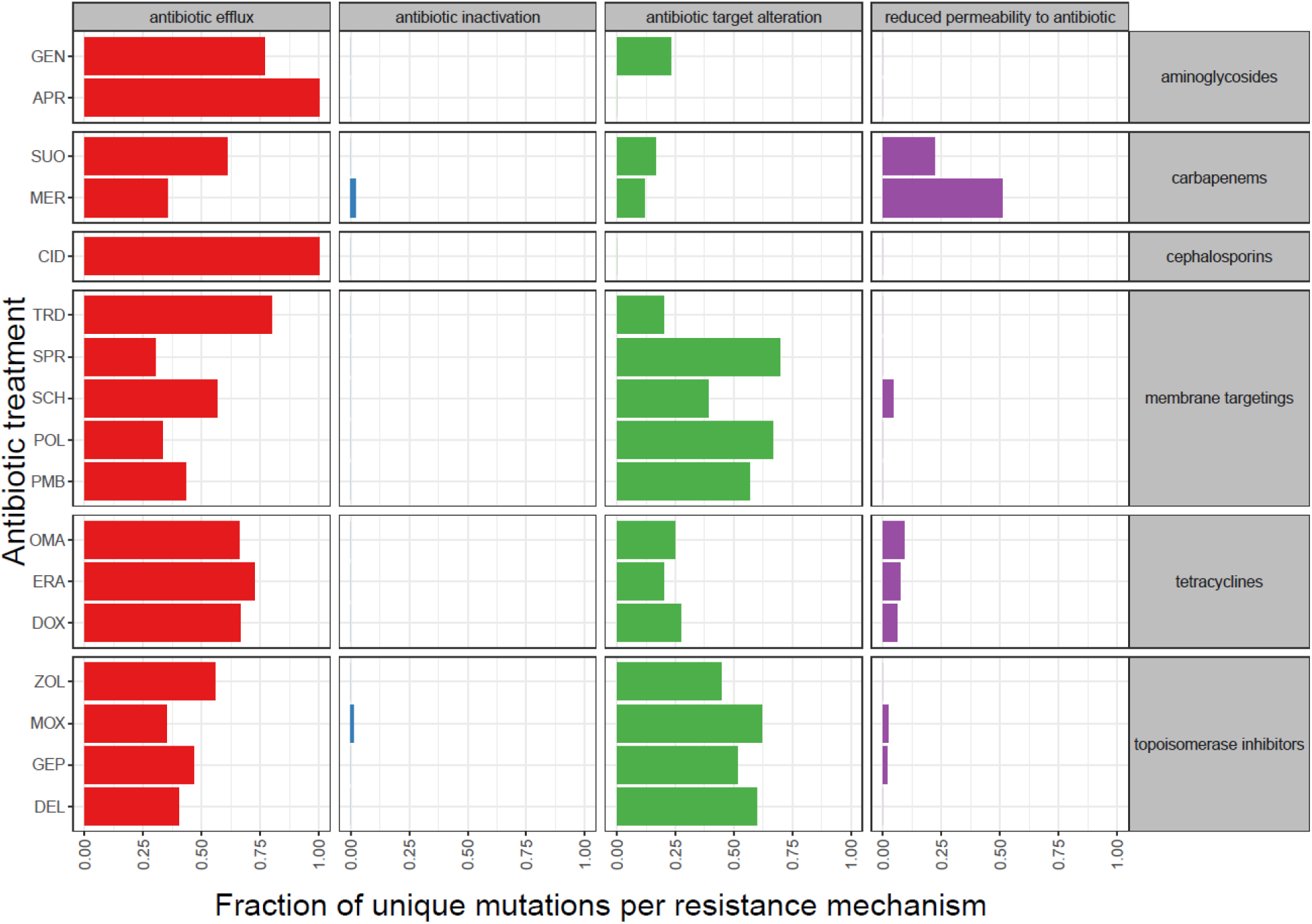
Distribution of canonical resistance mechanisms across antibiotic treatments. Unique de novo genomic mutations were assigned to four major resistance mechanisms based on homology to genes featured in the CARD and ResFinder databases. Those antibiotic treatments were disregarded that did not yield at least 10 unique mutations during the course of laboratory evolution. The four major resistance categories include reduced membrane permeability, antibiotic target alteration, antibiotic inactivation, and antibiotic efflux. The distribution of mutations belonging to each resistance mechanism differs across the antibiotics (Proportion test, p < 0.001). For antibiotic abbreviations see Table 1.

**Extended Data Fig. 32.**
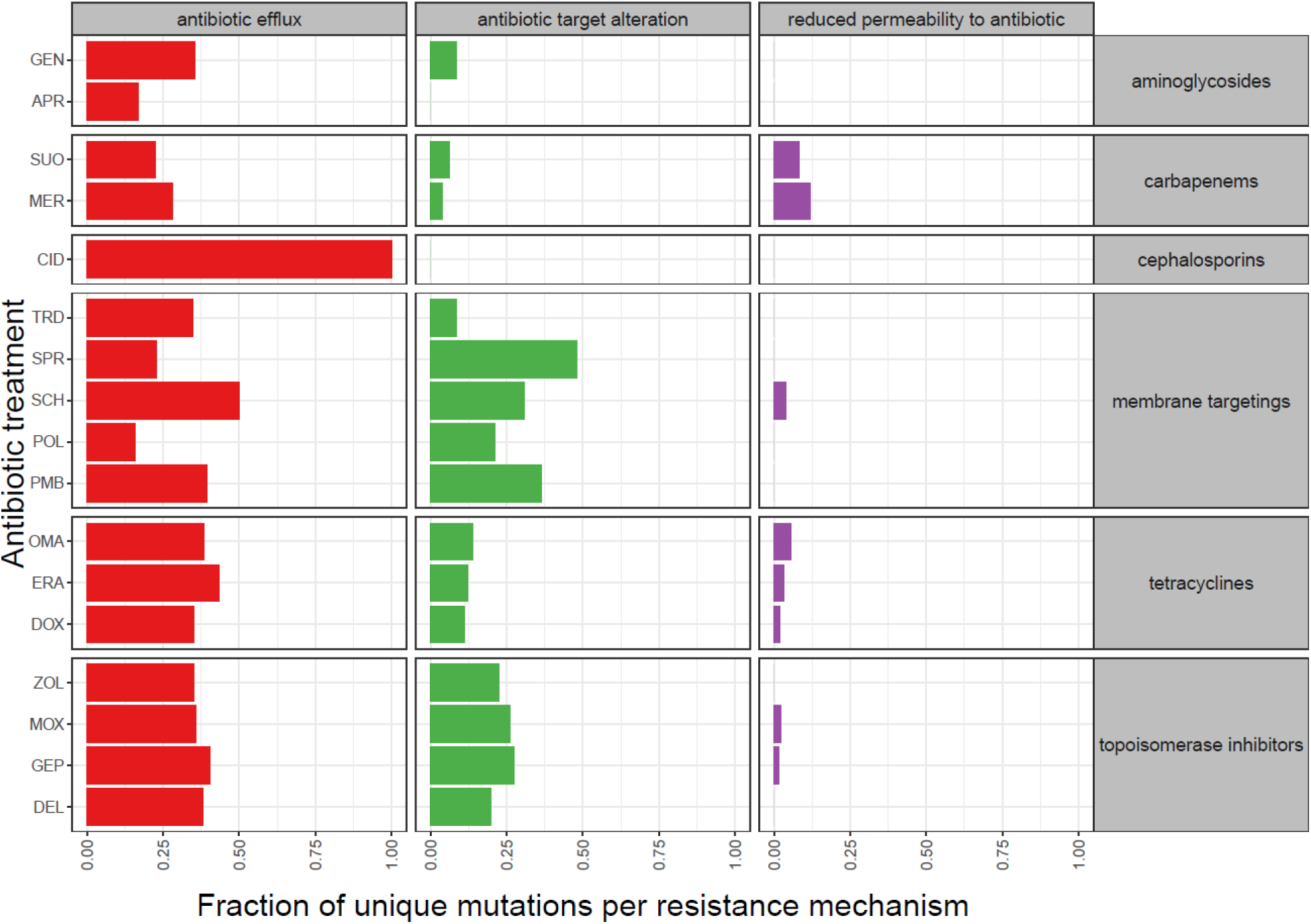
The distribution of canonical resistance mechanisms among putative multi-drug resistance genes shows strong heterogeneity across antibiotic treatments. Unique mutations were assigned to three major resistance mechanisms based on homology to genes featured in the CARD and ResFinder databases. Those antibiotic treatments were disregarded that did not yield at least 10 unique mutations. The three major resistance categories include reduced membrane permeability, antibiotic target alteration, and antibiotic efflux. The distribution of mutations differs in all three functional categories across the antibiotics tested (proportion test, p < 0.05), indicating that adaptations to certain antibiotics favor certain canonical resistance mechanisms over others. The figure depicts the distribution of canonical resistance mechanisms of putative multidrug resistance genes, i.e those mutated in at least two independent lines per genetic background and in adaptation against at least two antibiotic classes. For antibiotic abbreviations see Table 1.

**Extended Data Fig. 33.**
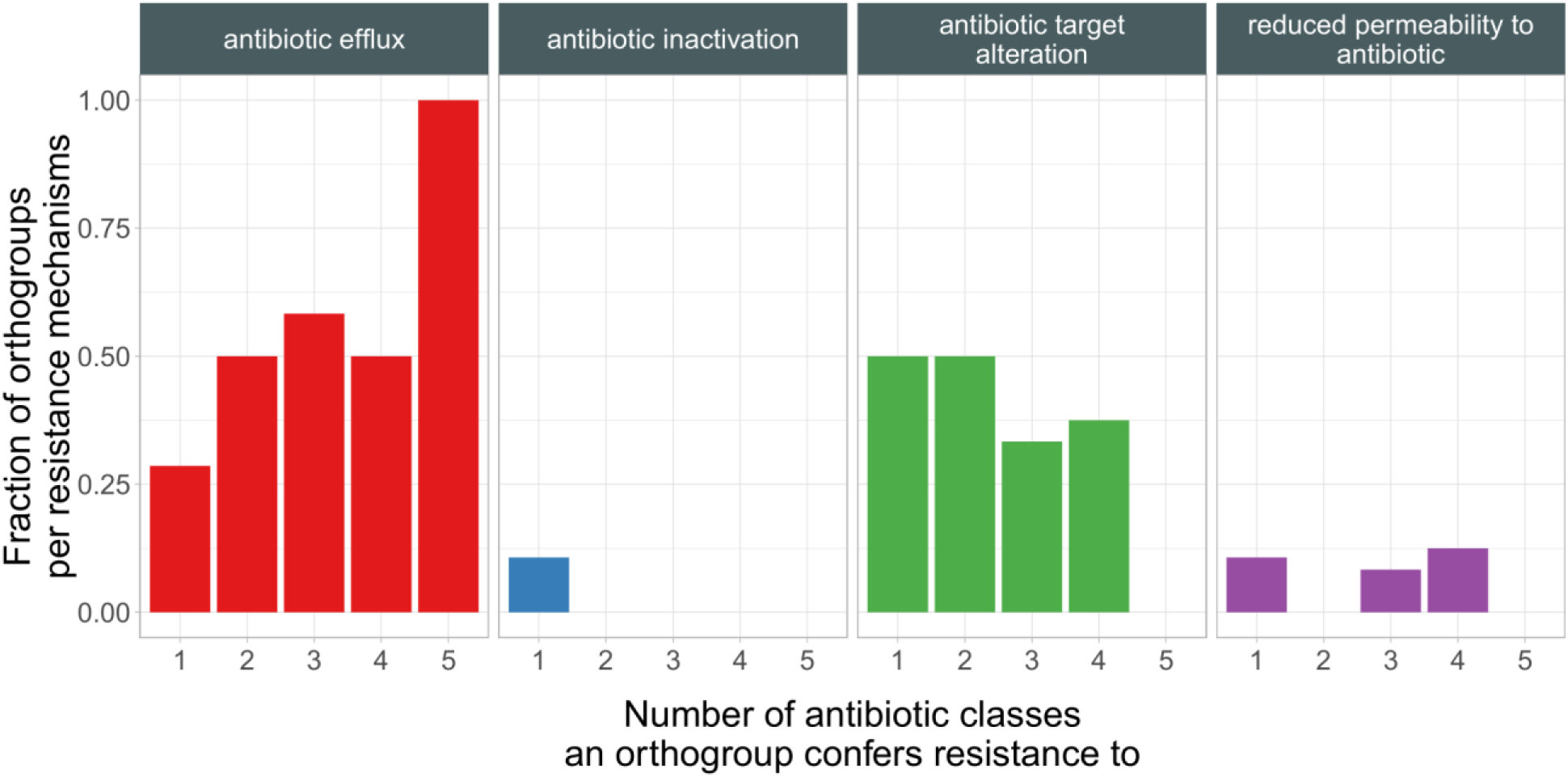
Genes with mutations conferring resistance to multiple antibiotic classes are preferentially involved in antibiotic efflux. The barplot shows the fraction of mutations related to various resistance mechanisms (panels) as a function of the number of antibiotic classes the mutations confer resistance to. Colors indicate different resistance mechanisms.

**Extended Data Fig. 34.**
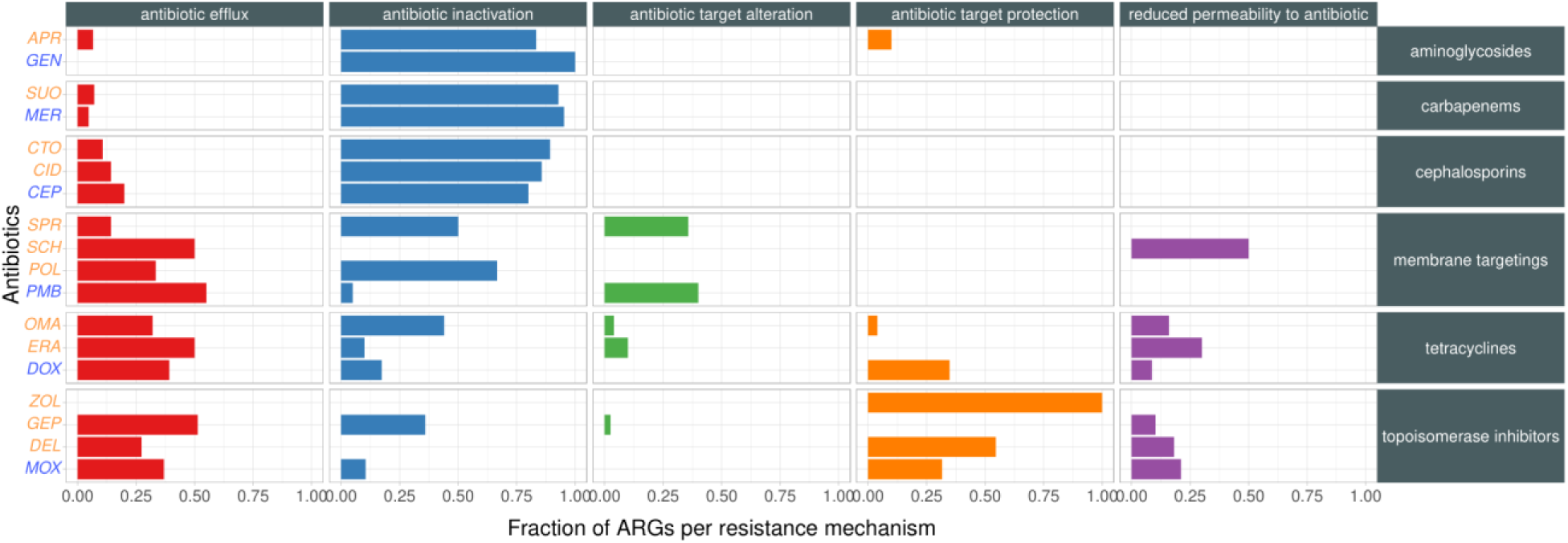
Distribution of resistance mechanisms across functional metagenomics screens. The barplot shows the fraction of ARGs per antibiotic resistance mechanisms across functional metagenomics screens in the presence of different antibiotics (blue and orange indicate control and recent antibiotics, respectively). ARGs were assigned to five major resistance mechanisms (vertical panels) based on homology to genes featured in the CARD and ResFinder databases. The five major resistance categories include antibiotic efflux. antibiotic inactivation, antibiotic target alteration, antibiotic target protection, and reduced permeability to antibiotics. The distribution of ARGs belonging to each resistance mechanism shows heterogeneity across the antibiotics (Proportion test, p < 0.001).

**Extended Data Fig. 35.**
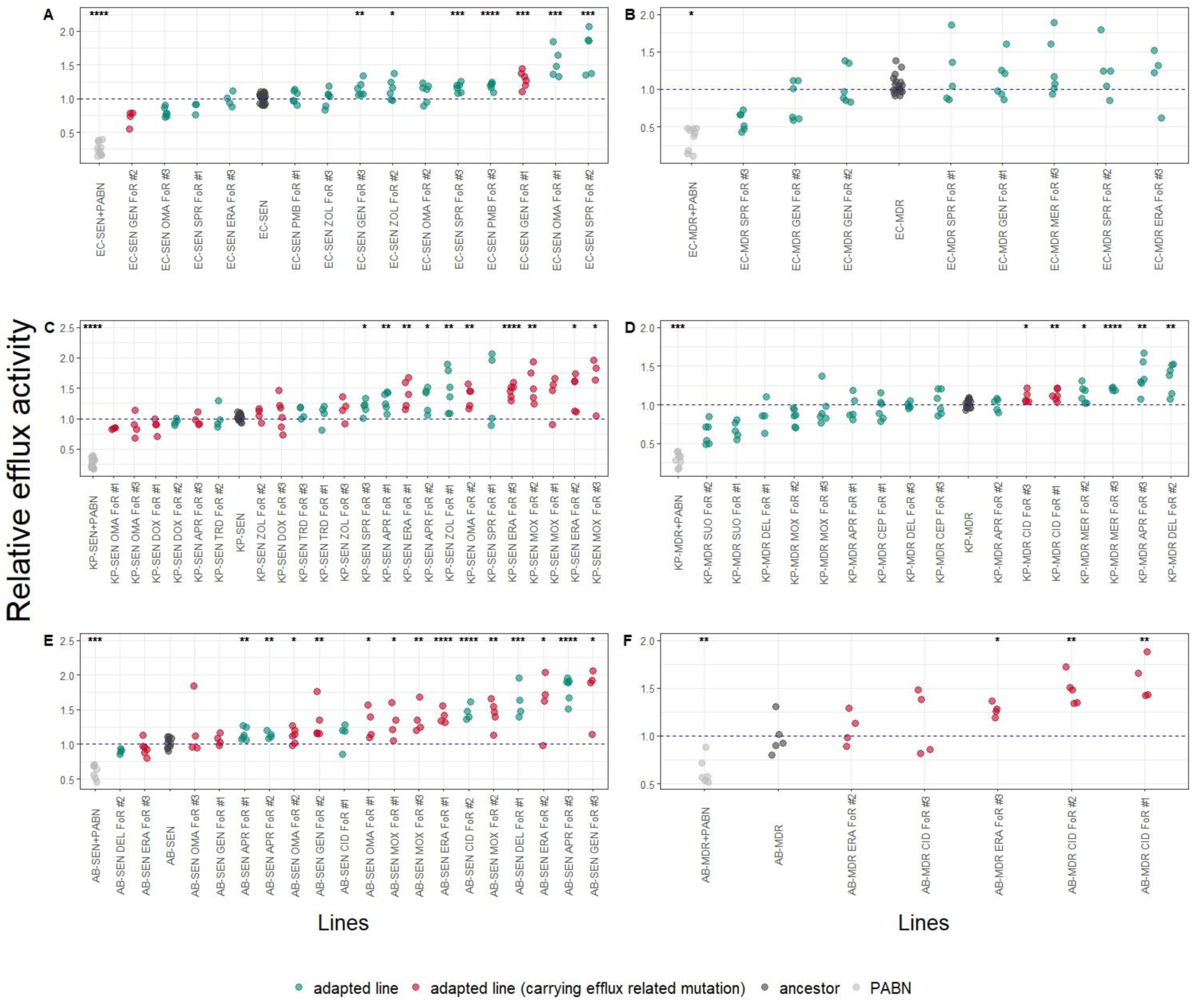
Efflux activity in antibiotic-adapted lines. The figure shows the efflux activity of the evolved lines from FoR assay in *E. coli* (A-B), *K.pneumoniae* (C-D), and *A.baumannii* (E-F). We measured the intracellular accumulation of a fluorescent probe (Hoechst 33342), where lower dye accumulation indicates higher efflux activity/reduced membrane permeability. We quantified the relative change in optical density-normalized fluorescence signal during a fixed timeframe (see Methods). Efflux activity was determined by normalizing this change measured in the tested strains against that in their respective ancestral strain and taking the inverse of its value. Lines displaying significantly reduced Hoechst-dye accumulation, i.e. higher efflux activity, compared to the ancestor strains are indicated by asterisks (one-tailed Student’s t- test: ****/***/**/* indicates p <= 0.0001/0.001/0.01/0.05, respectively). The dashed line indicates the average of the corresponding ancestor. Ancestor strains treated with an efflux pump inhibitor (Phenylalanine-arginine β-naphthylamide (PAβN)) served as a positive control, and displayed significantly larger Hoechst-dye accumulation value, compared to that of the non-treated ancestor strains.

**Extended Data Fig. 36.**
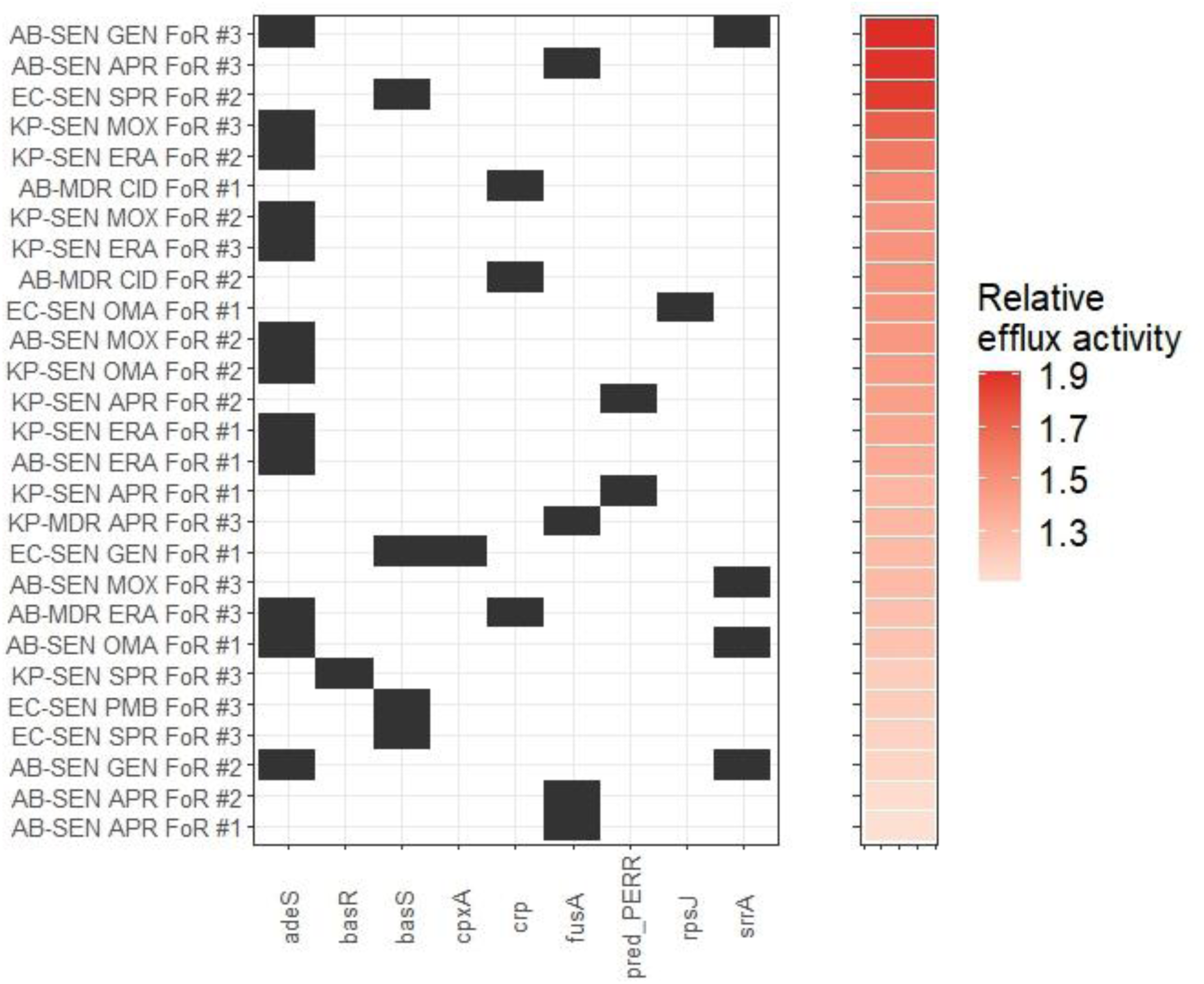
Mutated genes in lines with enhanced efflux pump activity. The heatmap represents the mutations of lines with increased efflux activity. The right panel shows the relative efflux activity.

**Extended Data Fig. 37.**
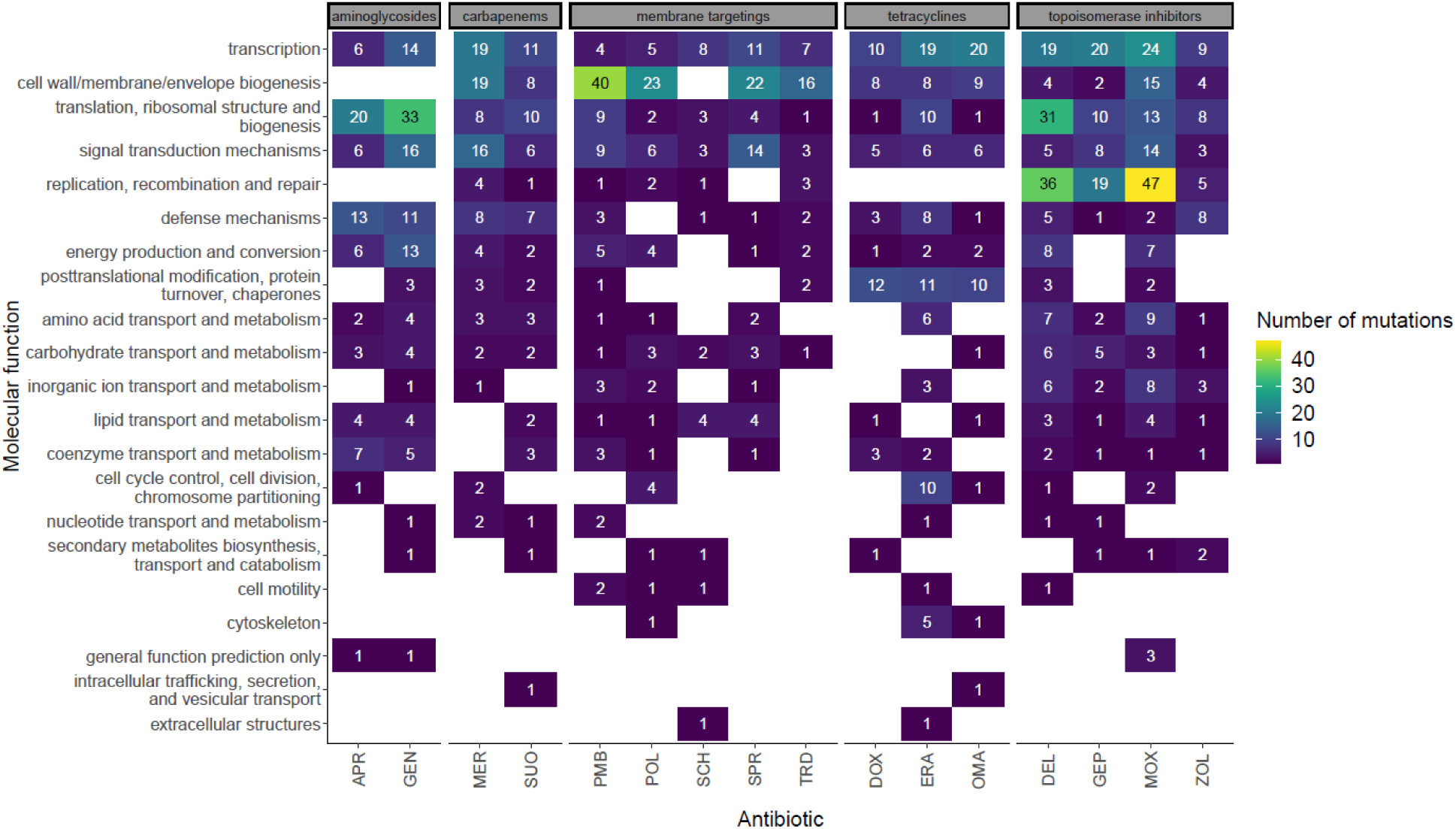
Molecular functions of the genes mutated in response to different antibiotics. Unique mutated genes were assigned to functional categories based on information obtained from the Clusters of Orthologous Groups of Proteins database (COG). The figure shows the total number of unique mutations associated with each molecular function category per antibiotic treatment.

