## Supplementary Notes and Figures for "Antibiotics of the future are prone to resistance in Gram-negative pathogens"

**The file includes:**

Supplementary Notes 1-4

**Other Supplementary Information for this manuscript includes the following:**

Supplementary Tables 1-12
Supplementary Data 1

**Supplementary Note 1.**

Recently approved antibiotics

Four recently approved antibiotics (cefiderocol, eravacycline, omadacycline and delafloxacin) were included, the reasons are as follows.

Since 2017, thirteen antibiotics have gained market authorization from European countries or the FDA[1]. Out of the 13 approved new antibiotics, **cefiderocol (CID)** (a siderophore cephalosporin) was specifically designed against carbapenem-resistant *P. aeruginosa* and *A. baumannii*, while five others act against bacterial strains belonging to the *Enterobacteriaceae* family.

These five antibiotics include **eravacycline (ERA)**[2], a new glycylcycline derivative, **ceftobiprole (CTO)**[3], a 5th generation cephalosporin, vaborbactam and relebactam, both β-lactamase inhibitors and plazomicin, a new aminoglycoside derivative. As mentioned earlier, the evolution of resistance to β-lactamase inhibitors will be studied elsewhere. Plazomicin was also excluded due to unavailability.

**Omadacycline (OMA)**[4]**,** an aminomethylcycline, and **delafloxacin (DEL)**[5]**,** a 4th generation fluoroquinolone were selected, as they inhibit the growths of *K. pneumoniae* and *P. aeruginosa* in preclinical studies.

The remaining antibiotics mainly target Gram-positive bacteria (e.g. lascufloxacin, levanadofloxacin, contezolid and lefamulin) or Mycobacteria (e.g. pretonamid), and therefore they were omitted from the present study.

Antibiotics in clinical development

According to the 2021 WHO report on the clinical developmental pipeline[1], there are 27 antibiotics in Phase 1-3 clinical trials. More than half of these (n = 14) were previously shown to be effective gram-negative ESKAPE pathogens. However, out of these 14 antibiotic candidates, 9 are β-lactam – BLI combinations that are beyond the scope of this study.

From the remaining 5 potential antibiotic candidates, we selected **SPR-206 (SPR)**[6], **apramycin (APR)**[7] (an aminoglycoside already used in veterinary practice), and **sulopenem (SUO)**[8] (carbapenem derivative).

We also study two topoisomerase inhibitors currently in clinical phase 3 development[9]: **zoliflodacin (ZOL)**[10] and **gepotidacin (GEP)**[11]. They are active against resistant Gram-negative pathogens (ZOL – *N. gonorrhoeae, M. catarrhalis*; GPC – *E. coli*), and both have novel modes of action, compared to other established topoisomerase inhibitors, such as fluoroquinolones.

Antibiotics in preclinical development

The preclinical antibiotic development pipeline in 2021 consisted of more than 200 compounds. We focused on compounds that specifically target specific components of the bacterial membrane or the cell envelope, for two reasons. First, such compounds have been previously suggested to be relatively immune to bacterial resistance, and second, as high as 25% of the antibiotic candidates in preclinical development belong to this category[1].

Furthermore, in an influential paper, Kim Lewis has described 7 promising novel compounds from the preclinical pipeline with efficacy in the mouse infection model. 5 out of the 7 are effective against Gram-negatives, and four target the bacterial cell membrane[12]. From this short list, we selected **POL7306 (POL)**[13], a polymyxin/murepavadin chimera, and **Tridecaptin M152-P3 (TRD)**[14]. POL7306 targets BamA, an outer membrane protein of Gram-negative cells. TRD targets lipid II, a precursor of peptidoglycan synthesis.

**SCH79797 (SCH)**[15] is a repurposed drug, as it is a selective antagonist of the Thrombin Receptor Proteinase Activated Receptor 1. This compound was recently shown to have promising antibacterial effects. SCH exhibits membrane disintegrating and folate synthesis inhibitor effects simultaneously, thus it was claimed to be a multi-targeting, resistance-free antibiotic.

**Supplementary Note 2.**

**<u>Elevated genomic mutation rate does not promote SCH79797 resistance in *A. baumannii*</u>**

SCH-79797, a dual-targeting antibiotic, effectively kills a wide range of bacterial pathogens without detectable resistance[15], but the underlying data was limited. By studying this issue in three Gram-negative bacterial pathogens, we found high variability in resistance evolution across bacterial species. In particular, we found as high as a 32-fold increment in SCH-79797 resistance level in *K.pneumoniae* (KP-SEN), while only an average 1.8-fold change was observed in *A. baumannii* (AB-SEN). One might argue that the evolution of SCH-79797 resistance might be limited by the mutational supply rate in our experimental system. Indeed, bacterial lines with an elevated mutation rate (mutators) are known to display an exceptionally rapid evolution of antibiotic resistance[16]. Therefore, we have tested whether higher resistance against SCH-79797 emerges in such mutator populations. To investigate this issue, we focused on an isogenic hypermutator *A. baumannii* strain isolated from an earlier SCH-79797 resistance screen. As a result of a mutation (Leu544*) in the mismatch-repair gene *mutS*, the strain exhibits an over 250-fold increment in genomic mutation rate (SN-Fig. 1.). The corresponding mutator populations were exposed to toxic SCH-79797 antibiotic exposure. As earlier, we used a standard protocol for spontaneous frequency-of-resistance analysis (FoR assay) at multiple SCH-79797 concentrations of each antibiotic. Approximately $10^{10}$ bacterial cells were exposed to antibiotic-containing agar plates for two days at concentrations where the given strain is susceptible (2x, 4x, and 8x MIC). Three parallel lines per genotype were investigated. We failed to detect any significant increment in resistance in the mutator lines, see SN-Table 1.

**SN-Table 1**. **SCH-79797 MIC values of AB-SEN and mutator *A.baumannii* lines derived from FoR assay.**

| | SCH-79797 MIC (µg/ml) | | | |
|---|---|---|---|---|
| Genotype | Ancestral | Replicate 1 | Replicate 2 | Replicate 3 |
| *A.baumannii* ATCC 17978 (AB-SEN) | 4 | 8 | 8 | 8 |
| *A.baumannii* ATCC 17978 mutator | 8 | 8 | 8 | 8 |

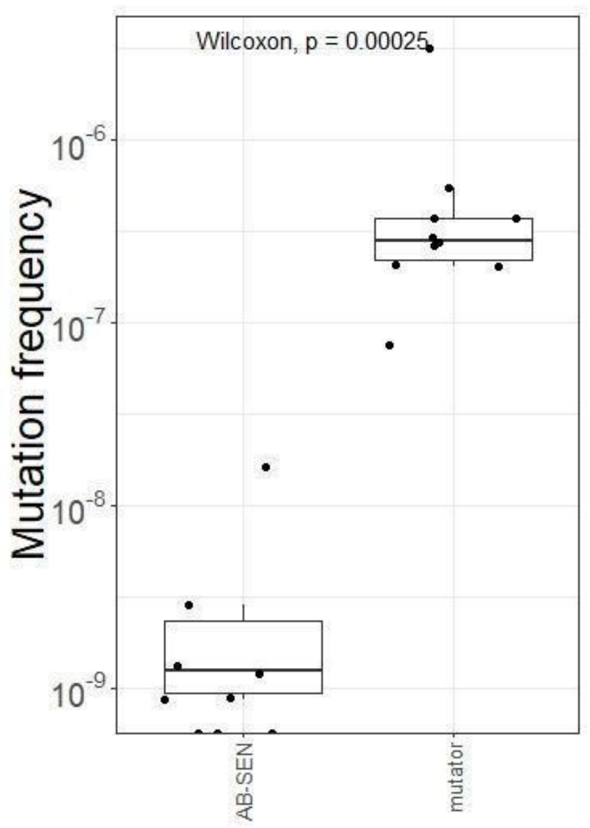


**SN-Fig. 1. Mutation frequency of wild-type and mutator *A. baumannii* strains.** Mutation frequency was assessed using a standard rifampicin assay. The mutation frequency *A. baumannii* ATCC 17978 (AB-SEN) is approximately 250-fold lower compared to the corresponding mutator strain (Wilcoxon rank-sum test, P= 0.00025). Boxplots display the median, first, and third quartiles, with whiskers indicating the 5th and 95th percentiles. Each data point represents one of the 10 parallel replicates. *Method:* The mutation frequency was measured with rifampicin assay[17]. Briefly, a single colony was selected and cultured in 10 ml of LB broth, incubated overnight at 37°C. Subsequently, 5 µl of this overnight culture was transferred into fresh LB medium (50 ml) and allowed to grow for another overnight period at 37°C. Following this, 10 parallel replicates were established in 1 mL of LB, each with an initial density of 5000 cells/mL. These replicates were then incubated until reaching the stationary phase (16-18 hours). Post incubation, 1 ml from each replicate was spread on selective LB agar plate (100 µg/ml rifampicin), and colony-forming units (CFUs) were calculated on non-selective LB agar plates. After 48h at 37°C, mutation frequency was calculated by dividing rifampicin-resistant colonies by the CFU on non-selective plates.

**Supplementary Note 3.**

**Shortage of mutated genes during antibiotic selection involved in adaptation to the laboratory medium**

An important open issue is the extent to which adaptation during the course of laboratory evolution was driven by the physical conditions, including the medium unrelated to antibiotic selection. Several independent lines of evidence indicate that antibiotic selection was the dominant force in our experiments.

*Bioinformatics analysis of mutations promoting growth in the laboratory.* To investigate this issue, we first used data from prior studies[18,19] devoted to exploring the mutations enabling rapid growth of *Escherichia coli* in specific laboratory conditions (in the absence of antibiotic stress). The experimental settings for the laboratory evolution used in these studies were comparable to those employed by us: batch cultures using synthetic defined medium (M9 and MHB) with glucose as a sole carbon source. We found that the overlap in the mutated gene sets was minimal: only 5% of the 218 *E. coli* genes mutated in our screen have been previously implicated in adaptation to laboratory conditions. When the analysis was restricted to the MHB medium used in our screens, the overlap declined further to a mere 1%. In addition, laboratory-observed mutations associated with medium-adaptation were present at a lower frequency in natural strains than putative resistance mutations found in our screens.

*Laboratory evolution in multiple bacterial species.* To expand the above analysis, we initiated laboratory evolution experiments intending to identify mutations that provide adaptation to the physical conditions in all four bacterial species. The same experimental settings, protocols, and 8 bacterial strains/species were employed, but no antibiotic was added to the medium. Parallel lines were serially passed in an antibiotic-free MHB medium and then subjected to whole genome sequencing, using the same experimental and bioinformatics protocols as earlier (Methods). For simplicity, we term these lines as control-adapted lines (n=40).

Two control-adapted lines (1 in *E. coli* and 1 in *K. pneumoniae*) accumulated exceptionally large numbers of mutations (83 and 115 in *E. coli* and *K. pneumoniae*, respectively), possibly due to becoming hypermutators during laboratory evolution. Therefore, they were excluded from further analysis. In the remaining lines, 198 unique mutational events, including 180 single nucleotide polymorphisms (SNPs) and 18 insertions or deletions (indels) were detected. In general, we found a significant excess of non-synonymous over synonymous mutations, indicating that the accumulation of the SNPs in protein-coding regions was largely driven by selection towards adaptation to the laboratory medium (SN-Fig. 2.).

Next, we sought to assess the extent of overlap of molecular mechanisms underlying adaptation in control-adapted and antibiotic-adapted lines. The analyses focused on non-synonymous mutations and short frame-shift only, as these mutational events are especially likely to change the functionality of the corresponding gene.

When mutated amino acid sites were considered, no overlap was found between control-adapted and antibiotic-adapted lines. Therefore, we focused on the overlap of mutated genes between the two groups and found that the overlap is negligible. In total, 99 and 487 mutated protein-coding genes were detected in control and antibiotic-adapted lines (ALE), respectively. However, altogether only 5.8 % of these genes were mutated in both groups. These results indicate that the gene sets mutated during antibiotic adaptation differ from those conferring a growth advantage in the specific medium used (SN-Table 2, SN-Fig. 3.). Of note, even this small set of overlapping genes has clear roles in antibiotic selection, and should not be discarded. For example, the *acrB* gene is part of the AcrAB-TolC efflux pump complex in Gram-negative bacteria and this efflux pump is responsible for pumping out a wide range of antibiotics from the bacterial cell, reducing the intracellular concentration of these drugs and thereby contributing to antibiotic resistance. Similarly, the role of the *fusA* gene is evident in antibiotic resistance. It encodes the elongation factor G (EF-G), involved in protein synthesis. Alterations in the target site of antibiotics that engage in ribosomal interactions, such as aminoglycosides, can occur due to mutations in *fusA*[20,21]. Another gene identified as mutated in control lines was *lptD*, which encodes a protein integral to the assembly complex of lipopolysaccharides (LPS). Mutations in *lptD* can alter the structure or assembly process of LPS, affecting the permeability of the outer membrane to antibiotics and thus contributing to resistance, mostly against membrane-targeting antibiotics[22–24].

As further support, we compared the mutated gene set in all possible combinations of evolutionary lines started from the same genotype. On average, pairs of control-adapted evolutionary lines shared 22 % of their mutated genes. Similarly, pairs of evolutionary lines adapted to the same antibiotic shared 25.4 % of their mutated genes. This antibiotic-specificity of parallel molecular evolution cannot be explained by adaptation to the medium. By sharp contrast, there was only a minimal or no overlap in the set of mutated genes when all possible pairs of control and antibiotic-adapted lines were considered (2% on average, SN-Fig. 4.).

Together, these results indicate that adaptation during the course of laboratory evolution was largely unrelated to the environmental medium employed.

**SN-Table 2. Comparison of the number of mutated genes between antibiotic-adaptation and medium-adaptation.**

| | N mutated genes antibiotic-adapted lines | N mutated genes control-adapted lines | % overlap |
|---|---|---|---|
| *Acinetobacter baumannii* | 156 | 33 | 3.8 |
| *Escherichia coli* | 177 | 21 | 2.6 |
| *Klebsiella pneumoniae* | 166 | 32 | 3.1 |
| *Pseudomonas aeruginosa* | 110 | 25 | 3.0 |

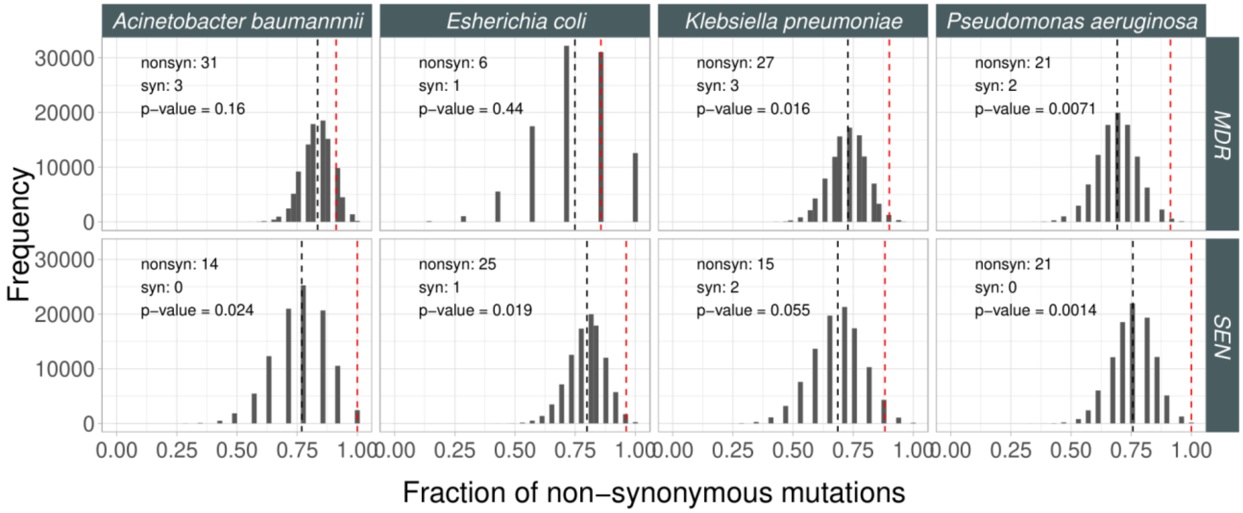


**SN-Fig. 2. Fraction of non-synonymous mutations in control adapted lines.** To assess the signatures of adaptive evolution in our genomic samples from lines adapted to growth conditions, we tested whether the fraction of non-synonymous mutational events within all SNPs in the coding region was higher than expected based on a purely neutral model of evolution using an established method (see methods). For each bacterial strain background, we identified all SNPs in the coding regions, counted the total number of non-synonymous (nonsyn) and synonymous (syn) SNPs across 5 replicate evolved lines, and calculated the observed fraction of non-synonymous as follows: nonsyn / nonsyn + syn, where nonsyn and syn are the number of non-synonymous and synonymous mutations, respectively. Next, we randomly generated the same number of SNPs at random coding positions along the genome as observed in the mutation dataset. We repeated this step 5000 times, then plotted the fraction of non-synonymous mutations as histograms for each species (columns) and strain type (rows). Next, we calculated the probability (P-value) that the fraction of non-synonymous mutations was equal to or higher in the real data than that of the randomly generated ones.

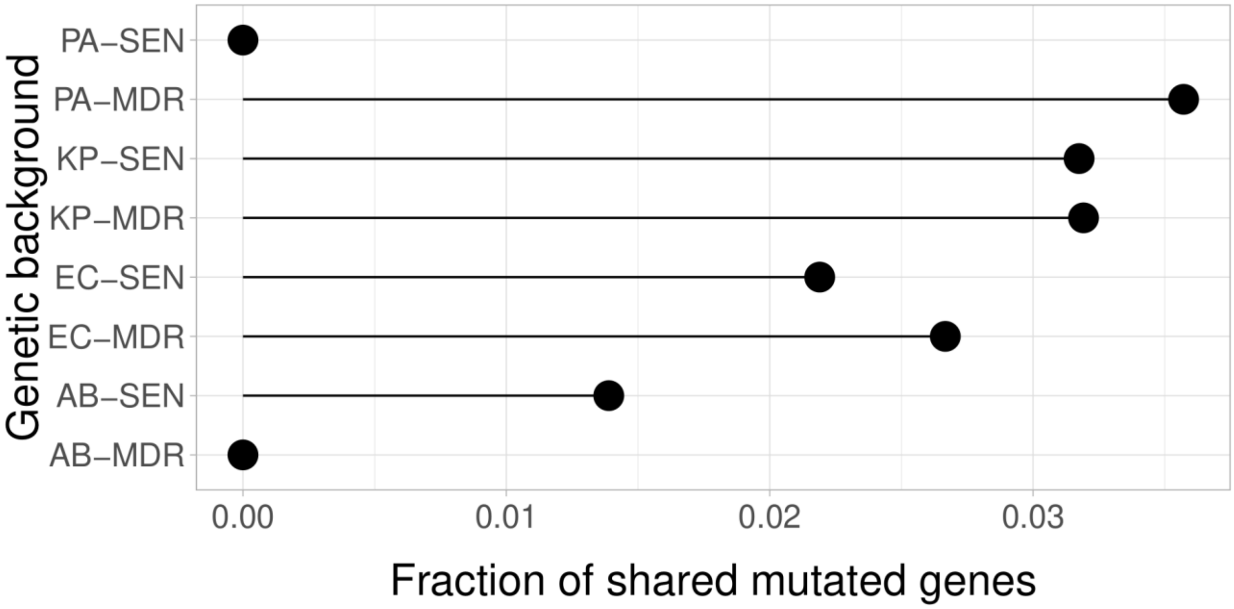


**SN-Fig. 3. The extent of overlap in the set of mutated genes between antibiotic and control-adapted lines.** The figure shows the average fraction of genes shared by all possible pairs of lines with a specific genetic background. See strain abbreviation in Supplementary Table 2.

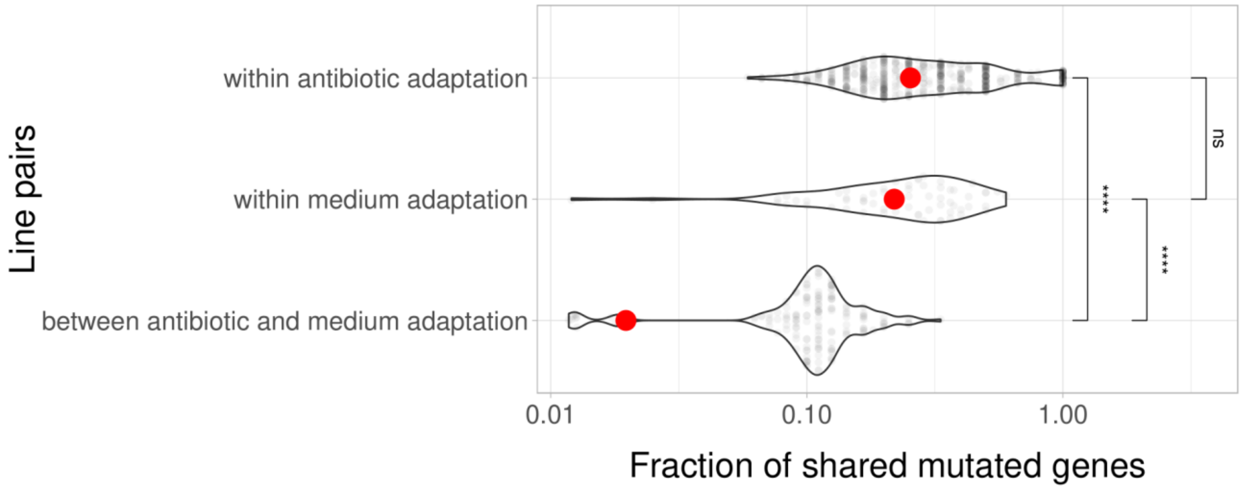


**SN-Fig. 4. The extent of overlap in the set of mutated genes.** The figure shows the distribution of mutational overlap for all possible pairs of lines founded from the same genotype across 3 different groups of pairs: i) pairs of evolved lines adapted to the same antibiotic (within antibiotic adaptation), ii) pairs of control adapted lines (within medium adaptation), and iii) pairs of control and antibiotic adapted lines (between antibiotic and medium adaptation). Red points mark the average fraction of mutated genes shared by all possible pairs of lines started from the same genotype for each group. Significance between groups was assessed by Wilcoxon's rank sum test. **** and ns indicates $P < 0.0001$ and P = not significant, respectively.

**Supplementary Note 4.**

### Non-canonical resistance mechanisms

Our analysis identified several other repeatedly mutated genes with no established roles in antibiotic resistance. Some of these genes could potentiate drug resistance evolution by shaping antibiotic tolerance or by promoting new pathways toward resistance. For example, Lon is an ATP-dependent protease responsible for the degradation of misfolded and rapidly degraded regulatory proteins. Interestingly, a prior work indicates that rapid resistance evolution in the laboratory resulted not from the lon mutant's tendency to survive in the presence of the selection drug, but rather from its capability to access new sets of mutations that enhance resistance[25]. Similarly, the ClpXP protease complexes were regularly mutated in response to distinct antibiotics. Indeed, this system ensures the integrity and proper functioning of the cell wall and membrane in times of stress, and by controlling the stability and turnover of key regulatory proteins, the ClpXP complex shapes the general stress responses in bacteria[26–29].

Prior works also suggested that the stringent response could speed up resistance evolution by increasing antibiotic tolerance. The stringent response is governed by the intracellular concentration of guanosine pentaphosphate and related molecules ((p)ppGpp), directly mediated by the RelA and SpoT enzymes[30]. These genes were regularly mutated, but only in response to specific antibiotic treatments. We found 24 evolved lines carrying mutations in the *spoT* or *relA* genes (SN-Fig 5.). 54% of the observed *spoT* mutations are frameshifts or small indels which most likely lead to reduced functionality. This aligns with the finding that SpoT inactivation yields elevated stringent response[31]. Additionally, we also identified 41 lines carrying mutations in different aminoacyl-tRNA synthetase genes (SN-Fig 6., Supplementary Table 7). This is in line with expectation, as prior studies showed links between elevated stringent response and reduced aminoacyl-tRNA synthetase activity leading to the presence of uncharged tRNAs[30,32]. Analysis of the mutations in these genes indicates that the stringent response could play a pivotal role in adaptation to topoisomerase inhibitors with new target sites or modes of action (e.g., gepotidacin, delafloxacin and zoliflodacin), the carbapenem derivative sulopenem, and specific membrane targeting antibiotics (e.g., polymyxin-B, SPR-206 and Tridecaptin-M152-P3). Interestingly, mutations in *spoT* and *relA* were notably common among *P. aeruginosa* strains, whereas aminoacyl-tRNA synthetase mutations exhibited an especially high prevalence in *E. coli* (SN-Fig 7. and SN-Fig 8.). The reasons for these species and antibiotic class-specific differences in molecular adaptation remain to be explored.

According to the Virulence Factor Database[33], 49 mutated genes in our dataset shape various aspects of bacterial virulence, including bacterial adherence to host cells, fimbriae formation, immune modulation, or biofilm formation (SN-Fig 9.). For example, laboratory-evolved SPR206 resistance was driven by mutations in the BasS/BasR two-component system (Supplementary Table 6). Remarkably, the BasS/BasR system shapes resistance to cationic peptides by regulating the production of lipopolysaccharide molecules, controlling the surface charge of the outer bacterial membrane[34,35]. Additionally, these genes play a central role in bacterial virulence in mice and are necessary for bacterial invasion of macrophages[36]. These data indicate that antibiotic resistance could simultaneously affect bacterial virulence, as a side-effect. This possibility should be explored in detail in future studies.

Regulatory mutations could also have an important accessory role in resistance to new antibiotics, for two reasons. First, mutations in specific transcription factors occurred repeatedly, and generally in an antibiotic-specific manner (SN-Fig 10.). For example, *soxR*, a regulatory gene involved in defense against oxidative stress, was regularly mutated in response to topoisomerase inhibitors (Fig ReRe19). Others directly regulate the expression of genes involved in multidrug transport, such as *acrR*, *nfxB*. Interestingly, we found that NusA, a transcription termination factor, was mutated in response to polymyxin-B treatment (SN-Fig 11.). This could be significant, as this gene was previously associated with antimicrobial peptide resistance. In addition, 66 % of the 276 detected non-codingmutations were located in putative promoter regions, as defined earlier[37] (PMID: 30980662). On average, non-coding mutations were relatively rare (SN-Fig 12A). Although the number of mutations in protein coding region surpasses the number of mutations in non-coding regions, it is important to note that in the studied bacterial genomes, the fraction of protein coding regions is larger. When normalized to the mutational target size, the important contribution of mutations in non-coding regions to antibiotic resistance becomes even more apparent (SN-Fig 12B), as suggested previously[37]. Indeed, promoter mutations occurred repeatedly in 1113 genes, some of which have established links to antibiotic resistance, bacterial virulence, or stress response (SN-Fig 13). For example, mutations in the promoter region occurred four times in MqsR in response to gepotidacin treatment. This could be significant as MqsR increases persister cell formation via toxin production[38,39]. Another example is *csuAB*, a gene involved in the formation of biofilms, surface adhesion, and antibiotic resistance[40]. Other genes, such as *mexA* [41,42], *acrE*[43,44], and *mdfA*[45] are important members of general efflux systems.

Finally, we note that mutations occurred in genes that have been considered promising targets for the future development of antibiotics with new modes of action. For example, MreB, a bacterial cytoskeleton protein, has been extensively researched for its role in determining crucial subcellular processes such as cell division, chromosome segregation, and cell wall morphogenesis[46]. As it has an essential function and is a conserved protein in most rod-shaped bacteria, it could be a promising antibiotic target of significant importance[47]. Of note, however, that MreB was mutated repeatedly in response to eravacycline and POL7306 exposures. We speculate that such mutations might interfere with the inhibitory effects of other antibiotics targeting the same gene.

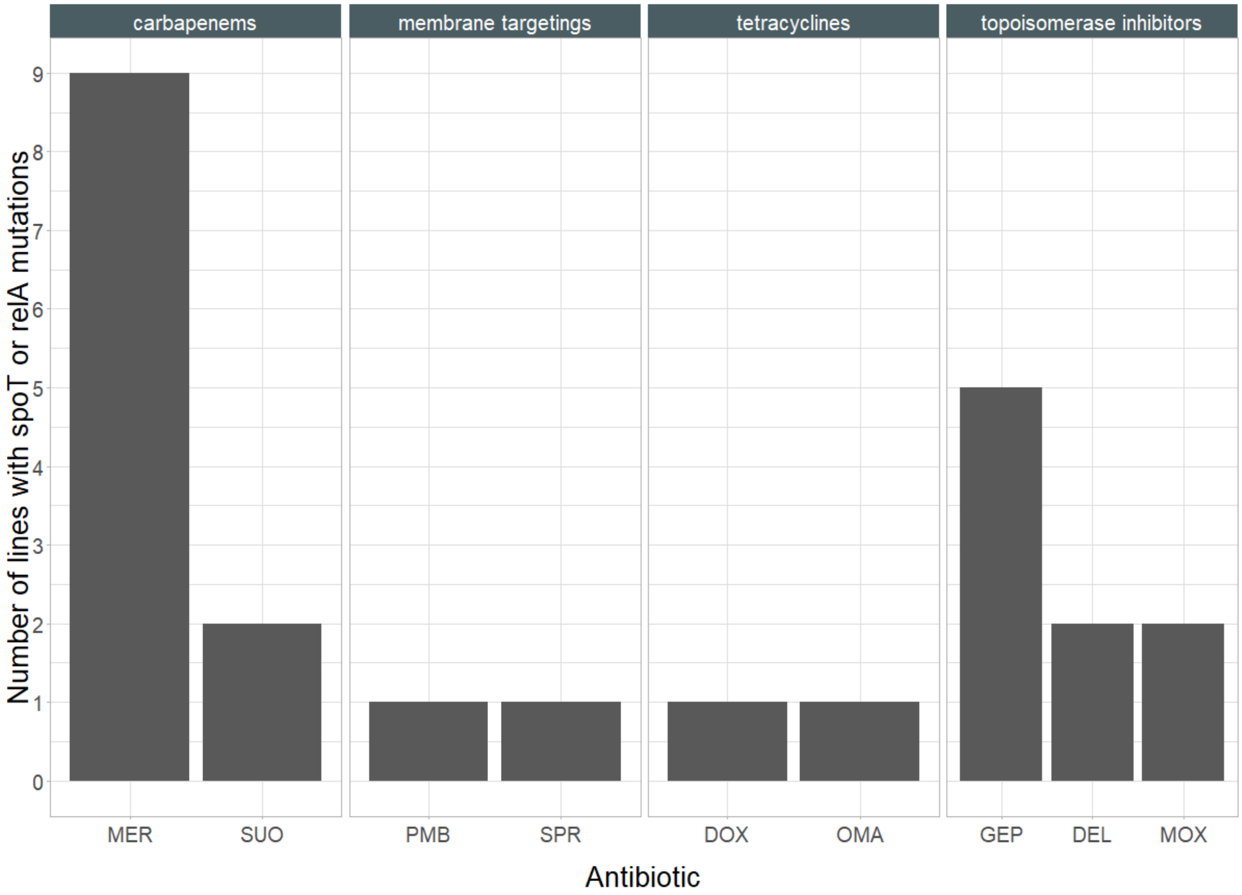


**SN-Fig 5. Distribution of the evolved lines with *spoT* or *relA* mutations.** Each bar represents the number of lines adapted to a specific antibiotic harboring mutations in the *spoT* or *relA* genes. Mutations in these genes are predominantly found in lines adapted to carbapenems, while they occur only occasionally in membrane targeting and tetracycline-adapted lines. No lines adapted to aminoglycoside and cephalosporin carried mutations in these genes. For antibiotic abbreviations, see Table 1.

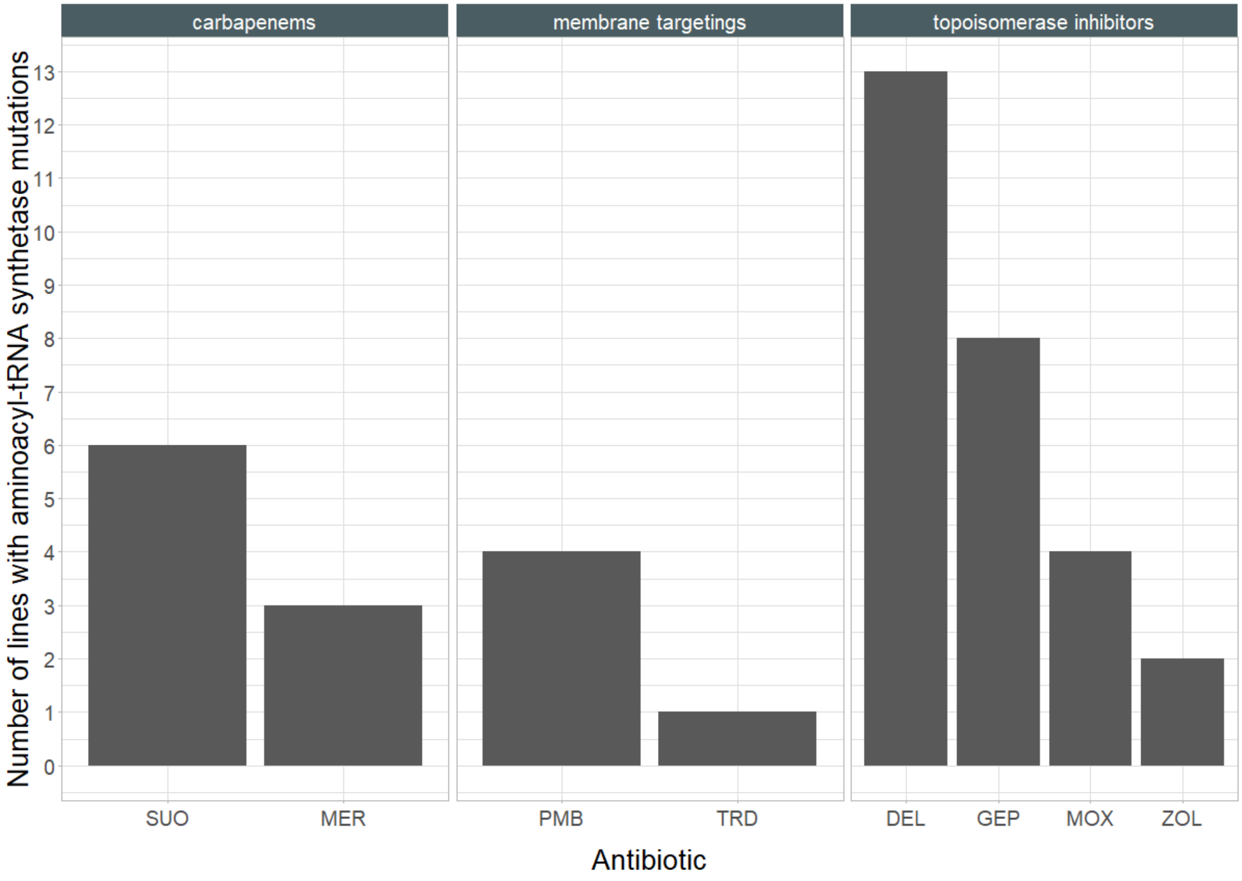


**SN-Fig 6. Distribution of the evolved lines with aminoacyl-tRNA synthetase gene mutations.** Each bar represents the number of lines adapted to a specific antibiotic harboring aminoacyl-tRNA synthetase gene mutations. Mutations in these genes were predominantly found in carbapenem, membrane targeting, and topoisomerase-adapted lines. For antibiotic abbreviations, see Table 1.

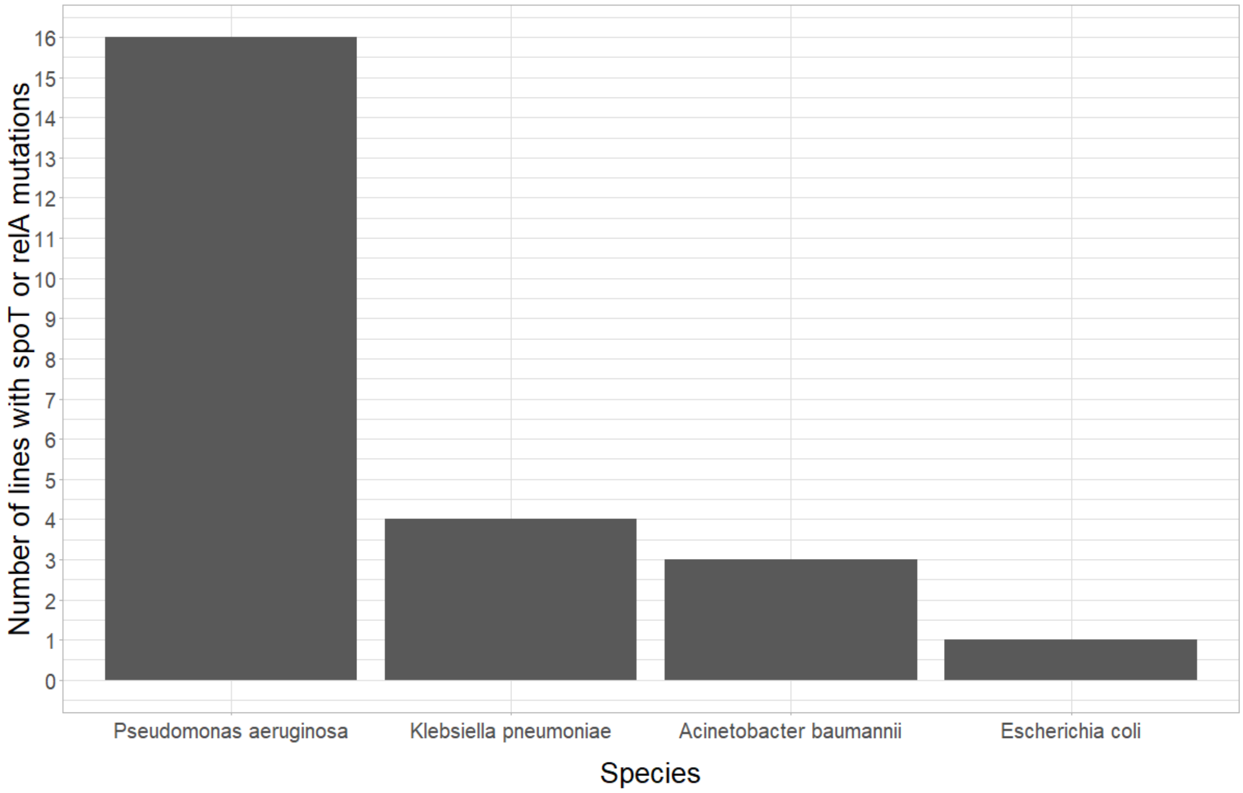


**SN-Fig 7. Mutations in *spoT* and *relA* in four bacterial species.** The figure illustrates the distribution of evolved lines harboring mutations in the *spoT* or *relA* genes across bacterial species. Columns represent the number of lines carrying either spoT or relA mutations. Mutations in *spoT* and *relA* exhibit a significantly higher frequency in *P. aeruginosa* strains, as evidenced by Fisher's exact test ($P < 0.0001$).

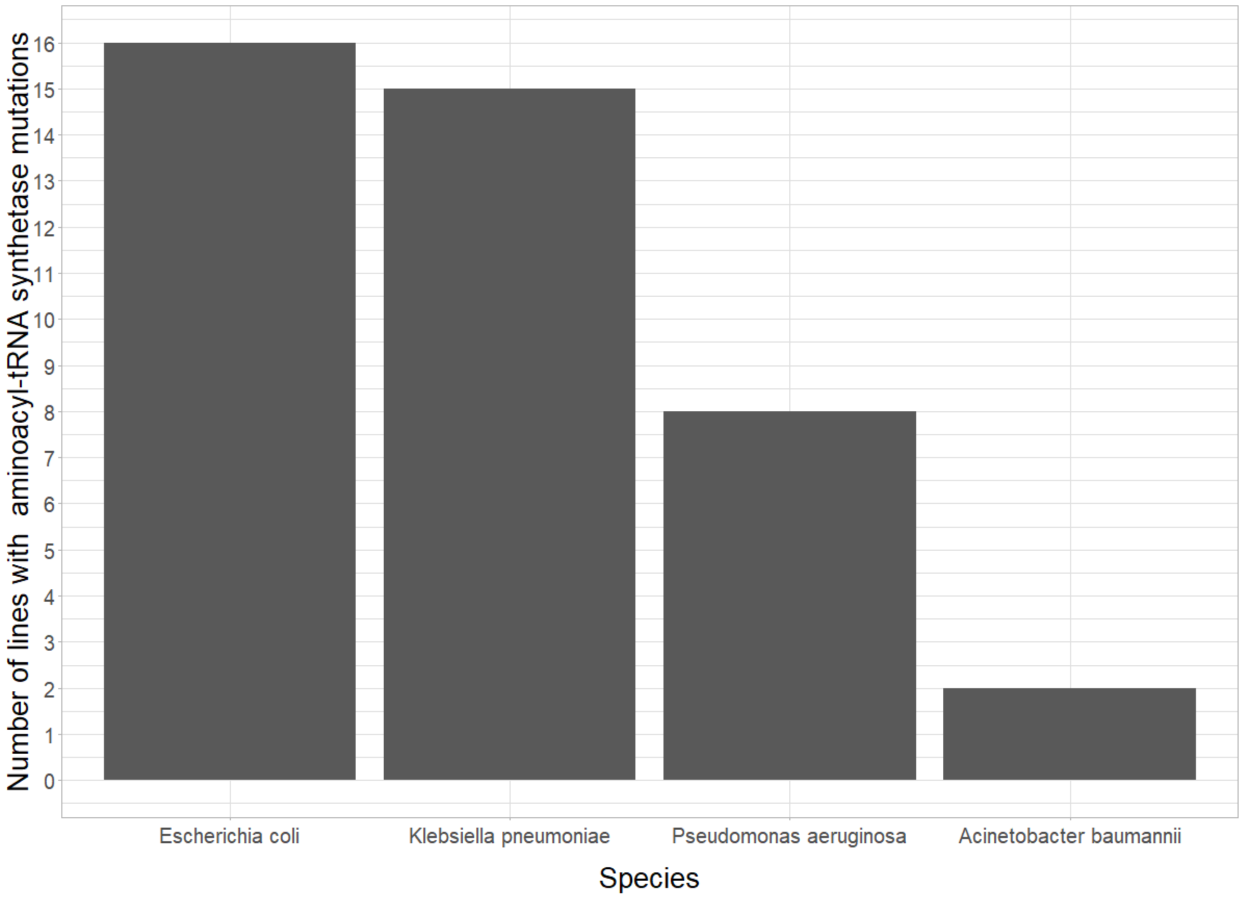


**SN-Fig 8. Mutations in aminoacyl-tRNA synthetase genes in four bacterial species.** The figure illustrates the distribution of the evolved lines harboring mutations in aminoacyl-tRNA synthetase genes across bacterial species. Mutations in aminoacyl-tRNA synthetase genes were especially frequent in *E.coli* (Fisher's exact test, P-value = 0.03).

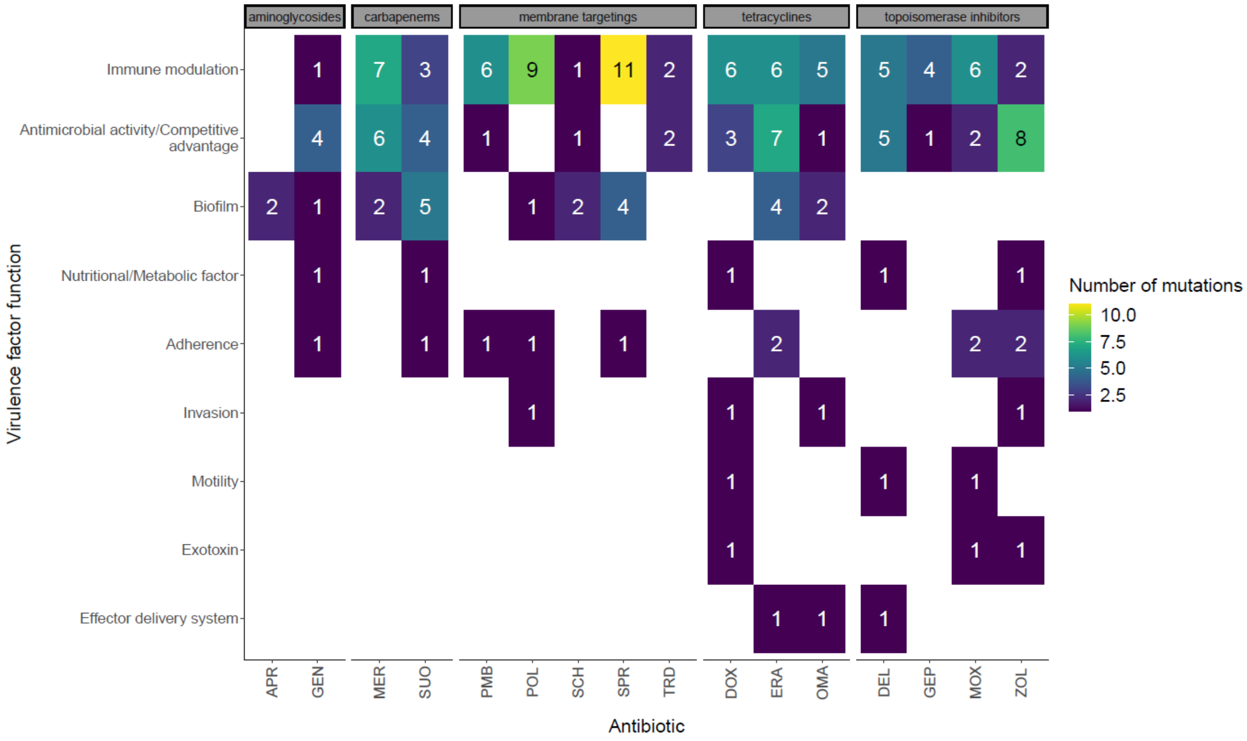


**SN-Fig 9. Virulence factor functions of the genes mutated in response to different antibiotics.** Unique mutated genes were assigned to functional categories based on information obtained from the Virulence Factor Database (VFDB). The figure shows the total number of unique mutations associated with each virulence factor function category per antibiotic treatment. For antibiotic abbreviations, see Table 1.

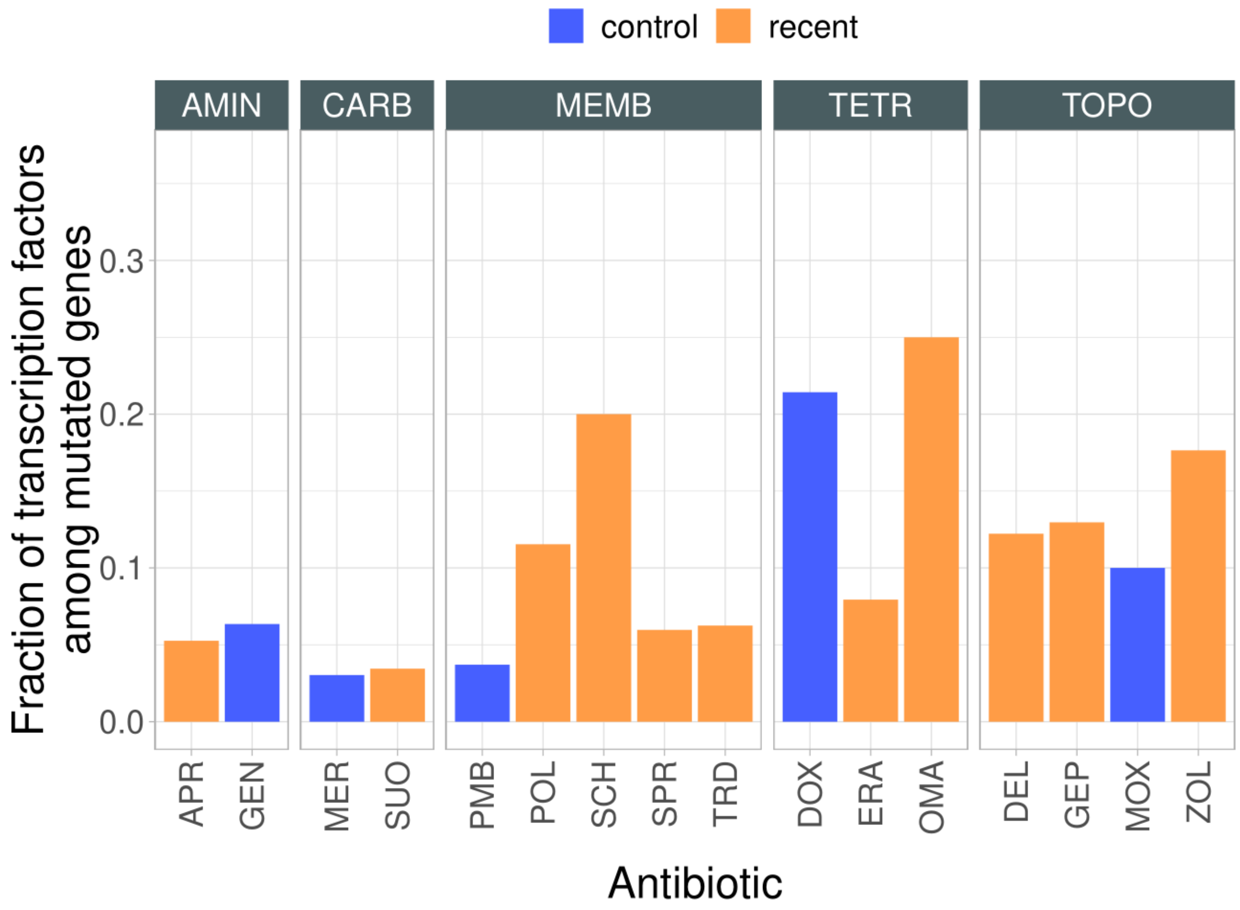


**SN-Fig 10. Fraction of mutated transcription factors.** The figure shows the fraction of transcription factors among mutated genes across lineages adapted to different antibiotics (adaptive laboratory evolution (ALE)). Identification of transcription factors was based on Gene Ontology (GO) term associated with DNA-binding transcription factor activity, using protein sequence homology. There is significant variation in the frequency of mutated transcription factors across the antibiotics tested (Proportion test, $p < 0.05$), indicating that adaptations to certain antibiotics are more likely to result in transcription factor mutations than others. Transcription factors were especially likely to mutate in response to doxycycline (DOX) and omadacyclin (OMA) treatments. Abbreviations: TOPO: topoisomerase inhibitors, TETR: tetracyclines, AMIN: aminoglycosides, CARB: carbapenems, CEPH: cephalosporins, MEMB: membrane targeting antibiotics. For antibiotic abbreviations, see Table 1.

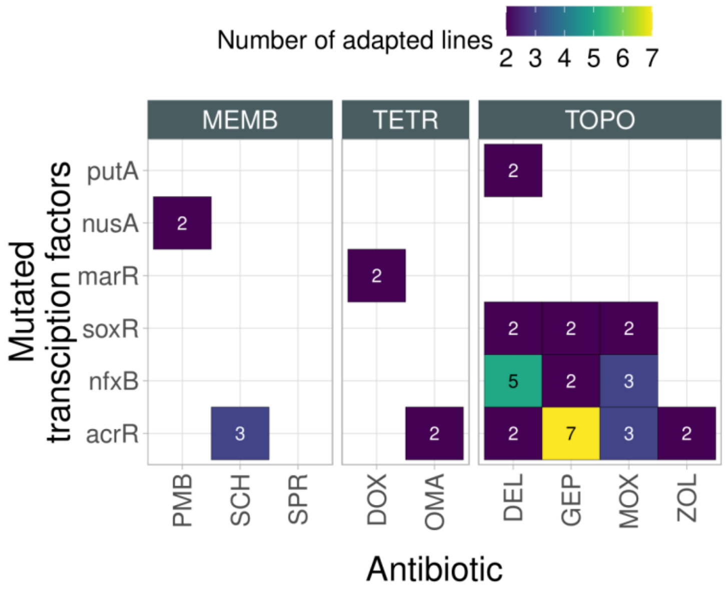


**SN-Fig 11. Transcription factors mutated across antibiotic treatments.** The heatmap shows the number of adapted lines with mutated transcription factors in lines derived from adaptive laboratory evolution (ALE). Only transcription factors mutated in at least two independent adapted lines are shown. Abbreviations: MEMB: membrane targeting antibiotics, TETR: tetracyclines, TOPO: topoisomerase inhibitors. For antibiotic abbreviations, see Table 1.

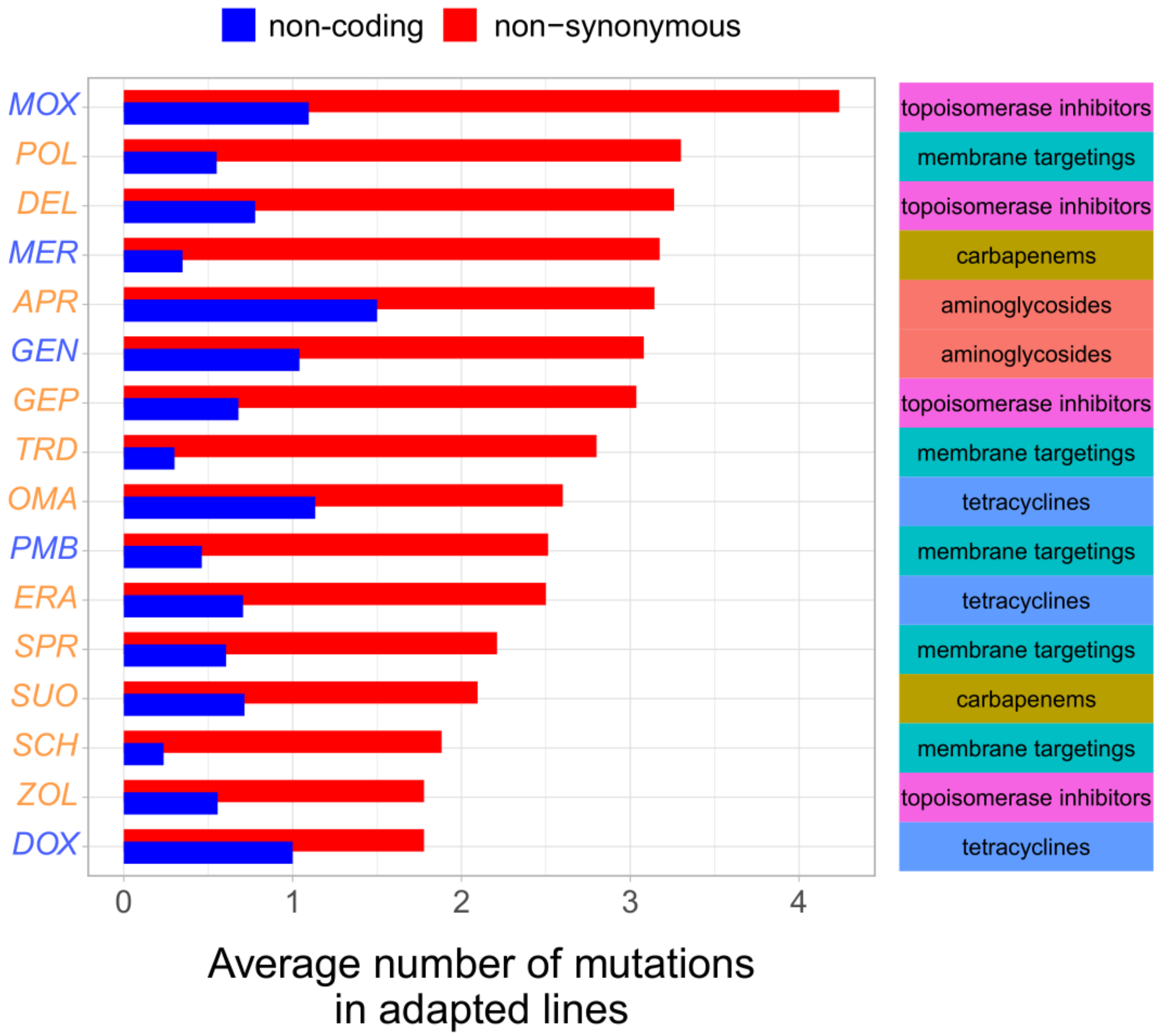


**SN-Fig 12A. Non-coding mutations in laboratory evolved lines.** The figure shows the average number of non-synonymous mutations in protein-coding regions (red) and non-coding mutations (blue), respectively, across evolved lines adapted to different antibiotics (orange and blue labels mark recent and control antibiotics, respectively). 66% of the detected 276 non-coding mutations were located in putative promoter regions, as defined earlier[37]. The heatmap on the right panel depicts the antibiotic class a given antibiotic belongs to. For antibiotic abbreviations, see Table 1.

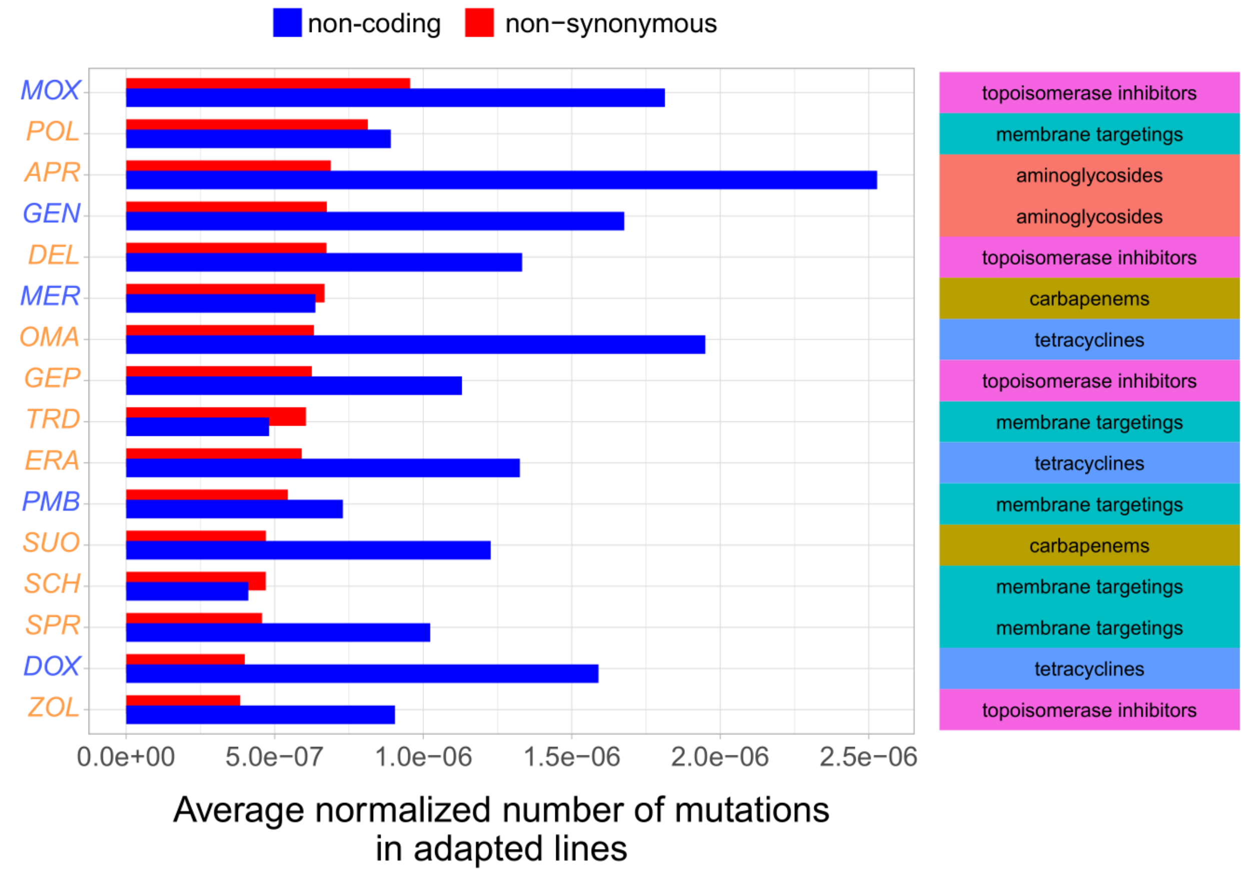


**SN-Fig 12B. Non-coding mutations in laboratory evolved lines.** The figure shows the normalized number of non-synonymous mutations in protein-coding regions (red) and mutations in non-coding regions (blue), respectively, across evolved lines adapted to different antibiotics (orange and blue labels mark recent and control antibiotics, respectively). Normalized number of mutations was by estimated by calculating the number of mutations per line, divided with the mutational target size (i.e. total size of the corresponding genomic region type). The heatmap on the right panel depicts the antibiotic class a given antibiotic belongs to. For antibiotic abbreviations, see Table 1.

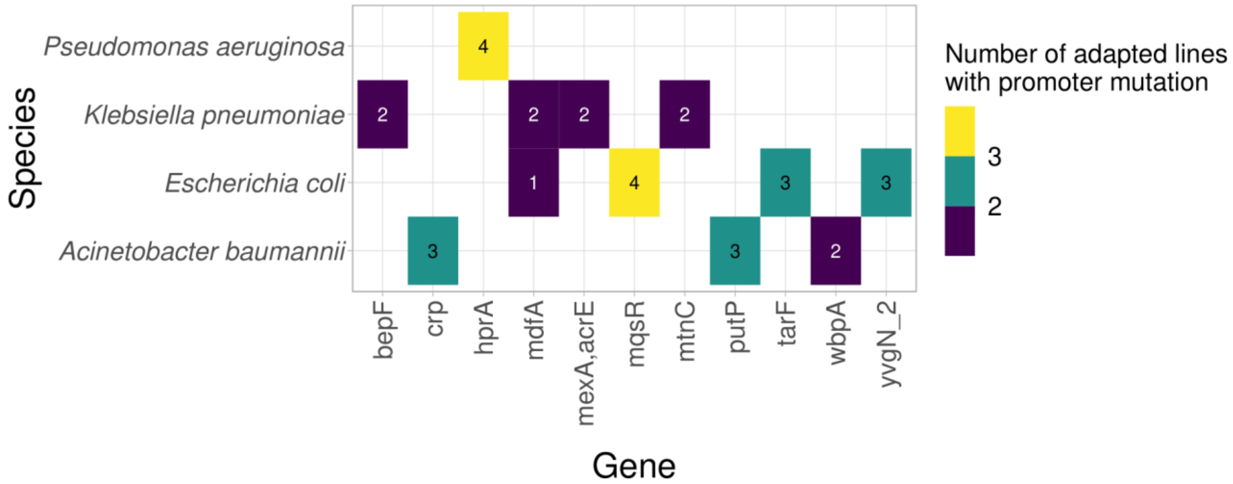


**SN-Fig 13. Genes repeatedly mutated in their promoter regions.** The heatmap shows the number of adapted lines with promoter mutations in the corresponding genes across different bacterial species.

**Supplementary Tables (in Auxiliary Supplementary File):**

1. Information about antibiotics: Supplementary Table 1 – Table_S01.xlsx
2. Information about bacterial strains: Supplementary Table 2 - Table_S02.xlsx
3. Data of the high throughput MIC measurement: Supplementary Table 3 - Table_S03.xlsx
4. Antibiotic Resistance Genes previously described in the literature and Claims of Superiority of recent antibiotics: Supplementary Table 4 - Table_S04.xlsx
5. Relative MIC following FoR assay and ALE: Supplementary Table 5 - Table_S05.xlsx
6. Detailed description of the mutations detected in adapted strains: Supplementary Table 6 - Table_S06.xlsx
7. Commonly mutated genes (at least 2 or 3 times per species), mutations of putative MDR genes and efflux activity: Supplementary Table 7 - Table_S07.xlsx
8. Description of identified MDR genes: Supplementary Table 8 - Table_S08.xlsx
9. Non-synonymous mutations found in natural strains of *E. coli* and *A. baumannii*: Supplementary Table 9 - Table_S09.xlsx
10. DIvERGE related data: Supplementary Table 10 - Table_S10.xlsx
11. Description of functional metagenomic libraries: Supplementary Table 11 - Table_S11.xlsx
12. Functional metagenomic sequencing results (DNA contigs, ORFs) and MIC data: Supplementary Table 12 - Table_S12.xlsx

**Supplementary Data:**

1: Metadata for laboratory-observed mutations found in natural strains: Data S1 – Data_S01.xlsb
